# The biosynthetic-secretory pathway, supplemented by recycling routes, specifies epithelial membrane polarity

**DOI:** 10.1101/2022.01.06.475118

**Authors:** Nan Zhang, Hongjie Zhang, Liakot A. Khan, Gholamali Jafari, Yong Eun, Edward Membreno, Verena Gobel

## Abstract

In prevailing epithelial polarity models, membrane-based polarity cues (e.g., the partitioning-defective PARs) position apicobasal cellular membrane domains. Intracellular vesicular trafficking expands these domains by sorting apicobasal cargo towards them. How the polarity cues are polarized and how sorting confers long-range vesicle directionality is still unclear. Here, a systems-based approach using two-tiered *C. elegans* genomics-genetics screens identifies trafficking molecules that are not implicated in apical sorting yet polarize apical membrane and PAR complex components. Live tracking of polarized membrane biogenesis suggests that the biosynthetic-secretory pathway, linked to recycling routes, is asymmetrically oriented towards the apical domain during its biosynthesis, upstream of PARs and independent of polarized target domains. This mode of membrane polarization could offer solutions to questions of current models of polarity and polarized trafficking.

**One-Sentence Summary:** Biosynthetic trafficking polarizes epithelial membranes by asymmetrically expanding the apical domain into the growing membrane.

## Main text

Current epithelial polarity models order the highly-conserved extra-, intracellular- and membrane-based polarity cues sequentially during the process of cellular polarization: (1) extracellular matrix signals orient cells within the epithelium; (2) plasma-membrane- and junction-based apical and basolateral core polarity determinants (the partitioning-defective PAR-, Crumbs- and Scribble polarity complexes) specify apicobasal domain positions at the plasma membrane; (3) intracellular vesicular trafficking expands and maintains these domains by supplying polarized membrane components (*1–3*). This hierarchy of events is thought to control both the initial polarization of the cell membrane during polarized tissue morphogenesis and the initial asymmetric delivery of membrane components during polarized membrane biogenesis itself. Such a polarity model is consistent with the current model of polarized trafficking where vesicles sort apicobasal cargo to cognate recognition sites at target plasma membrane domains, a process that requires these domains’ prior polarization (*1, 4–6*). Unresolved difficulties of these models include the question how the membrane-based polarity cues themselves are polarized to domains whose positions they ought to specify; and how the sorting of polarized cargo confers long-range apicobasal directionality to vesicles.

We previously demonstrated that glycosphingolipids/GSLs, saturated obligate membrane lipids, function in apicobasal membrane polarity in the *C. elegans* intestine. Reducing and restoring GSL biosynthesis reversibly shifts apical membrane components, -junctions, -PARs, and the lumen between apical and basolateral sides of growing cells that no longer divide and move but still expand their plasma membranes (*in vivo* apicobasal polarity conversion/reversion) (*7*). GSLs, components of endo- and plasma membrane microdomains (rafts), have conserved roles in apical sorting on Golgi- and post-Golgi vesicle membranes (4), suggesting that polarity conversion arose from a trafficking defect involving apicobasal sorting. However, the ability of trafficking to generate junction-bounded apical domains on basolateral membranes and thus override pre-existing membrane-based polarity cues also raised the possibility that trafficking might directly polarize the membrane while synthesizing it, e.g., on biosynthetic routes whose directionality was regulated independent of previously polarized target membrane domains. Early epithelial polarity models had envisioned a similar scenario, where the fixed (apical or basolateral) directionality of bulk membrane delivery directly polarizes the growing membrane (*8, 9*).

In support of such an upstream function of trafficking in membrane polarity, we and others subsequently identified components of the conserved post-Golgi vesicle coat clathrin and its AP-1 adaptor by a similar loss-of-function polarity conversion phenotype in the *C. elegans* intestine (*10, 11*). However, GSLs have a wide range of trafficking-dependent and independent functions and clathrin and AP-1, the former best known for its role in endocytosis, also operate on multiple trafficking routes (*12, 13*). These molecules could therefore alter membrane polarity by other mechanisms: independent of trafficking; by canonical sorting; on other than biosynthetic trafficking routes. They might, for instance, sort on endocytic-recycling routes that reposition previously delivered, rather than position newly synthesized, membrane components and thus maintain or rearrange the polarity of the membrane rather than polarizing it during its biosynthesis. In favor of this latter possibility, various endocytic- and transcytotic-recycling molecules have been identified *in vitro* and *in vivo* that recruit apical PARs to maintain apical/anterior polarity in different cell types and species (*14–18*) and to position the apical domain in tubular epithelia (*19–26*). In contrast, no unambiguous biosynthetic-secretory molecule were identified in multiple screens on the establishment and maintenance of apicobasal polarity, despite these molecules’ expected contribution to polarity via apicobasal sorting (*18*).

To investigate the role of trafficking in membrane polarity, we here use the setting of *in vivo* apicobasal polarity conversion in the *C. elegans* intestine and employ a systems-based approach, taking advantage of the high molecular and functional conservation of vesicular trafficking components. First, we collected all trafficking molecules identified in several genome-scale multicellular (intestinal) and unicellular (excretory canal) *C. elegans* tubulogenesis screens by their function in apical membrane biogenesis. Next, we examined these molecules’ ability to modify the GSL-dependent apicobasal polarity conversion. This approach identified unequivocal biosynthetic-secretory (secretory) pathway components among enhancers of GSLs’ polarity function. The analysis of their function in wild-type polarized membrane biogenesis revealed that SEC-23/SEC-24 and multiple (9) additional secretory carrier molecules, most implicated in pre-Golgi trafficking and not in apical sorting, are independently required to specify the polarized distribution of apical membrane and PAR polarity complex components on the expanding membrane, whether or not this membrane has already been polarized. Conversely, GSL suppressors identified components of polarized endocytic-recycling routes, not required to specify membrane polarity, but supplementing this biosynthetic route and revealing new functions for the small GTPase RAB-7, DAB-1/Disabled, and the V-ATPase subunit VHA-6. A secretory pathway, linked to recycling routes, whose directionality is regulated upstream of membrane-based polarity cues during *de novo* polarized membrane biogenesis, could address open questions of current models of epithelial polarity and polarized trafficking.

### An approach to track polarized membrane biogenesis in single cells *in vivo*

The analysis of polarized membrane biogenesis under both *in vitro* and *in* and *ex vivo* conditions has been complicated by tissue complexity and by flatness of epithelial cells, as well as by confounding effects of polarized cell migration and division that occur simultaneously with polarized membrane biogenesis. To overcome these limitations, we here examine polarized membrane biogenesis live in single postmitotic cells at their invariant positions within the 3D (apicobasolateral) context of single-layered, expanding *C. elegans* tubular organ epithelia (Fig. 1A). We track ERM-1, the ancestral ortholog of the ERM/ezrin-radixin-moesin family of membrane-actin linkers that are enriched at the lumen of tubular epithelia where they denote apical membrane identity and thus successful apical membrane polarization (Fig. 1A legend; (*27–29*)). In the 20-cell *C. elegans* intestine and the 1-cell excretory canal, ERM-1 delineates the coincident events of *de novo* apical membrane and lumen biogenesis from the early embryo (initial membrane polarization and expansion; cells divide and move), through late embryogenesis and 4 larval stages (ongoing membrane polarization and expansion; cells no longer divide and move), to the adult (membrane polarity maintenance; no/minimal membrane expansion; Fig.S1) (*30, 31*). Polarized membrane biogenesis is separable from polarized cell division and migration in late-embryonic/larval tubes, where net membrane expansion can be monitored *in situ* in single postmitotic cells over ∼48 hours required for full tube extension in animals growing at room temperature (longer if growth is delayed; a ∼34- and 2000-fold increase in apical membrane length in the intestine and canal, respectively; Fig.S1B, C). ERM-1::GFP+ membranes are tracked along apicobasal cellular axes between cells during inTERcellular lumenogenesis in the intestine and along anterior-posterior cellular axes during inTRAcellular lumenogenesis inside the single canal cell. This approach permits the distinction of: (1) apical membrane expansion (*de novo* apical membrane biogenesis; extension of the ERM-1+ membrane) and (2) apical membrane polarization (acquisition of apical membrane character; ERM-1 recruitment to the membrane) from (3) apical membrane positioning (regulation of apicobasal membrane polarity; apicobasal ERM-1 distribution along the membrane).

**Figure 1.**
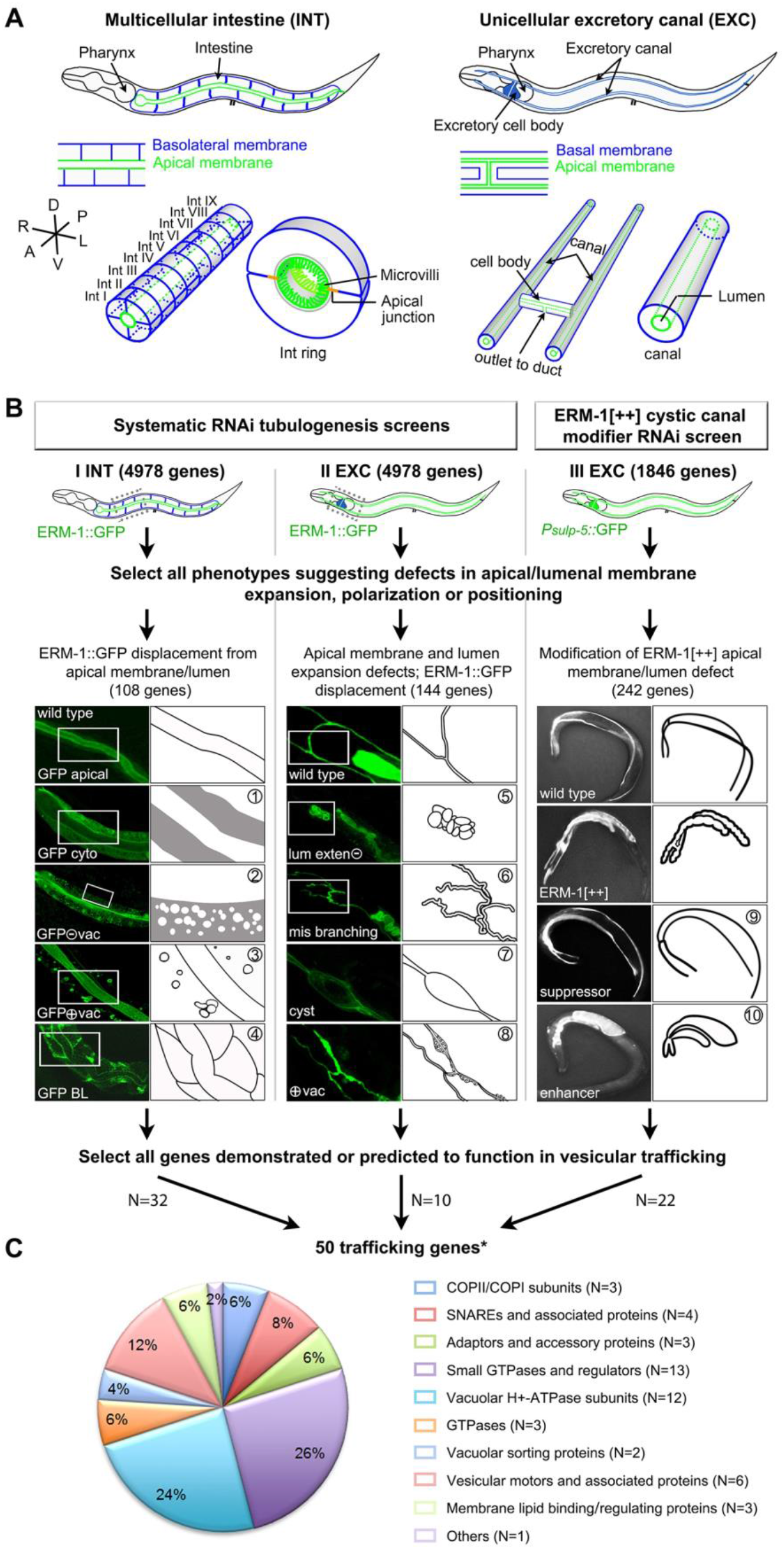
*C. elegans* intestinal and excretory canal tubulogenesis screens identify 50 trafficking molecules for inTRA- and inTERcellular lumenogenesis. **(A)** Schematics of the *C. elegans* intestine (INT, here and below), and excretory canal (EXC, here and below). INT (left): 20 cells form 9 rings in bilateral symmetry, 4 cells in ring 1. EXC (right): 2 anterior and 2 posterior canal arms extend from one single cell. Top: whole animal; middle: tube portion at higher magnification; bottom: 3D view. Here and below: apical = lumenal membrane green, basolateral membrane blue, A/anterior left, P/posterior right; D/dorsal up, V/ventral down; R/right left, L/left right. **Fig.S1** for tubulogenesis process. **(B)** Screen layouts and phenotype classes. **INT screen** (I, left column): ERM-1::GFP displacement to: the cytoplasm, homogenously (GFP cyto); a vacuolized cytoplasm (GFP○- vac); cytoplasmic vacuoles (GFP○+vac); the basolateral membrane (GFP BL) (①-④). **EXC screen** (II, middle column): no lumen extension (lum exten○-); branch loss/excess (mis-branching); cystic lumen (cyst); cytoplasmic GFP+ puncta/vacuoles (○+vac) (⑤-⑧). **EXC modifier screen** of ERM-1-overexpression/ERM-1[++]-induced short cystic canals (III, right column): suppressor = canal re-extension, enhancer = cyst enlargement and further canal shortening, in ERM-1[++] (⑨,⑩). Screens I and II are unbiased (>90% genome coverage expected, based on pilot screen), screen III targets ERM-1 interacting genes (see Methods for screen design). Confocal images left, corresponding schemata right (boxed areas of schematics or whole animals are shown). *10 genes identified in 2 screens, 2 genes in 3 screens. Note that previously characterized trafficking genes are not included (see text) (*7, 11*). **(C)** Pie chart: percentage distribution of functional classes among the identified trafficking genes. Same color coding as in **Figs.2, 3, S2**.

### Multi- and unicellular *C. elegans* tubulogenesis screens identify 50 trafficking molecules required for apical domain polarization, positioning, or expansion

From a broad spectrum of lumenogenesis phenotypes identified in genome-scale RNAi-based intestinal and excretory canal tubulogenesis screens (tier-1 screens, I-III, Fig. 1B; Methods for screen design), all phenotypes affecting apical membrane polarization, positioning or expansion were collected and sorted into 10 classes (Fig. 1B). Most intestinal phenotypes involved defects in apical membrane polarization (cytoplasmic ERM-1 displacement, with or without ERM-1+/- vacuoles; Fig. 1B, classes 1-3; N=90). As expected, *bona fide* apicobasal polarity defects (basolateral ERM-1 mislocalization; Fig. 1B, class 4), narrowly defined as such to separate defects in the regulation of membrane polarity from other aspects of membrane polarization, were only rarely encountered (N=18).

We selected all genes with demonstrated or predicted roles in vesicular trafficking from a total of 453 genes that were isolated by their functions in any of these aspects of apical membrane biogenesis. 50 such genes were identified, all encoding proteins highly conserved among species by structure, endomembrane/vesicle location, and trafficking function (corresponding to ∼3% of annotated *C. elegans* trafficking genes; Wormbase release WS267; Fig. 1C, Tab.S1; Methods; genes encoding GSL-biosynthetic enzymes and clathrin/AP-1 subunits, previously identified in these screens (*7, 11*), are not included here). 26/50 gene products belong to functional classes with members implicated in directional trafficking and polarized sorting in diverse species, e.g.: Rab GTPases/Rab-related (N=13, the largest subgroup; RAB-5, 8, 11 with documented functions in apical domain positioning) (*19, 32, 33*); motors (N=6; KLP-16/KIFC3 with documented functions in apical transport) (*34*); v-/t-SNAREs (N=4); coat adaptors (N=3; Tab.S1, Figs.1C, 2A and legends). The second largest subgroup consists of 12 subunits of V-ATPases, endo- and plasma-membrane-based proton pumps, (*35*) recently also implicated in apical secretion and polarized tubulogenesis (*36–38*). Validating screen specificity and consistent with genome coverage, multiple subunits of protein complexes and of proteins previously shown to interact were independently identified by similar phenotypes (Tab.S1, Fig. 2A legend).

**Figure 2.**
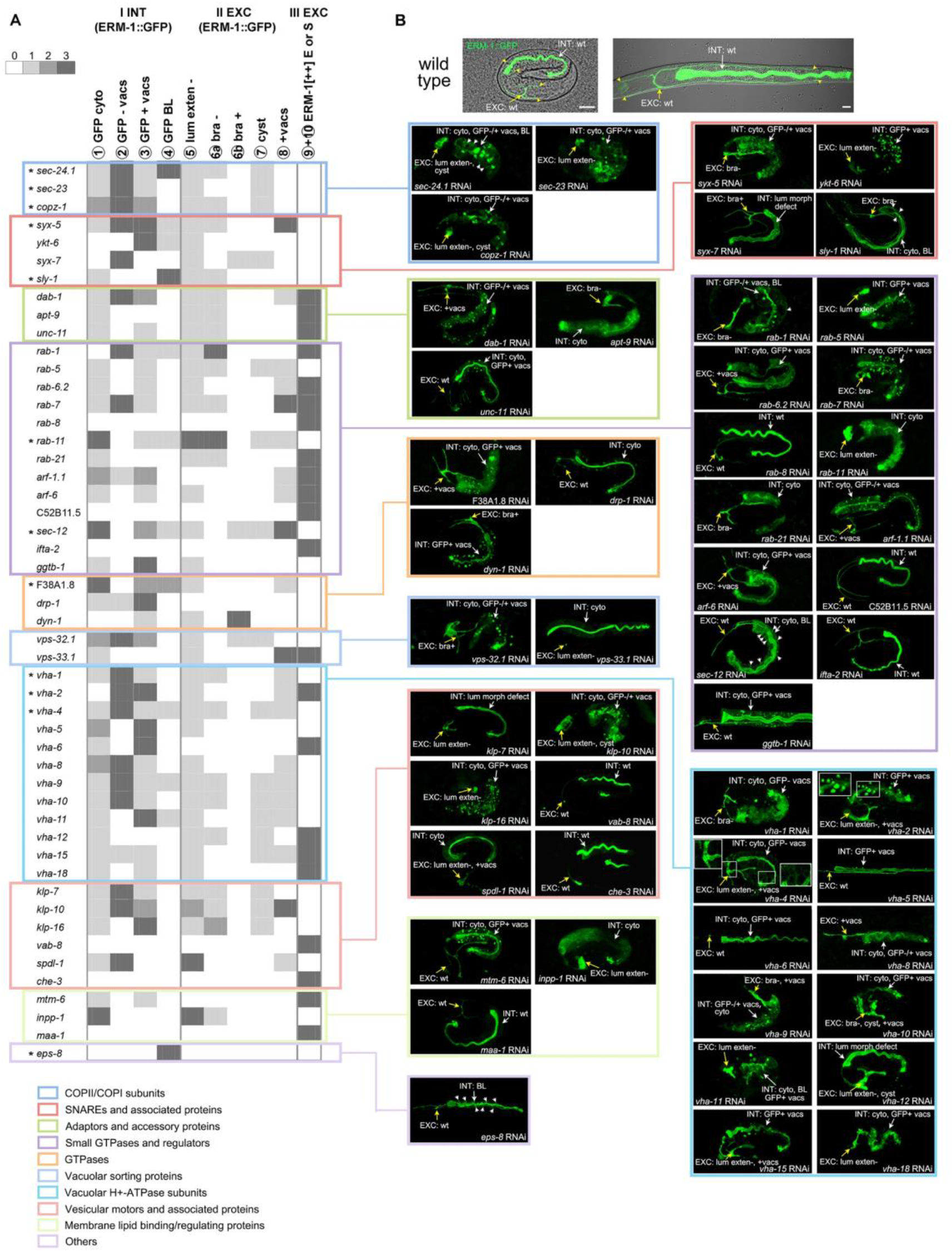
A shared set of trafficking molecules directs apical membrane polarization, positioning and expansion inside the single canal cell and between pairs of intestinal cells. **(A)** 50 trafficking genes identified in INT- (left-), EXC- (middle-) and EXC ERM-1[++] modifier screens (right column), arranged by trafficking function (colored boxes) and phenotype class (columns; bra-/+: branch loss/gain, E: enhancers, S: suppressors; Fig. 1B for screen layouts, classes, acronyms). Phenotypes are distinguished as: none (0, no grey), weak (1, light grey), strong (2, grey) and phenotype that led to identification of gene (3, dark grey). Results of repeat RNAi screens are included. List does not include: genes identified by early combined tissue and membrane polarity defects; genes subsequently identified by membrane polarity defects (*sec-23, copz-1, syx-7, ggtb-1, vha-1, -2*; see text). *Genes recovered in tier-2 screens as GSL enhancers. The following gene products (all phylogenetically conserved) have documented physical or functional interactions in various species: VHA-1/-2/-4/-5/-6/-8/-9/-10/-11/-12/-15/-18, SEC-23/- 24 (protein complex subunits); SYX-5/YKT-6, SYX-5/SLY-1, SYX-5/SEC-24.1, SYX-5/RAB- 1, YKT-6/SYX-7, RAB-1/SEC-23/SEC-24.1, DAB-1/APT-9, APT-9/ARF-1.1 (interacting proteins; **Tab.S1** for references). All 50 trafficking genes are expressed in the *C. elegans* INT and/or EXC by RNAseq (*94*) and/or other experimental studies (see text). **(B)** Selection of corresponding INT/EXC embryonic/larval apical membrane expansion, polarization and positioning defects. Not all phenotypes are shown/indicated. See **Fig.S2** for description of these and additional phenotypes. Top panel: Nomarski/confocal overlay of wild- type (wt) late (3fold) embryo (left) and late L1 larva (right). 2-3fold late embryos shown for most genes; L1-larvae for *ggtb-1, vab-8, eps-8, vha-5, vha-6, vha-8*; early embryos for *sec-23, sec-24* and *ykt-1* (INT and EXC tubulogenesis arrest stages can be discordant). Early-embryonic phenotypes with concomitant tissue morphogenesis defects are not shown (except for INT intercalation defects). Lum morph defect: lumen morphogenesis defect (see **(A)** for other acronyms). Confocal projections and sections are shown (all specimens are fully sectioned to determine lengths of canals). White arrows point to INT, yellow arrows to EXC (arrowheads in wt mark canal tips). ERM-1::GFP-labeled animals shown throughout (Methods for ERM-1::GFP and other fluorescently-labeled fusion proteins used in this study). Scale bars: 10μm. Color coding of functional classes (boxes) as in Fig. 1C. The spectrum of phenotypes (depending on gene, developmental stage, strength and time of interference; Methods) ranges from severe/moderate (‘no/partial apical membrane polarization’ = full/partial ERM-1 displacement/INT; ‘no/short lumen/no/incomplete apical membrane expansion’ = no ERM-1 extension/EXC) to wild-type. Basolateral (BL) ERM-1 mislocalization (with/without residual apical ERM-1) indicates an apical domain positioning or apicobasal membrane polarity defect. Defects in ‘apical membrane polarization’ (absence of ERM-1 at the apical domain) are distinct from defects in ‘apical domain positioning’ (BL ERM-1 mislocalization). ‘No apical membrane polarization’ (full ERM-1 displacement from the membrane) is distinct from ‘no apicobasal membrane polarity’ (pan membranous ERM-1): *dab-1* provides an example for the former, *sec-23/24* for the latter; see SFig.2 for additional examples). 36/50 images show combined INT/EXC phenotypes, with 16/36 canals revealing ‘no/minimal lumen/ERM-1+ apical membrane extension’.

Since these 50 genes were isolated from different screens (intestine or canal) and by different phenotypes, we carried out a repeat RNAi analysis of all 50 genes to comprehensively examine their effects on both tubes (intestine and canal), all aspects of apical membrane biogenesis (phenotype classes 1-10), and all developmental stages (note that most knockdowns are sterile or embryonic/early-larval lethal). Further validating screen quality, this analysis revealed unexpected common properties among these genes’ functions. Most operated in: (1) both, intestinal and canal lumenogenesis (42/50 genes; Figs.2B, S2, Tab.S1) and in similar aspects of apical membrane biogenesis in these different tubes (Fig. 2A for phenotype signatures across classes 1-10); (2) multiple aspects of apical domain biogenesis in each tube (43/50 genes; Fig. 2A; notably, some genes identified by their requirement for canal lumen expansion were also found to regulate intestinal polarity [class 4, Fig. 1B]); (3) both, embryonic and larval development, i.e. during and after completion of polarized tissue (tube) morphogenesis (44/50 genes; Figs.2, S2, Tab.S1)

In sum, tier-1 screens identify 50 trafficking molecules required for apical domain and lumen morphogenesis in multi- or unicellular tubular *C. elegans* epithelia. Their comparative analysis reveals that a shared trafficking machinery supports the different tissue-morphogenetic processes of inTERcellular/intestinal and inTRAcellular/canal lumenogenesis and the distinct membrane-biosynthetic processes of apical membrane polarization, positioning, and expansion. All genes function in growing epithelia, i.e., during *de novo* apical membrane and lumen biogenesis, and most function in both embryonic cells that still divide and move and in larval cells that no longer divide and move but still expand.

### Genetic modifiers of the GSL-dependent apicobasal polarity conversion identify new trafficking molecules that regulate apicobasal membrane polarity

We next used these 50 trafficking molecules with documented functions in apical membrane biogenesis to investigate whether GSLs indeed regulate apicobasal membrane polarity via trafficking, and if so, by which trafficking route (see Introduction). We carried out genetic enhancer and suppressor screens (tier-2 screens) and examined the ability of each of the 50 tier-1 trafficking genes to modify ERM-1’s membrane position in a GSL-biosynthetic-enzyme-deficient mutant (*let-767[s2819]*; Fig. 3B). GSL depletion results in basolateral displacement of all tested integral and submembranes apical membrane components, with subsequent ectopic basolateral lumen and junction formation (apicobasal polarity conversion; Fig. 3A) (*7*). ERM-1::GFP tracks this full membrane domain identity change. We defined enhancement as ERM-1 emergence at basolateral domains in initially wild-type-appearing intestines that were only weakly sensitized by the loss of one *let-767* copy. RNAi was induced post-embryogenesis, requiring enhancers to directly affect polarized membrane biogenesis, independent from polarized tissue morphogenesis (see above, approach, and Fig.S1). We defined suppression as the prevention/reversion of basolateral ERM-1 displacement and ectopic lumen formation and/or arrest/lethality in the absence of both *let-767* copies. RNAi was induced before embryogenesis, requiring suppressors to rescue/revert the fully expressed phenotype.

**Figure 3.**
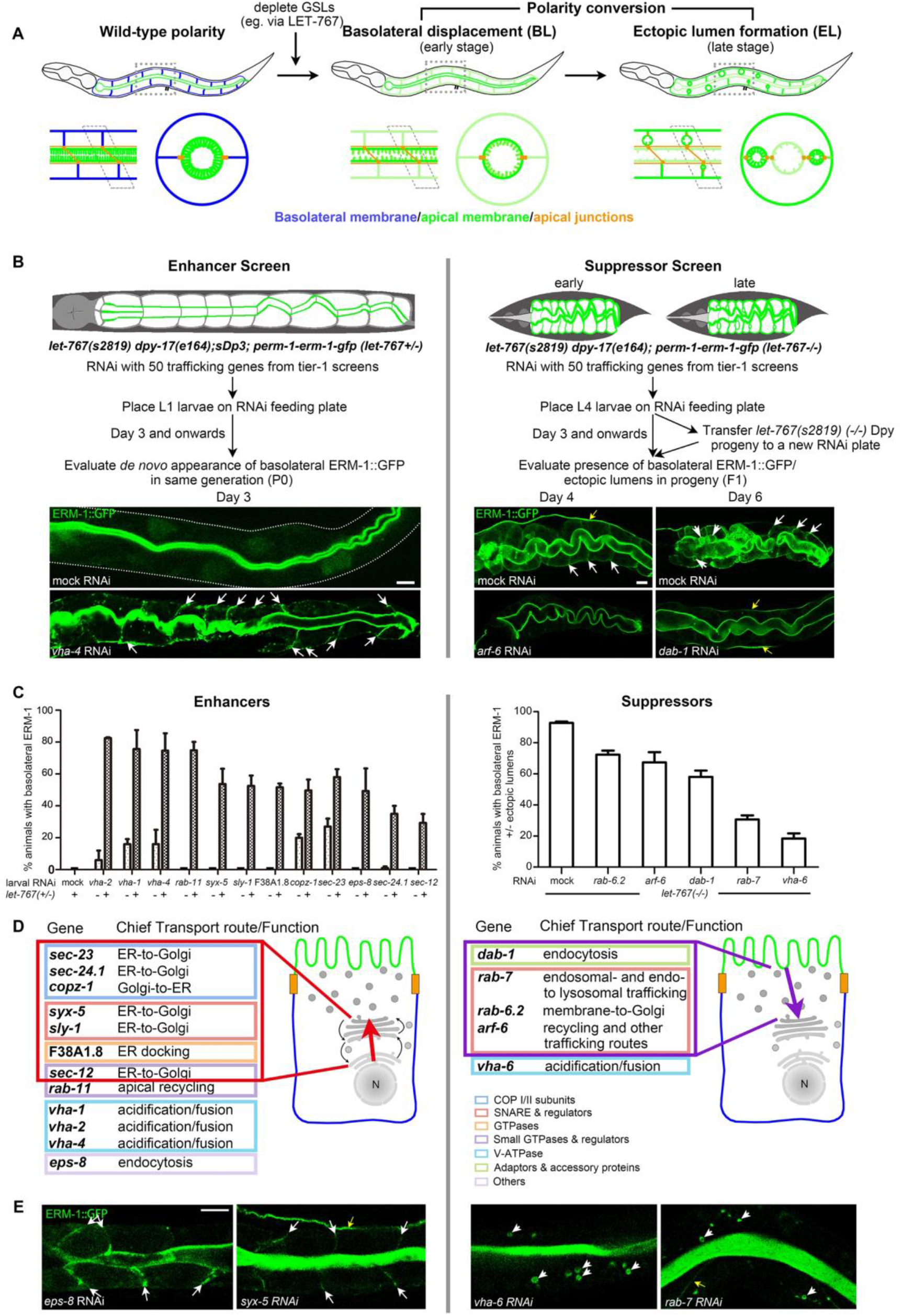
Enhancers and suppressors of the GSL-dependent apicobasal polarity conversion. **(A)** Schematics of *in vivo* apicobasal polarity conversion. Top: whole larva; bottom: longitudinal (left) and transverse section (right) through 1 INT ring. Basolateral (BL, here and below) displacement of apical membrane components precedes ectopic lumen formation (EL, here and below). Note loss of apical versus gain of BL microvilli and intact apical, but new BL lumen-surrounding junctions, indicating full structural apical membrane-, membrane-microdomain and apical domain formation at (prior) basolateral side. Over time, ERM-1 and apical membrane characteristics (microvilli) are lost from the (prior) apical domain (*7*) (compare **Fig.S6A, B** for timing of events and polarity reversion). **(B)** Genetic interaction screens (enhancer screen left, suppressor screen right). Top: schematics, left: *let-767(+/-)* larva (intact polarity; duplication/*sDp3* provides one *let-767* copy; designated *let-767(+/-)* for simplicity here and below); right: dumpy L1-arrested *let-767(-/-)* larvae (polarity conversion; *sDp3* lost, no *let-767* copy; designated *let-767(-/-)* here and below; note that maternal product is present). ERM-1::GFP identifies apical membrane (green). Middle: screen layout. Enhancer screen (left): phenotype is evaluated in the same generation, with enhancement required to induce BL ERM-1 mislocalization in wild-type appearing *let-767(+/-)/RNAi* INT (see text); suppressor screen (right): phenotype is evaluated in the next generation and requires prevention or reversion of the full phenotype (BL ERM-1/ELs; L1-arrest) in *let-767(-/-)/RNAi* progeny (see text). Bottom images: representative examples of enhancement (left) and suppression (right). Note punctate nature of BL ERM-1 displacement. Compare to **Fig.S3A, B**. **(C-E)** Screen results (enhancers left, suppressors right). **(C)** Quantification of enhancement/suppression. Mean ± SEM shown, n=3/N>50. **(D)** Schematics: genetic modifiers, arranged by conserved primary site of function/route (compare **Figs.1C, 2A**): enhancers are enriched for pre-Golgi components of anterograde biosynthetic-secretory (red rectangle/arrow), suppressors for post-Golgi components of retrograde endocytic-recycling, routes (yellow rectangle/arrow; cartoons as in **Fig.S1A;** color coding of functional classes (boxes) as in Fig. 1C). Note that V-ATPase subunits are among enhancers and suppressors, suggesting specific yet distinct roles of pump subunits (subunit isoforms can assemble in an organelle-specific manner (*35*); see the VHA-6 analysis below and **Fig.S7A** legend for discussion). **(E)** Representative images of membrane-positional (BL) ERM-1 displacement (enhancers, left) and vacuolar cytoplasmic ERM-1 displacement (suppressors, right) in wild-type INT. Confocal sections of whole **(B)** and partial **(E)** larval INTs are shown. White arrows: BL mislocalization; arrowheads: ELs in **(B**)/cytoplasmic vacuoles in **(E)**. Yellow arrows: EXC. Scale bars: 5μm.

These tier-2 screens recovered an unexpectedly large percentage of tier-1 trafficking molecules as modifiers of the GSL-dependent apicobasal polarity change (17/50; Figs.3C, S3A-B), supporting the hypothesis that GSLs regulate membrane polarity via trafficking and underscoring GSLs’ critical contribution to such a trafficking-based polarity function. However, few of these 17 trafficking molecules came from functional classes previously implicated in apical sorting or other aspects of polarized trafficking (only 6/26 such molecules identified in tier-1 screen: e.g., 3/13 Rabs/Rab-related and 0/6 motors), whereas most lacked such implication (e.g., 5/6 vesicle coat/coat assembly components; see color-coded boxes in Figs.1C, 2A, 3D).

Intriguingly, GSL enhancers (12) and suppressors (5) fell into different classes based on their conserved wild-type trafficking functions (Fig. 3D, Tab.S1). Notably, enhancers, whose mild depletion mispositions the apical domain in a haplosufficient *let-767[s2819; sDp3]* background (Fig. 3B), were enriched for biosynthetic-secretory pathway components of membrane-directed (anterograde) vesicle trajectories (7/12), implicating this pathway in the regulation of membrane polarity and suggesting that GSLs position the apical domain on biosynthetic-secretory or parallel trafficking routes. Validating this finding, enhancers also identified molecules previously implicated in apical domain positioning via apical recycling or by functions at the apical-membrane-vesicle interface (RAB-11, VHA-1/4, EPS-8; (*19, 33, 38-40*) note that clathrin/AP-1, previously characterized as GSL enhancers (*11*), are not included in this analysis). In contrast, suppressors, whose strong depletion reduces apicobasal polarity conversion in a *let-767(-/-)* background (Fig. 3B), were enriched for components of (retrograde) endocytic-recycling and vesicle-degradative trajectories leading away from the membrane (Fig. 3D, Tab.S1).

Enhancers and suppressors also fell into different classes based on their loss-of-function phenotypes in a wild-type background. Hierarchical clustering of the 50 tier-1 knockdowns by phenotype profile could predict whether they would enhance or suppress GSLs’ polarity function (Fig.S3C). The strongest predictor for enhancement was a combined defect in wild-type apical membrane positioning and polarization (basolateral and cytoplasmic ERM-1 displacement; 8/12 enhancers, 0/5 suppressors). An extended analysis of polarized membrane expansion in postmitotic larval cells (using a graded scale of RNAi strength and number of time points at which expanding membranes were analyzed; Methods) revealed that depleting each enhancer (12/12) was independently able to displace ERM-1 to basolateral domains, whereas depleting suppressors (5/5) displaced ERM-1 to cytoplasmic vacuoles or had no effect (Fig. 3E). We conclude that all trafficking molecules that interact with GSLs in membrane polarity are themselves required for apical domain polarization, but only enhancers are required for the regulation of apicobasal membrane polarity.

Among GSL enhancers, tier-2 screens thus identify multiple new trafficking molecules that position the apical domain on expanding wild-type *C. elegans* intestinal membranes, most of them biosynthetic-secretory molecules, while confirming the role of several endocytic-recycling molecules in this function. In contrast, GSL suppressors, chiefly operating on endocytic and vesicle-degradative routes, seem dispensable for the regulation of wild-type polarity although required for the GSL-dependent polarity conversion. We conclude that the trafficking molecules that enhance and suppress GSLs are distinguished by their: (1) primary site of function in wild-type (antero-versus retrograde trafficking routes); (2) role in wild-type apical membrane biogenesis (apical domain positioning versus polarization). This constellation suggested enhancer- and suppressor specific distinct mechanisms of action and interaction of these trafficking molecules with GSLs in membrane polarization that might involve trafficking directions. Significantly, these sensitized tier-2 screens identify multiple unequivocal biosynthetic-secretory pathway components that regulate apicobasal membrane polarity.

### The biosynthetic-secretory pathway specifies the polarized distribution of apical membrane and PAR-complex components during *de novo* membrane biogenesis

The biosynthetic-secretory (secretory) pathway delivers biosynthetic cargo from the ER via pre-Golgi, Golgi- and post-Golgi endo-/vesicle membranes to all sides of the epithelial plasma membrane. Its apicobasal directionality is thought to be determined by polarized cargo signals that are decoded only on the level of Golgi- and post-Golgi endomembranes (e.g. by GSLs) to sort post-Golgi vesicles to cognate recognition sites at corresponding target membranes (e.g. the apical domain) (*4, 41*). Pre-Golgi (early) pathway components are thus not expected to specify this cargo’s apicobasal distribution at the membrane. Yet, interference with each of 7, predominantly early pathway components (the COPI/II coat/coat assembly components SEC-23, SEC-24, SEC-12, COPZ-1; the SRPR ortholog F38A1.8; the t-SNARE SYX-5 and its assembly factor SLY-1), enhances the GSL-dependent apicobasal polarity conversion and shifts ERM-1 to basolateral domains of expanding membranes in the *C. elegans* intestine (Fig. 3C-E, S3A, TabS1). Non-specific or secondary effects of early pathway disruption on any apical delivery route (direct, indirect or transcytotic; Fig. 6A) would be expected to displace ERM-1 to the cytoplasm, not to the basolateral membrane; and to suppress, not enhance, apicobasal polarity conversion (by reducing cargo flux; see the suppressor analysis below). Disrupting upstream (pre-Golgi) trafficking should preclude compensatory mechanisms in downstream (post-Golgi) trafficking (e.g., increased basolateral delivery) that depends on it. We therefore conclude that the secretory pathway has novel, non-canonical functions in apical sorting or affects membrane polarity by some other unknown mechanism in this expanding epithelium.

To further investigate the secretory pathway’s polarity function, we re-examined all pathway components identified among the 453 molecules identified in tier-1 screens by their requirement for apical domain biogenesis. Although only 7 of 14 such components were recovered as GSL enhancers in tier-2 screens, 3 more displayed an enhancer-specific phenotype signature in wild-type (Fig.S3C). An extended RNAi analysis of all 14 components (as above; Methods) identified 4 more secretory molecules required to position the apical domain (RAB-1, YKT-1, SYX-7, GGTB-1; Fig. 4C). Among different classes of tier-1 trafficking molecules, apicobasal polarity defects thus track with the loss of secretory pathway components (Fig. 4A, Tab.S2). In total, we identify 14 such components, with conserved sites of function ranging from ER-, pre-Golgi-, Golgi- and post-Golgi endomembranes to the plasma membrane (Fig. 4B). 11 are required for apical domain polarization and positioning in the intestine (Fig. 4B, pink), 7 in conjunction with GSLs (Fig. 4B, pink/boxed). All 14 expand the apical membrane between pairs of cells (intestine) and inside one cell (excretory canal). We conclude that the secretory pathway harbors directional (apical) cues throughout its endomembrane system and concomitantly expands, polarizes, and positions the apical domain. This finding suggests that, at the time of polarized membrane expansion in developing epithelia, the directionality of this anterograde vesicle trajectory is fixed from the ER towards the nascent apical domain.

**Figure 4.**
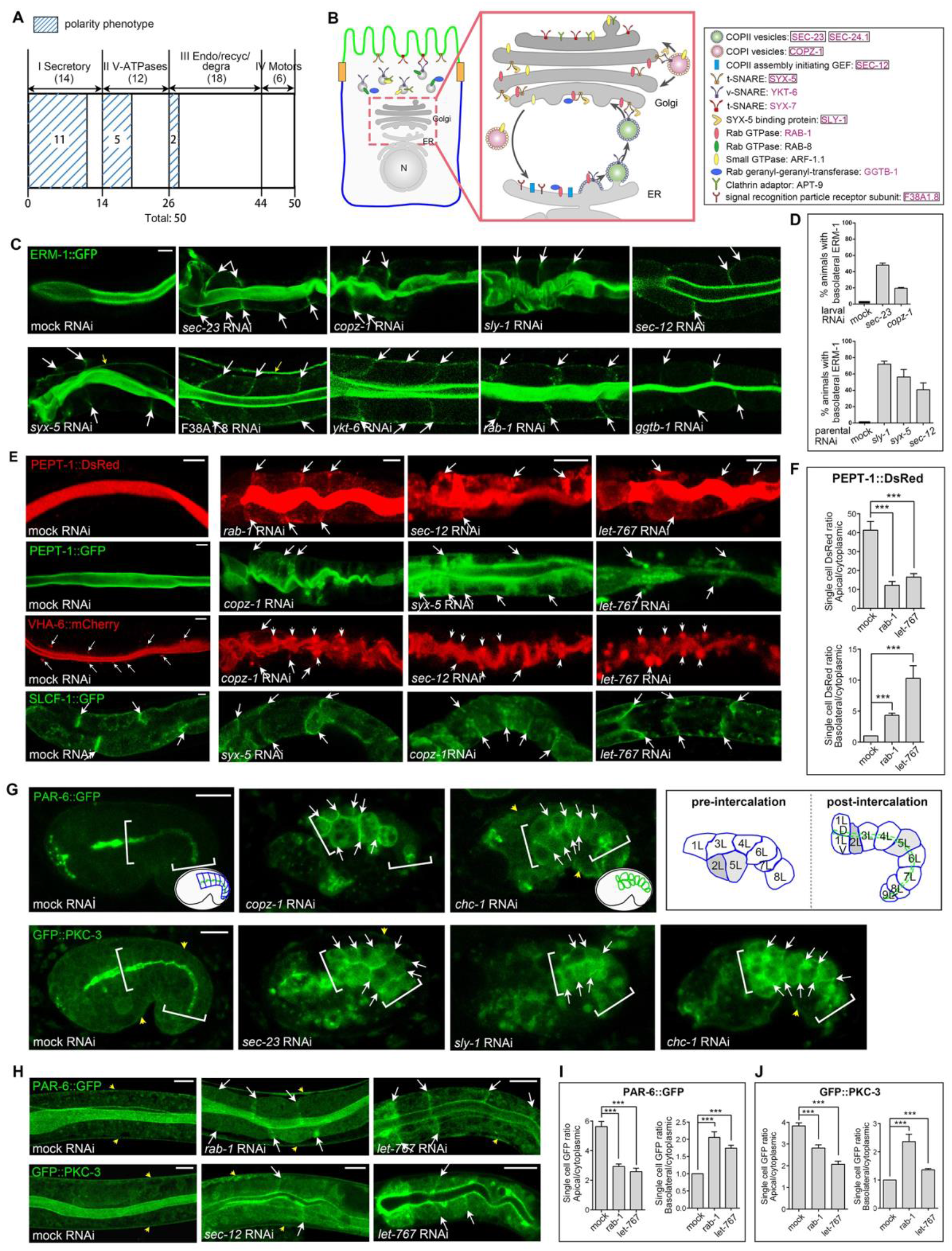
Interference with the biosynthetic-secretory pathway displaces expanding apical to basolateral domains during polarized membrane biogenesis. **(A)** Among 50 trafficking molecules for apical domain biogenesis, polarity defects (basolateral/BL ERM-1 mislocalization) track with the loss of secretory molecules. Molecules are classified by their conserved trafficking functions/routes (endo/endocytosis; recyc/recycling; degra/degradation; see text and **Tabs.S1**, **S2**). **(B)** Endomembrane location of 14 secretory pathway components identified by their requirement for apical domain biogenesis (compare **Tab.S1, S2**). Components required for wild-type membrane polarity pink; enhancers of GSLs’ polarity function boxed. Cartoon of polarized cell as in **Fig.S1A.** **(C-D)** Mild interference with each of multiple secretory pathway components misdirects the apical domain identity marker ERM-1 to BL membranes during polarized membrane biogenesis (white arrows; **Fig.S1**). Set of 2 - 3 opposing cells of expanding larval intestines are shown; yellow arrows EXC. **(D)** Quantification of polarity defects subsequent to parental (top) and L1 RNAi (bottom; see text). Mean ± SEM shown, n=3/N>50. **(E-F)** Effects of mild secretory pathway perturbation on the polarized distribution of integral membrane components on expanding membranes of wild-type larval INT cells: cytoplasmic and BL (arrows) fluorophore-independent mislocalization of the apical peptide-transporter PEPT-1; BL (arrows) and BL-aligned vacuolar (short arrows) mislocalization of the apical V-ATPase subunit VHA-6 (thin arrows indicate presence of small VHA-6+ subapical vesicles in wild-type; compare Fig. 6F); minor cytoplasmic/punctate but no apical displacement of the BL (arrows) sugar transporter SLCF-1. Defects copy the mild GSL-dependent polarity conversion (BL ERM-1, no ectopic lumens/ELs (*7*); *let-767(mildRNAi)* INT shown for comparison). **(F)** Quantification of PEPT-1::dsRed apical membrane/cytoplasm (top) and BL membrane/cytoplasm (bottom) fluorescence intensity ratios (3 cells/animals used for fluorescence intensity quantification; >10 animals for calculating the ratio). **(G-I)** The polarized distribution of apical PARs depends on the secretory pathway. **(G)** Failure of PAR-6/PKC-3 polarization in intercalating *copz-1-, sec-23-, sly-1(mild-to-moderateRNAi)* early-embryonic INT (arrows indicate BL mislocalization; PAR-6 and PKC-3, like ERM-1, are strictly apical in wild-type). The phenotype copies the *chc-1(RNAi)* induced polarity conversion, shown in comparison (*11*)). Note concomitant intercalation defects (compare schematics to the right and **Fig.S1A**; (*42, 43*)). Inset shows cartoon of embryo outline in the eggshell. Strong secretory pathway disruption fully displaces PAR-6/PKC-3 to the cytoplasm and prevents intercalation and lumen formation (not shown). Brackets: INT; yellow arrows: hypodermal PAR-6 and PKC-3. **(H)** BL (arrows) and cytoplasmic PAR-6/PKC-3 mislocalization in *rab-1- and sec-12(mildRNAi)* larval INT (compare to *let-767(mildRNAi)* phenocopy (*7*)). Set of 2 opposing cells are shown. **(I-J)** Quantification of PAR-6::GFP and GFP::PKC-3 apical membrane/cytoplasm and BL membrane/cytoplasm fluorescence intensity ratios. Methods as in **D/F**. Mean +/- SEM shown, ***p<0.001. Confocal sections of portions of expanding larval INTs are shown in C, of whole embryos in G. Scale

Strong interference with the essential secretory pathway aborts all membrane-directed traffic, making it impossible to detect its putative effect on membrane polarity (Tab.S1). To analyze the pathway’s polarity function throughout development, we used conditional and reduced-strength loss-of-function conditions with single pathway components (Methods). Mild *sec-23-, copz-1, sly-1, sec-12-, syx-5-, F38A1.8, ykt-6,* and *rab-1* RNAi each mislocalized ERM-1 to basolateral sides of expanding membranes in postmitotic late-embryonic and larval intestinal cells (Figs.4C, S1). Conditional mild RNAi, induced in L1-larvae (ongoing net membrane expansion), was sufficient to change polarity, whereas strong RNAi, induced in adults (no/minimal membrane expansion), had no such effect (Fig. 4C, D; not shown). We conclude that the secretory pathway’s role in membrane polarity is: (1) a direct function of polarized membrane biogenesis (still effective after completion of polarized tissue morphogenesis); (2) developmentally regulated (restricted to embryonic and larval epithelia); (3) linked to membrane expansion (dispensable for polarity maintenance in fully expanded epithelia).

Mild *sec-23-, sec-12-, syx-5-, sly-1-* and *copz-1* RNAi also displaced other apical membrane components (submembranous actin/ACT-5 and intermediate filaments/IFB-2; the integral membrane components PEPT-1 and VHA-6, a V-ATPase subunit also identified as a GSL suppressor; see below) to the basolateral domain, the cytoplasm and to cytoplasmic vacuoles. In contrast, it only modestly displaced the basolateral integral membrane protein SLCF-1 to the cytoplasm, without misdirecting it to the apical domain (Figs.4E-F, not shown). Apical junction integrity was intact, suggesting that junction leakage did not cause the polarity change; yet ectopic junction material could be observed in the cytoplasm and at lateral membranes (Figs. 5A-C, S5, MovieS1). We conclude that the secretory pathway’s role in the polarized distribution of membrane components is not limited to ERM-1 and is specific to apical membrane, including apical junction, components.

**Figure 5.**
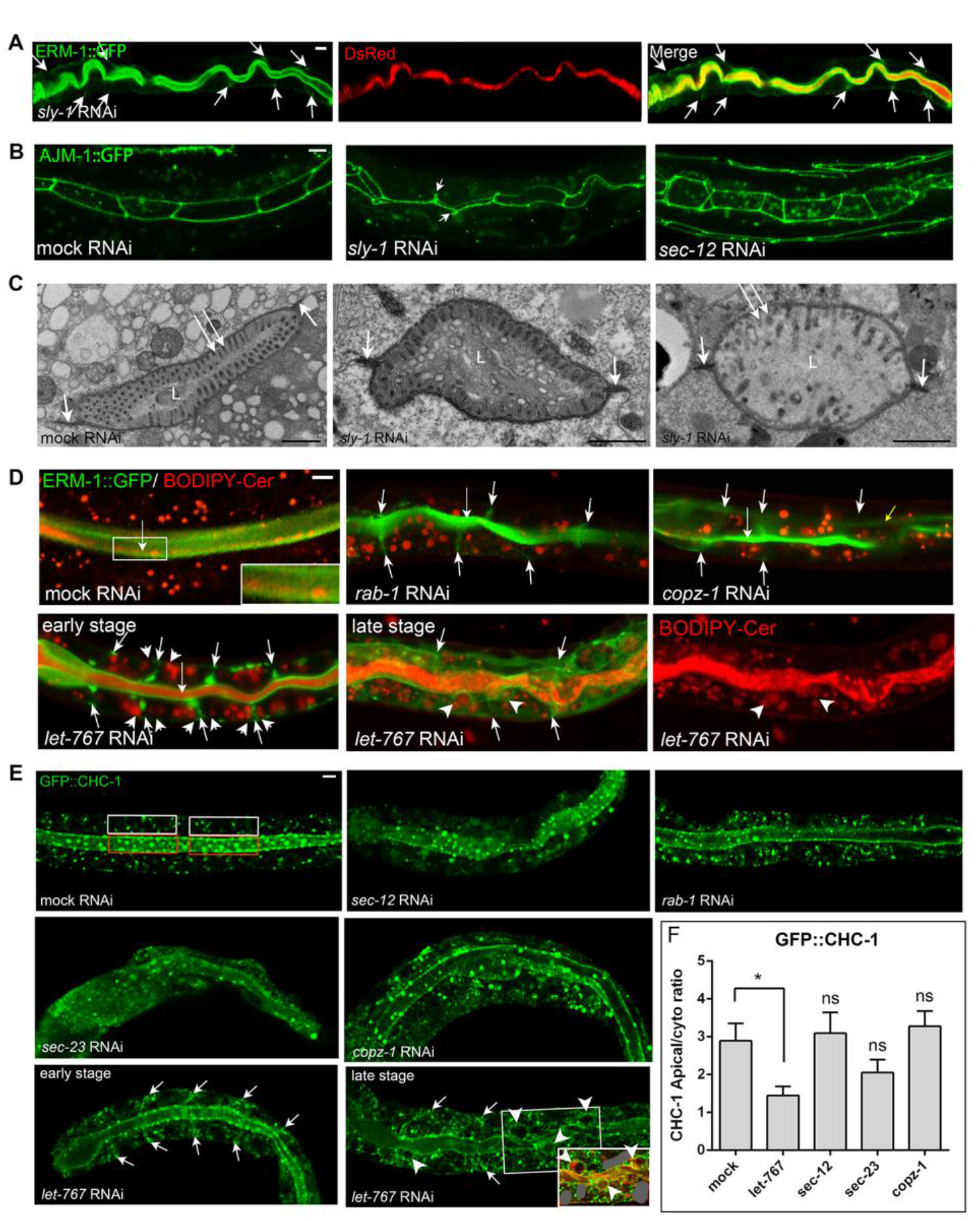
Mild perturbation of pre-Golgi trafficking leaves junctions intact and affects the biogenesis but not directionality of post-Golgi vesicles. **(A-C)** Apical junction integrity is not affected by *sly-1-* and *sec-12* mild RNAi. **(A)** Larvae fed with dsRed-labeled bacteria retain the label in the lumen despite basolateral (BL) ERM-1 displacement (arrows). **(B)** The junction integrity molecule AJM-1 remains contiguous at its apicolateral position (with occasional lateral broadening; short arrows; see **Fig.S5A** for *sec-23-* and *copz-1(mildRNAi)* INT and ectopic cytoplasmic junctional material; **Movie S1** for junction contiguity [rotation of 3D projection]). **(C)** Transmission electron microscopic/TEM images: structurally intact junctions (short arrows) in *sly-1(mildRNAi)* INT (increasing phenotype severity from left to right; representative images shown). Note reduced/absent number and length of apical microvilli (long arrows), deformed lumen (L), paucity and dissolution of endomembranes (**Fig.S5B** for whole intestines). **(D-F)** Defects in GSL-rich [BODIPY-Cer+] and clathrin-coated [CHC-1::GFP] post-Golgi vesicle biogenesis but not positioning in *rab-1-, copz-1, sec-23, sec-12(mildRNAi)* INTs. **(D)** BODIPY-Cer+ vesicles. Left to right, 1^st^ row: pancytoplasmic distribution in wild-type (note BODIPY-Cer at outer leaflet [=intralumenal] of the apical membrane [enlarged in inset], absent in *rab-1-* and *copz-1(mildRNAi)* INT cells); no change in distribution, but reduction in number and increase in size in *rab-1-* and *copz-1(mildRNAi)* INT cells. BL ERM-1 displacement (short arrows) demonstrates presence of polarity defect. 2^nd^ row: compare to positioning change during *let-767(RNAi)* polarity conversion: BL displacement to both sides (very short arrows) of ERM-1+ prior BL membrane (short arrows) in early-stage-, and displacement to ERM-1+ ectopic lumens/ELs (2 ELs indicated by arrowheads) in late-stage, polarity conversion (note BODIPY-Cer at the outer membrane leaflet [=intralumenal] of ELs). **(E)** CHC-1::GFP+ vesicles. Left to right, 1^st^ and 2^nd^ row: pancytoplasmic distribution and apical enrichment in wild-type; reduced number but sustained albeit reduced apical enrichment in *sec-12-, rab-1-, sec-23-* and *copz-1(mildRNAi)* INT cells (note that minor positional changes could be obscured by reduced number of vesicles). 3^rd^ row: compare to positioning change during *let-767(RNAi)* polarity conversion: BL displacement (arrows) in early-stage, and displacement to ELs (arrowheads) in late-stage, polarity conversion; inset: ERM-1::mCherry documents apical character of EL membrane (additional ERM-1 greyed out for clarity). **(F)** Quantification of GFP::CHC-1 apical-membrane/cytoplasm ratio. Two areas (60 sqmm) per INT cell were counted for the apical membrane (red box in **E**) and the cytoplasm (white box in **E**) and >10 worms were used for the calculation. Mean ± SEM shown, *p < 0.05. Confocal sections of portions of expanding larval intestines are shown (projections in **A**). Scale bars: 5μm; 1μm in (**C**).

**Figure 6.**
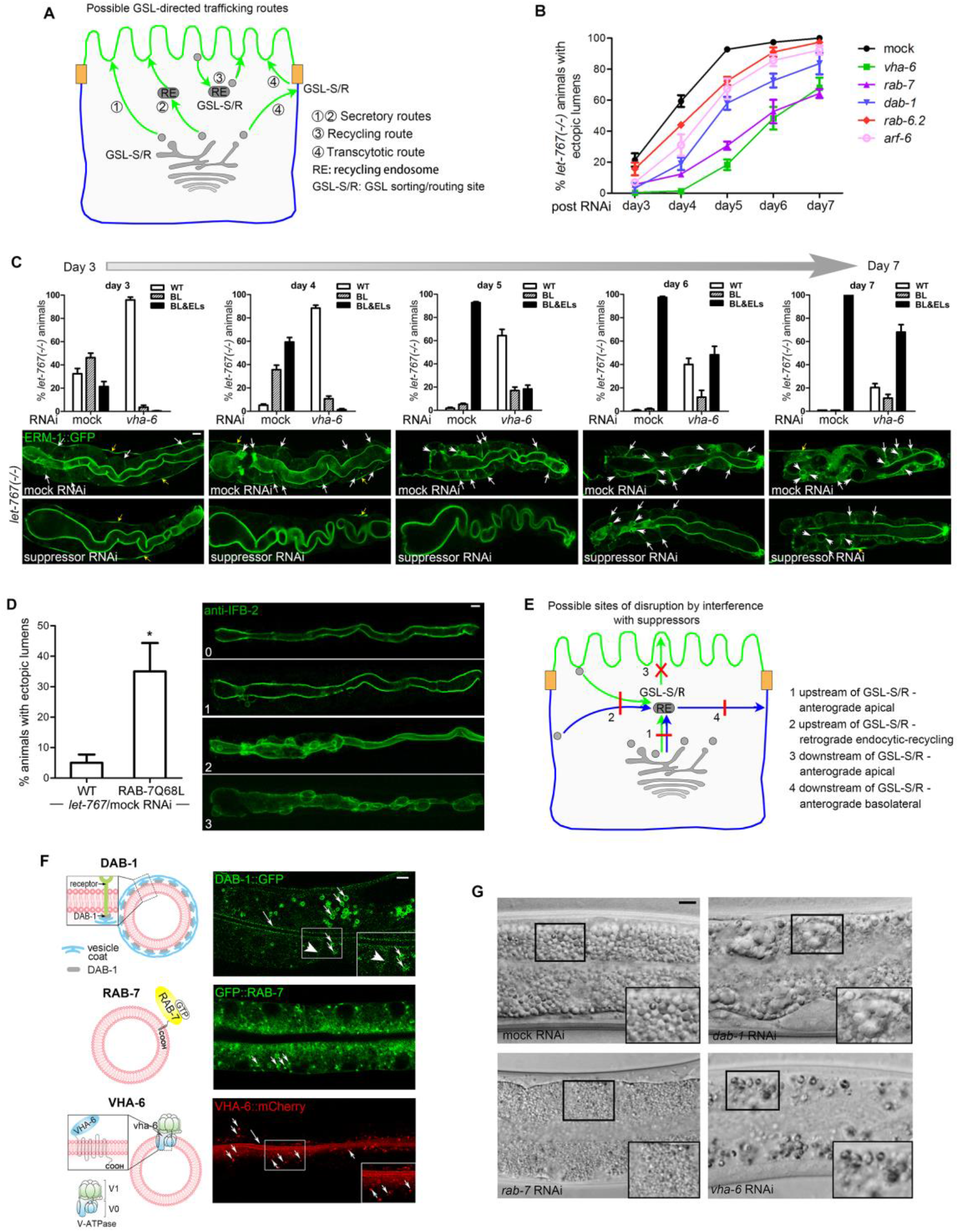
Depletion of VHA-6, RAB-7, DAB-1, ARF-6, or RAB-6.2 suppresses the GSL-dependent apicobasal polarity conversion. **(A)** Possible GSL-directed post-Golgi trafficking routes to the apical membrane (see text). Anterograde secretory routes could be direct (1) or indirect (2), i.e. passing through recycling, sorting or other endosomes (recycling endosome/RE shown for simplicity). Recycling routes (3) could pass through different endosomes and Golgi/ER compartments (retrograde recycling, not shown). Possible GSL sorting or routing sites (GSL-S/R) are indicated. Note, that on transcytotic routes (4), such sites must reside at or beyond the basolateral (BL) membrane (apical cargo accumulates on BL membranes in GSL-depleted epithelia). **(B)** Suppression of polarity conversion, expressed as percent *let-767(-/-)(RNAi)* larvae with ectopic lumens/ELs over time (compare **(C)** below). **(C)** Depleting suppressors reduces polarity conversion but does not induce reversion (VHA-6 shown as example). Compare to **(B)**, see **Tab.S3** for N of all 5 suppressors, **Figs.S6A-B** and legend for GSL-dependent polarity conversion/reversion in growth-delayed larval INTs. Bar graphs show *let-767(-/-)/vha-6(RNAi)* larval INTs with changing proportions of: wild-type (WT); early-stage (BL ERM-1::GFP displacement; arrows), and late-stage (ELs; arrowheads), polarity conversion over time. Corresponding representative images of each stage shown below (note that all images show *let-767(-/-)* INTs; yellow arrows: EXC). Mean ± SEM shown. **(D)** Activated RAB-7 enhances polarity conversion. Increase in number of *let-767(mildRNAi)* animals with intestinal ELs in the presence of a RAB-7Q68L transgene (left). Increase in EL number and size per animal increases proportionally to increase in number of animals with ELs (right: progression was graded from 0-3: larval intestines shown, quantification not shown; Methods). Anti-IFB-2 marks apical intermediate filaments. Mean ± SEM shown, n=3/N=20. **(E)** Possible sites of disruption of post-Golgi trafficking routes by interference with suppressors (see text). Only disruption at red bars (routes 1, 2, 4) reduces BL misrouting under GSL depletion conditions; disruption at red X (route 3) is neutral or enhances misrouting (except if suppressor inhibits). GSL sorting/routing site (GSL-S/R) arbitrarily placed at recycling endosome (RE). Routes upstream of GSL-S/Rs could be secretory (route 1) or endocytic-recycling routes (routes 2), downstream routes could be either. Only post-Golgi routes are shown, and predictions are based on current maps of polarized trafficking in epithelia (see **A**). Routes are colored (green: apical; blue: basolateral) according to destination, not cargo (apical cargo could be misrouted on a BL route). **(F)** Subcellular localization of DAB-1, RAB-7 and VHA-6 in wild-type larval INTs. Schematics (left): relation to endomembranes. Images (right): localization on membranes/endomembranes (long arrows: apical-, arrowheads: BL membranes; short arrows: vesicles). DAB-1, thought to be restricted to BL membranes (*46*), was also found on apical membranes and on 2 distinct vesicle populations (compare **Fig.S6F**); VHA-6, thought to be restricted to apical membranes (*38, 47*), was also found on subapical vesicles (short arrows). RAB-7’s vesicular localization, previously characterized, (*48*) is shown for comparison (RAB-7 colocalizes with early and late endosomal-, lysosomal- and LRO markers and also forms subapical aggregates in L1 larval intestines; not shown). (*7, 48*) **(G)** Vesicle biogenesis defects in *dab-1-, rab-7-, vha-6(RNAi)* larval INTs. Note vacuolization and aggregation (*dab-1*), size reduction (*rab-7*) and reduced number (*vha-6*) of vesicles. Confocal images of larval INTs are shown (Nomarski images in **G**); full INTs in **C, D**, 2 pairs of opposing cells in **F, G**. Scale bars: 5μm. Cartoons of polarized cells/colors (**A, E)** as in **Fig.S1A.**

We considered that a secretory pathway whose directionality is fixed towards the apical domain during *de novo* polarized membrane biogenesis might operate upstream of membrane-based apical polarity cues and also initiate membrane polarization. Indeed, stronger interference with multiple pathway components prohibited membrane polarization in the intercalating early-embryonic intestine, at the time when its definitive polarity and lumen position are initially established (Fig.S1A). In these still dividing and moving cells, ERM-1::GFP was either fully displaced to the cytoplasm (with and without ERM-1::GFP+/- vacuoles; Fig. 2B: e.g.: *sec-23, copz-1, syx-5, ykt-6*) or, if still membrane-associated, non-polarized (panmembranous ERM- 1::GFP; Figs.2B, S2, e.g.: A, B, J, P; note concomitant intestinal intercalation arrest and absence of intestinal and canal lumenogenesis) (*42, 43*). Moreover, moderately strong RNAi with *sec-23-, sec-12-, syx-5-, copz-1* and *rab-1* prevented polarization of PAR-6 and PKC-3 during polarity establishment in the early-embryonic intestines, while mild RNAi mislocalized these apical PAR complex components to basolateral domains in postmitotic but still expanding late-embryonic/larval intestines (Fig. 4G-J; not shown). We conclude that the secretory pathway specifies the polarized distribution of apical membrane and PAR complex components on the expanding membrane of developing intestines, whether or not this membrane is already polarized. The panmembranous distribution of strictly apical PAR complex components in *sec-23-, copz-1- and sly-1(RNAi)* early embryonic intestines (Fig. 4G) reveals that the secretory pathway’s directional (not only secretory) property is required to initiate membrane polarization.

The early (pre-Golgi) secretory pathway is thought to be required for the biogenesis of all post-Golgi vesicles, but not their directionality (see above; (*41, 44*)). The enrichment of polarized cargo *en route* to the plasma membrane makes it difficult to visually track their vesicle-based directional delivery from the ER to the plasma membrane (*4, 9*). To assess possible nonspecific or secondary effects of early pathway disruption on vesicle directionality, we therefore examined the positioning of clathrin-coated- and GSL-rich post-Golgi vesicles, previously shown to change their location from apical to basolateral domains during polarity conversion induced by interference with post-Golgi trafficking (*11*). Mild *sec-23-, sec-12-, copz-1-* and *rab-1* RNAi reduced the number of these vesicles but left their polarized distribution remarkably intact, despite ERM-1’s basolateral displacement (Fig. 5D-F; legends for details). Thus, mild perturbation of the early secretory pathway, sufficient to change membrane polarity, affects post-Golgi vesicle biogenesis but not directionality and fails to collapse the dependent endomembrane system. This finding is consistent with the current model of polarized trafficking that restricts directional cues to Golgi-/post-Golgi endomembranes, but also with the here suggested alternative model where both pre- and post-Golgi endomembranes harbor directional cues required to guide the secretory pathway from the ER towards the apical domain.

We conclude that the biosynthetic-secretory pathway specifies the position of the apical domain on expanding *C. elegans* intestinal membranes, upstream of PARs, from the time of the membrane’s initial polarization in the embryo throughout further net polarized membrane addition required to maintain this polarity in the growing larval, but not in the mature adult, epithelium. The 11 identified pathway components that execute this function have conserved sites of action ranging from the ER to the plasma membrane and are enriched for pre-Golgi vesicle coat and coat assembly components, none implicated in apical sorting. Together, these findings suggest that, during net polarized membrane addition, this anterograde vesicle trajectory acquires apical directionality and is routed from the ER to the plasma membrane to asymmetrically insert the apical domain into the growing membrane, thereby partitioning its apical and basolateral membrane domains.

### Apicobasal polarity conversion is suppressed by decreasing the basolateral misrouting of apical membrane components

In an attempt to characterize this biosynthetic-secretory route for apical domain positioning, we next turned to the GSL suppressors. Tier-2 interaction screens had placed GSLs on, or in parallel to, this anterograde apical trafficking trajectory. Interference with the small GTPases RAB-6.2, ARF-6, and RAB-7, the clathrin adaptor DAB-1/Disabled and the V-ATPase subunit VHA-6 each suppressed apicobasal polarity conversion induced by GSL depletion (Fig. 3C). If these trafficking molecules were effectors of GSL-directed trafficking, reducing their activity should lift an inhibition on apical transport that might reveal this trajectory’s directional regulation, map its itinerary, and identify sites of GSL-dependent cargo sorting or vesicle routing (Fig. 6A). However, the identification of these 5 suppressors in tier-1 screens by their requirement for apical membrane biogenesis suggested, conversely, that they promote apical transport.

To address this difficulty, we first determined when and where the suppressors operate during polarity conversion and reversion (polarity conversion proceeds through stages 1 [basolateral ERM-1 mislocalization] and 2 [ectopic basolateral lumen formation]; reversion proceeds in the reverse order; Fig.S6A). Monitoring membrane expansion in double *let-767(-/-)*/suppressor*(RNAi)* intestines over 7 days (Fig.S6B for timing and sequence of events in GSL-deficient growth-delayed L1-larval intestines) revealed that RNAi with each suppressor reduced the number of *let-767(-/-)* animals with stage 1 and 2 polarity conversion and delayed its progression but failed to restore wild-type polarity and viability (Fig. 6B-C; Tab.S3). Thus, polarity conversion is suppressed by reducing the basolateral misrouting of apical membrane components rather than by increasing their apical delivery. This finding resolved the apparent paradox of the suppressors’ concomitant requirement for apical transport.

Several suppressors have conserved roles on endocytic and vesicle-degradative routes. To assess whether suppression might have been caused by nonspecific disruption of these processes, we depleted various molecules, previously shown to function on these routes in the *C. elegans* intestine, in a *let-767(-/-)* background (Fig.S6C). Depletion of most of these molecules failed to suppress polarity conversion. This analysis did, however, identify two more suppressors (the AP-2 adaptor DPY-23 and the ARF-like GTPase ARL-8), implicating the disruption of specific endocytic routes, but not the general disruption of endocytosis or vesicle degradation, in the mechanism of suppression. We also failed to find evidence for a suppressor-dependent loss of lysosomal GSL degradation that might have been expected to suppress polarity conversion by increasing GSL levels (Fig.S6D-E). If, in contrast, suppression was caused by the loss of gene-specific trafficking functions, increasing these functions should enhance polarity conversion. Consistent with this latter prediction, expression of the constitutively active RAB-7Q68L(*45*) increased the severity of polarity conversion in *let-767(RNAi)* intestines (Fig. 6D).

We conclude that RAB-6.2, ARF-6, RAB-7, DAB-1, and VHA-6 are required for the basolateral mislocalization of apical membrane components in GSL-depleted *C. elegans* intestines, likely affecting this process by their specific functions in trafficking. They would therefore be expected to operate on either: (1) anterograde basolateral trafficking routes downstream of GSL-directed apical sorting/routing sites (Fig. 6E, route 4); or (2) routes upstream of these sites (Fig. 6E, routes 1, 2). Disrupting these routes must reduce basolateral displacement of apical membrane components upon GSL depletion either directly (Fig. 6E, route 4) or indirectly, by reducing influx into the GSL-directed apical trajectory (Fig. 6E, routes 1, 2). These scenarios suggested that the suppressor analysis was unlikely to identify downstream effectors of the GSL-dependent apical trafficking trajectory (Fig. 6E, route 3), but might instead identify trafficking routes supplementing it (Fig. 6E, routes 1, 2, or 4).

### The GSL suppressors DAB-1, RAB-7 and VHA-6 are required for apical and basolateral membrane biogenesis, but not apicobasal polarity, in wild-type intestines

To identify such trafficking routes, we examined the 3 strongest suppressors’ function in wild-type polarized membrane biogenesis. DAB-1, RAB-7 and VHA-6 have documented roles in *C. elegans* intestinal endocytosis, endo-to-lysosomal transport, and lumen acidification, respectively (*45–47*). Tier-1 screens had revealed their shared, suppressor-specific phenotype profiles in wild-type inTER- and inTRAcellular lumenogenesis, marked by defects in apical membrane biogenesis but not polarity (Figs. 2, S2, Tab.S1). Consistent with these suppressors’ trafficking-based function in *de novo* apical membrane biogenesis, confocal analysis of fluorescent fusions detected all 3 molecules on endosomal vesicle- and/or apical membranes in expanding larval intestines, in which *dab-1-, rab-7-* and *vha-6* RNAi each also induced distinct vesicle biogenesis defects (Figs. 6F-G, S6F; legends for details).

We confirmed that these trafficking molecules affect apical membrane polarization but not positioning (and thus membrane polarity) by monitoring apical membrane expansion in *dab-1-, rab-7-,* and *vha-6(RNAi)* postmitotic cells at their fixed positions in growing larval intestines (Fig.S1). ERM-1 and ACT-5/actin were increasingly retained on vacuolar inclusions during net apical membrane expansion but were not misdirected to basolateral domains (Fig. 7A). By confocal microscopy, apical-membrane-bounded vacuolar inclusions are indistinguishable from ectopic lumens, a feature of stage-2 polarity conversion. To characterize the nature of these vacuoles, we performed serial TEM sectioning of *dab-1*-, *vha-6* and *vha-5 rrf-3(RNAi)* larval intestines. This analysis failed to detect vacuoles with inward-pointing microvilli, suggesting they were endomembrane intermediates *en route* to or from the apical membrane, not basolateral ectopic lumens indicative of a defect in the polarized positioning of the apical domain (Figs. 7A, S7A-B; S7A legend for discussion of V-ATPase subunits).

**Figure 7.**
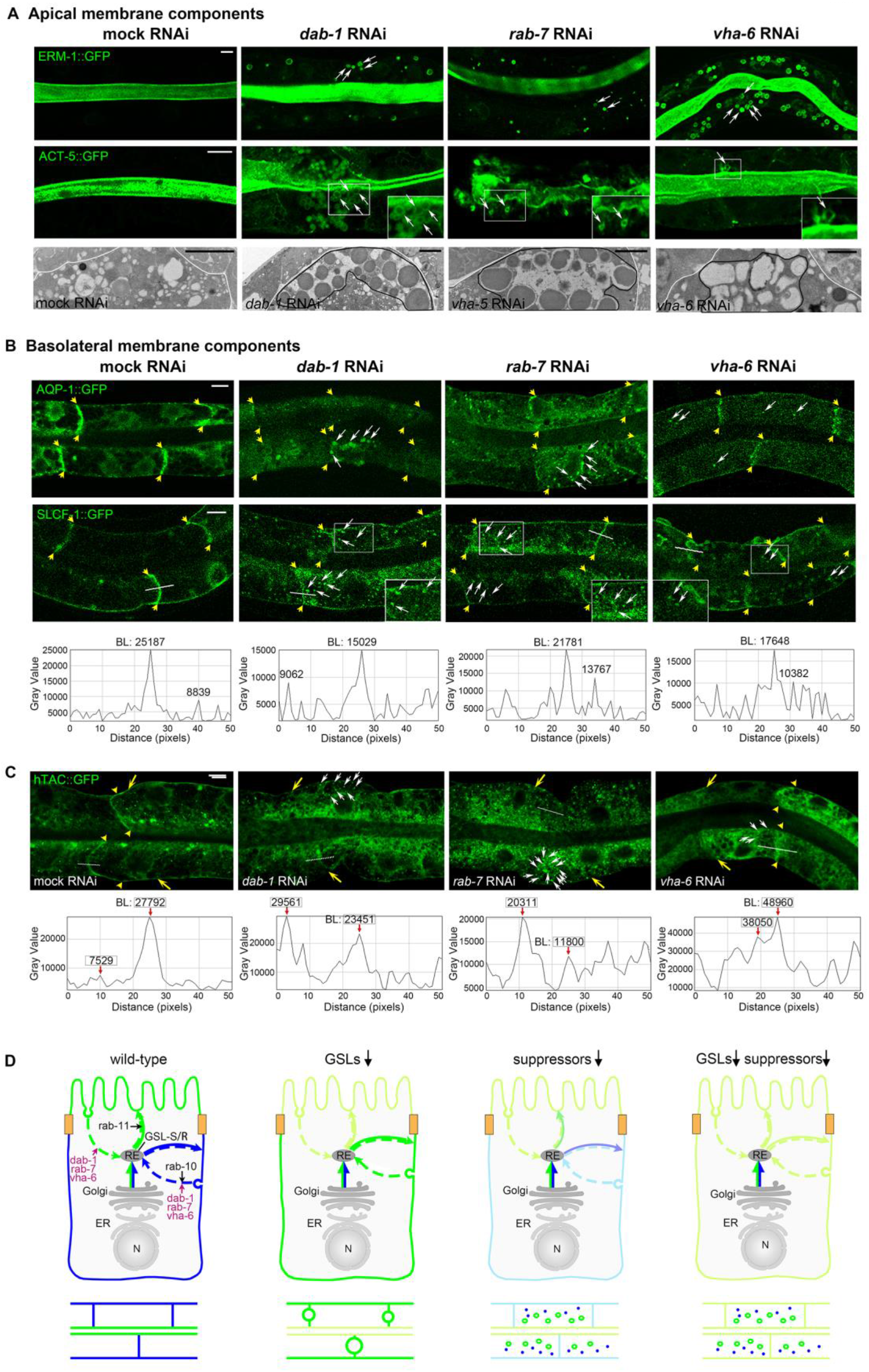
DAB-1, RAB-7 and VHA-6 are required for apical and basolateral membrane polarization but not positioning and function on polarized endocytic-recycling routes. Model for suppression. **(A)** *dab-1-rab-7 and vha-6* RNAi displaces ERM-1 and ACT-5/actin to cytoplasmic vacuoles (a few indicated by white arrows) but not to basolateral (BL) membranes in larval INT cells during polarized membrane biogenesis. Vacuolar aggregates outlined in black in TEM images below (1 cell/image shown, cell boundary outlined; see **Fig.S7A** for whole INTs). No vacuoles with inward pointing microvilli (=ectopic lumens/ELs) were found in serial sections (see text and **Fig.S7A** legend). ERM-1 tracking and TEM analysis were performed in a sensitized RNAi (*rrf-3*) background (Methods). **(B)** The BL integral membrane components AQP-1/aquaporin and SLCF-1 are lost from *dab-1-rab-7 and vha-6(RNAi)* membranes of expanding larval INT cells (lateral portion of BL membranes bracketed by yellow arrows) and displaced to cytoplasmic vesicles/tubulovesicles (a few indicated by white arrows; note difference to ERM-1+ vacuoles in **(A)**) but are not mislocalized to apical membranes. Fluorescence intensity across SLFC-1+ membranes (white line) shown beneath, with highest cytoplasmic intensity point indicated (BL = basolateral membrane; Methods; note that only few membranes retain enough SLCF-1 to qualify for quantification). **Fig.S7C** for quantification of membrane/cytoplasm GFP ratio. **(C)** The recycling marker hTAC is lost from BL membranes in *dab-1-, rab-7-,* and *vha-6(RNAi)* larval INT cells (yellow arrows; arrowheads bracket lateral portions) and displaced to vesicles (a few indicated by white arrows). Fluorescence intensity across hTAC+ membranes shown beneath (see **B** for legend and acronyms; Methods; note that only few membranes retain sufficient hTAC to qualify for quantification). **Fig.S7D** for quantification of membrane/cytoplasm GFP ratio. Thin confocal sections of 2 pairs of opposing cells of *dab-1-, rab-7-,* and *vha-6(RNAi)* larval INTs are shown throughout. Scale bars: 5μm; 1μm in TEM images. **(D)** Model for suppression. Reduction of flux from polarized endocytic-recycling routes into the GSL-dependent apical secretory route reduces apical-to-basolateral misrouting upon GSL depletion and hence suppresses polarity conversion. Wild-type-, single- and double GSL/suppressor depletion conditions are shown in single cells, corresponding tissue phenotypes beneath. Direction of delivery of polarized membrane components (arrows) and putative location of suppressors and of RAB-10 and RAB-11 on corresponding delivery routes are indicated. GSL-dependent apical-cargo-sorting or apical-vesicle-routing site (GSL-S/R) is arbitrarily placed at the recycling endosome (RE). Secretory routes: solid lines, recycling routes: dashed lines. Reduced cargo flux to (or arrival at) plasma membranes indicated by lighter colors. Mild suppression is shown, with membrane light-green rather than blue in far-right cartoon (indicating remaining displaced apical cargo). Only post-Golgi trafficking routes and only changes in direction and quantity of apicobasal cargo delivery are indicated. Cartoons of polarized cells/colors (apical green/basolateral blue) as in **Fig.S1** and throughout. N: nucleus. The itinerary of a post-Golgi GSL-directed secretory apical vesicle trajectory (direct, indirect, transcytotic) and the location of GSL-S/R sites can be in part deduced from routes on which suppressors function (compare Fig. 6A,E). The apical trajectory could be: direct (route 1; Fig. 6A) if suppressors function on pre-Golgi-/Golgi-directed retrograde recycling routes (GSL- S/R at the Golgi or beyond); indirect (route 2; Fig. 6A) if suppressors function on endocytic-recycling routes (GSL-S/R at the RE [shown] or other, e.g. apical, sorting endosomes). A transcytotic route (route 4; Fig. 6A) is less likely, as it would implicate a suppressor function on secretory routes and GSL-S/R sites at, or beyond, the basolateral membrane (see Fig. 6A legend).

To assess whether the suppressors’ function in polarized membrane biogenesis was limited to the apical membrane, basolateral membrane expansion was monitored under the same conditions. *dab-1-, rab-7-* and *vha-6* RNAi also displaced the sugar transporter SLCF-1 and the water channel AQP-1 from basolateral membranes of expanding intestines, retaining them on small vesicles/tubulovesicles, but did not misdirect them to the opposing (apical) domain (Figs.7B, S7C).

We conclude that DAB-1, RAB-7, and VHA-6, each required for apicobasal polarity conversion in GSL-depleted intestines, are also required for apical and basolateral membrane polarization, but not apicobasal membrane polarity, in wild-type intestines. Their dual roles in apical and basolateral membrane biogenesis suggested that these trafficking molecules operate upstream of a GSL-dependent apical vesicle trajectory, on either: (1) secretory routes, moving unsorted apicobasal cargo towards GSL-dependent sorting/routing sites, as part of, or parallel to, the GSL-directed trajectory (Fig. 6E, route 1); or (2) interfacing endocytic-recycling routes, moving both apical and basolateral cargo towards these sites (Fig. 6E, routes 2).

### The suppressors of apicobasal polarity conversion uncover polarized recycling routes that replenish biosynthetic-secretory routes to manufacture new polarized membrane

To distinguish between these possibilities, we examined if DAB-1, RAB-7 and VHA-6 function in recycling or secretion during wild-type *de novo* polarized membrane biogenesis and monitored basolateral trafficking routes, well-characterized in mature intestines (*48*), in expanding larval intestinal cells. During membrane expansion, *dab-1-, rab-7-* and *vha-6* RNAi: (1) retained the recycling marker hTAC::GFP on cytoplasmic vesicles (Fig. 7C, S7D); (2) failed to accumulate Yolk, a marker for secretion (Fig.S8A; see legend for Yolk assay); and (3) did not abort membrane delivery of TGN-38(Y314A), a Golgi resident trapped at the membrane due to a recycling defect (Fig.S8B).(*49*) We conclude that the 3 GSL suppressors operate in recycling, not secretion, during net membrane expansion in wild-type intestines.

To determine if the 3 GSL suppressors could position polarized vesicles, we examined RAB-11+ and RAB-10+ recycling endosomes, two of few vesicle components with documented (albeit not exclusive) functions on apical (predominantly anterograde) and basolateral (predominantly retrograde) routes, respectively (Fig. 8A-B) (*48*). Tier-2 interaction screens had identified RAB-11 as a loss-of-function enhancer of the GSL-dependent polarity conversion (Fig. 3D), suggesting RAB-11 acts in parallel to, or downstream of, GSLs on an apical biosynthetic trajectory that might pass through the recycling compartment (Fig. 6A, route 2; 6E, route 3). If so, the 3 suppressors, expected to promote recycling upstream of GSLs, would be expected to function via RAB-10+, but not RAB-11+, recycling carriers during net membrane expansion. Indeed, in *dab-1-, rab-7-* and *vha-6(RNAi)* larval intestinal cells, GFP::RAB-11+ vesicles were mostly retained at apical, while GFP::RAB-10+ vesicles were lost from basolateral, membranes and aggregated in the cytoplasm (Fig. 8A-B). These results are consistent with a function of DAB-1, RAB-7, and VHA-6 on retrograde arms of polarized recycling routes that operate during *de novo* polarized membrane biogenesis.

**Figure 8.**
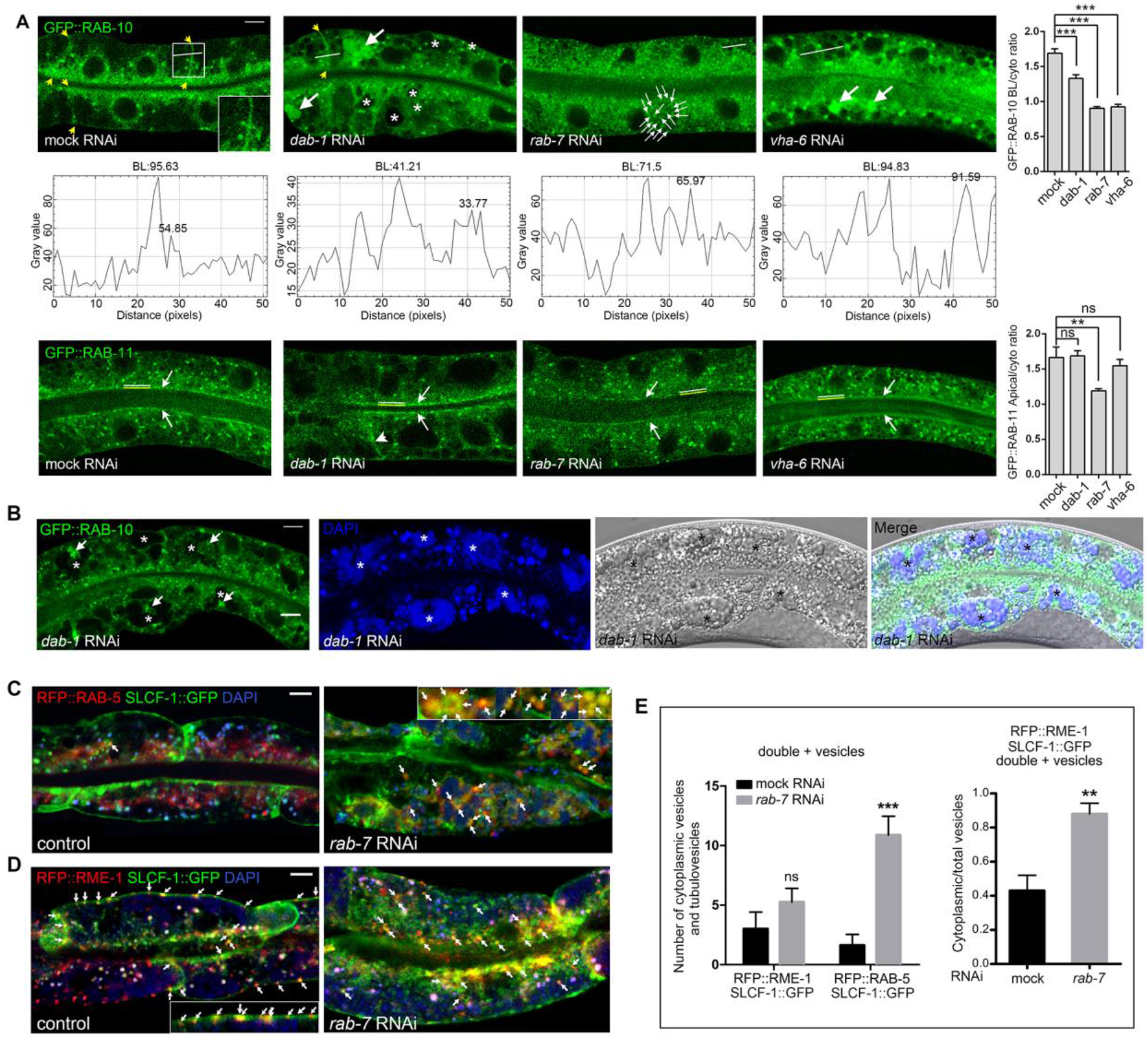
The GSL suppressors recycle polarized membrane components during *de novo* polarized membrane biogenesis in wild-type. **(A)** RAB-10+ vesicles are displaced from basolateral (BL) membranes (lateral portion bracketed by yellow arrows) and retained in cytoplasmic aggregates (*dab-1* RNAi), clusters of small vesicles (*rab-7* RNAi) and enlarged vesicles (*vha-6* RNAi; examples of each indicated by white arrows) in expanding larval *dab-1-, rab-7-* and *vha-6(RNAi)* INT cells. \**dab-1(RNAi)* specific vacuoles can contain RAB-10+ material (see **B**). Fluorescence intensity across RAB-10 enriched BL membranes (white line) shown beneath with highest cytoplasmic intensity point indicated (Methods; note that only few membranes retain enough RAB-10 to qualify for quantification). RAB-11+ vesicles largely maintain apical enrichment (white arrows; note some loss in *dab-1-* and *rab-7-,* and occasional BL displacement in *dab-1(RNAi)* cells [arrowhead]). Bar graphs: quantification of fluorescence intensity ratio of membrane-associated to cytoplasmic vesicles. **(B)** GFP::RAB-10+ material accumulates in *dab-1(RNAi*) specific DAPI+ vacuoles (*; compare SFig.6F). White arrows point to GFP::RAB-10+ material inside such vacuoles (asterisk). **(C, D)** The integral membrane component SLCF-1 traverses endocytic-recycling carriers during wild-type polarized membrane biogenesis, a process requiring RAB-7. Wild-type larval INT cells (controls, left) reveal occasional SLCF-1 passage through RAB-5+ presumed membrane-derived, and RME-1+ presumed membrane-directed, arms of endosomal-recycling routes (arrows). Note close association of RFP::RME-1+; SLCF-1::GFP+ vesicles with membrane-attached tubules (inset in **D**). *rab-7* RNAi (right) increases cytoplasmic SLCF-1::GFP+; RFP::RAB-5+ (inset in **C**), but not SLCF-1::GFP+; RFP::RME-1+, endosomal vesicles, although it broadens membrane-associated SLCF-1::GFP+; RFP::RME-1+ tubulovesicles. Also note loss of BL SLCF-1 and enlarged LROs (blue), demonstrating *rab-7(RNAi)* effect. Arrows point to GFP+ RFP+ DAPI-vesicles/tubulovesicles (compare **Fig.S8C, D,** Methods). **(E)** Quantification of SLCF-1::GFP+; RFP::RME-1+ and SLCF-1::GFP+; RFP::RAB-5+ cytoplasmic vesicles/tubulovesicles, and ratio of SLCF-1::GFP+; RFP::RME-1+ cytoplasmic to total (cytoplasmic plus membranous) vesicles/tubulovesicles. Mean ± SEM shown, n=3/N=8. Note increase of cytoplasmic versus tubular RFP::RME-1+; SLCF-1::GFP+ vesicles in *rab-7(RNAi)* intestinal cells, consistent with SLCF-1’s slowed progression through the recycling compartment. This could suggest additional RAB-7 functions on membrane-directed rotes during *de novo* polarized membrane biogenesis that might be independent of its GEF SAND-1 (SAND-1 fails to suppress polarity conversion; **Fig.S6C**). Thin confocal sections of 2 pairs of opposing cells of expanding larval INTs are shown throughout, corresponding Nomarski and Nomarski overlay images in B. Scale bars: 5μm.

Collectively, these findings suggested that the GSL suppressors might return previously polarized membrane components to the GSL-dependent biosynthetic route. This scenario would predict that, during wild-type polarized membrane addition, integral membrane components traverse endocytic-recycling carriers in which they become trapped in the absence of the GSL suppressors. Recycling of polarized cargo is thought to increase the fidelity of sorting rather than supplement *de novo* polarized membrane biogenesis, but the delivery of polarized membrane components to or from the expanding membrane has not yet been visually tracked. We used double-labeling of polarized cargo-carrier pairs to assess this process. The integral membrane component SLCF-1 could indeed be detected in RAB-5+ (presumed membrane-derived) and RME-1+ (presumed membrane-directed) endocytic-recycling carriers (*48*) during net membrane expansion in wild-type larval intestinal cells (Fig. 8C-E, 8SB-D). Moreover, depleting RAB-7 increased RFP::RAB-5; SLCF-1::GFP-but not RFP::RME-1; SLCF-1::GFP double-positive vesicles in these cells, indicating that RAB-7 depletion trapped SLCF-1 on endocytic arms of polarized recycling routes (Figs. 8C-E, 8SB-E). RAB-7 depletion also increased the cytoplasm-to-plasma-membrane-associated ratio of RFP::RME-1; SLCF-1::GFP double-positive vesicles, suggesting that RAB-7’s function downstream of RAB-5 may not be limited to vesicle degradative routes at the time of *de novo* membrane biogenesis (Fig. 8C-E; legends for details).

In summary, the suppressor analysis identified polarized recycling routes that promote the biogenesis of new apicobasal membrane, with novel functions on such routes for DAB-1, RAB-7, VHA-6 and 4 additional candidate pathway components (ARF-6, RAB-6.2, DPY-23, ARL-8). While dispensable for regulating wild-type membrane polarity, these trafficking molecules interact with GSLs in polarity via their canonical roles in endocytic-recycling that here serve to supply the GSL-directed biosynthetic route with recycled polarized membrane components. Reducing influx into this anterograde secretory vesicle trajectory reduces basolateral misrouting of apical membrane components upon GSL depletion and hence suppresses apicobasal polarity conversion (Fig. 7D). We conclude that polarized membrane components are directly recycled back to manufacture new membrane in the expanding *C. elegans* intestine, revealing polarized recycling as an integral part of *de novo* polarized membrane biogenesis.

## DISCUSSION

Here, we identify the biosynthetic-secretory (secretory) pathway, supplemented by polarized recycling routes, as a key determinant of apicobasal membrane polarity in the *C. elegans* intestine. We find that the directionality of this essential trafficking pathway is regulated upstream of PARs and independent of polarized target domains to asymmetrically insert apical membrane components into the growing epithelial membrane and thereby polarize it. We identify multiple components of secretory carriers, located throughout the pre- and post-Golgi endomembrane system, that function as directional (apical) cues during net membrane expansion, whether or not this membrane is polarized. Together, these findings suggest that, during *de novo* polarized membrane biogenesis, this anterograde vesicle trajectory, required to supply all sides of the plasma membrane, is transiently routed to a nascent apical domain that it concomitantly expands, polarizes, and positions. This alternative mode of membrane polarization shares features with early polarity models that had envisioned the fixed (apical or basolateral) directionality of bulk membrane delivery rather than membrane-based polarity cues to polarize the epithelial membrane (*8, 9*). It could also offer solutions to unresolved difficulties of current models of polarity and polarized trafficking (Fig. 9; Introduction and below).

**Figure 9.**
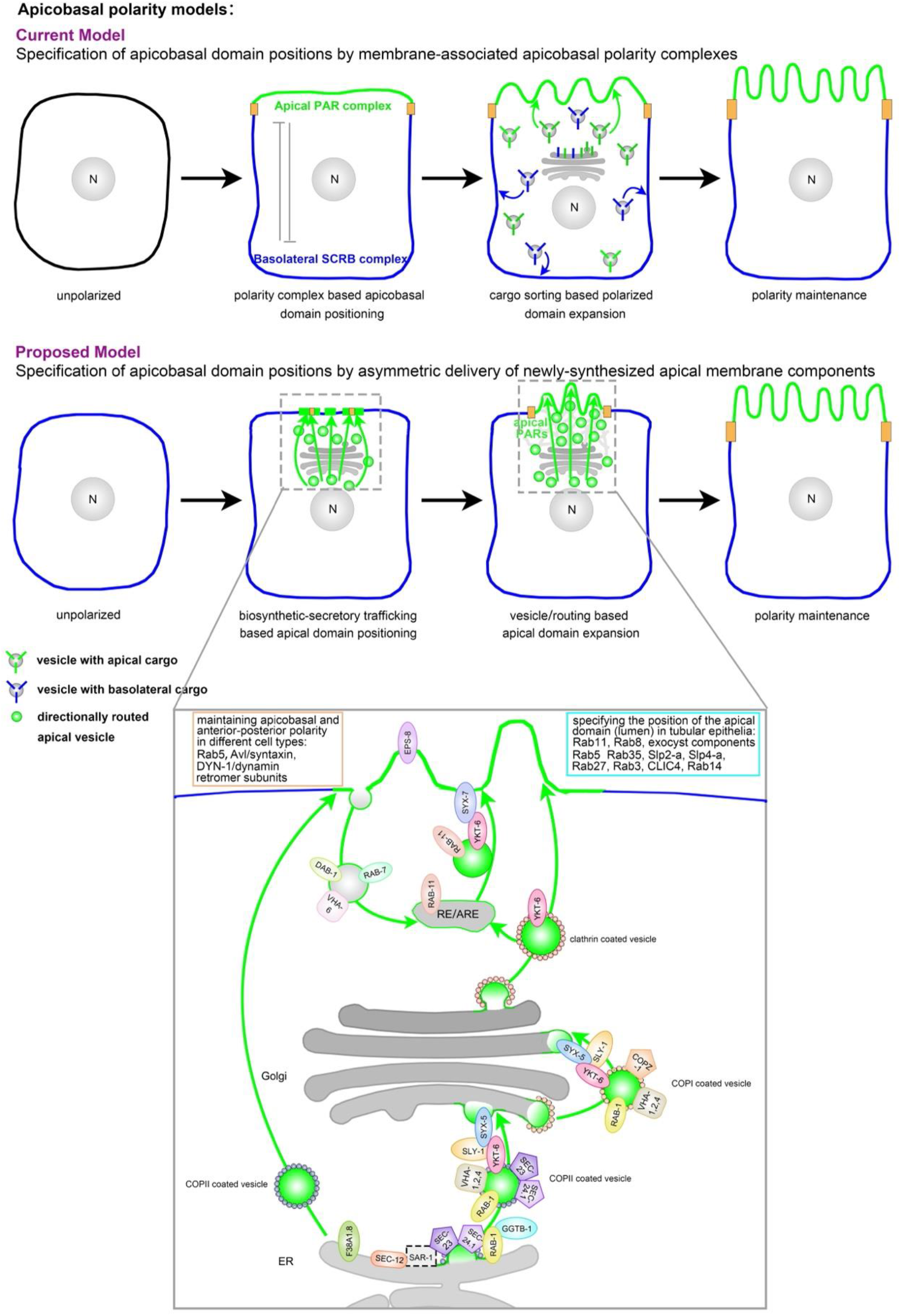
Current and proposed apicobasal polarity models. Schematics of plasma membrane polarization (left to right). Cartoons of cells and colors (apical green/basolateral blue) as in **Fig.S1** and throughout. Polarized membrane domains, polarity cues, endomembranes, cargo, and trafficking directions are indicated; boxed areas in second row are magnified beneath (see **Figs.1A, S1A, S9** for 3D tissue context). In the proposed model (row 2), the biosynthetic-secretory pathway is routed to the nascent apical domain by transient cytoskeletal structures (indicated as grey web in cell 3). In this model, the positioning of membrane-based apical polarity cues and junctions/spot junctions (orange) depends on the asymmetric delivery of apical membrane components, while apical junctions subsequently or concomitantly assemble laterally to secure the expanding apical domain (see text). This notion (secondary lateral junction assembly) is supported by earlier appearing apical spot junctions during wild-type *C. elegans* intestinal polarity establishment (*85*) and by the *de novo* assembly of junctions around ectopic basolateral lumens in GSL or clathrin/AP-1 depleted intestines (*7, 10, 11*). The mutual inhibition of apicobasal membrane- and junction-based polarity cues (PAR: partitioning defective; SCRB: Scribble) is indicated by inhibitory arrows in cell 2 in row 1 (current polarity model) to emphasize its role in initiating membrane polarity in this model. It is, however, assumed to be an ongoing process during polarity maintenance in both models (see text). Membrane-/junction- and trafficking-based polarity cues could furthermore reciprocally depend on each other, as suggested by effects of apical PAR complex components on the regulation of polarized trafficking (*68, 86*). Organelles, polarity cues and all trafficking routes are simplified for clarity. N: nucleus; Golgi: grey structure above nucleus. The magnified image depicts a unifying model of one single route for a biosynthetic vesicle with apical destination that traverses the recycling compartment. Possible additional or alternate Golgi bypass routes and direct routes from the Golgi to the apical domain are also indicated. Vesicle cages (COPI/II and clathrin) could provide the physical connection to the guiding cytoskeleton (not shown). Note that most of the here identified secretory trafficking molecules are components of any of these 3 known vesicle coats and/or the machinery that assembles them. Also note, that each of them is required to prevent basolateral deviation of apical cargo delivery during membrane polarization. RE/ARE: recycling-/apical recycling endosome. Boxes on top list components of apical endocytic and transcytotic recycling routes and of the apical membrane/vesicle interface that have been identified by their requirement for maintaining apicobasal and anterior-posterior polarity in different cell types (Rab5 (*14*); Avl/syntaxin (*14*); DYN-1/dynamin (*15*); retromer subunits (*16–18*)) and for positioning the apical domain (lumen) in tubular epithelia in different tissues and species (Rab11, Rab8, exocyst components (*19*); Rab5, Rab8, Rab11 (*33*); Rab35 (*20*); Slp2-a, Slp4-a, Rab27, Rab3 (*21*); CLIC4 (*22*); Rab14 (*23–26*)).

An upstream polarity function of intracellular vesicular trafficking, regulated to assume apical directionality but not intrinsically polarized, addresses the current polarity model’s paradox that the canonical apicobasal polarity cues and junctions must themselves be polarized to membrane domains whose positions they ought to specify. These membrane-based apicobasal polarity cues’ well-characterized signaling network of mutual inhibition (*1, 2*), along with the assembly of apical junctions, would, conversely, be well placed to secure the boundaries of a nascent apical domain that is expanded into the growing membrane by the secretory pathway, as suggested here (Fig. 9). A flexible, regulated directionality of the secretory pathway also addresses difficulties of sorting-based models of polarized trafficking that place the directional control with polarized cargo signals and their carrier-based decoders (*1, 4*). For example, cargo- or carrier-based signals fail to consistently sort in one direction (e.g., clathrin/AP-1 sort to both apical and basolateral domains, a problem unresolved by cell-type-specific adaptors such as AP-1B) (*50–53*). It is also still unclear how sorting confers long-range apicobasal directionality to vesicles and whether and where these vesicles travel: (a) randomly (e.g. captured only at their polarized target domain by cognate v-/t-SNARE- or coincident phosphoinositide pairs) (*54, 55*); (b) guided (e.g. by cytoskeletal tracks/motors) (*56, 57*); or (c) by endomembrane maturation (e.g. via Rab cascades) (*58*). Our findings suggest that vesicle-associated, e.g., cytoskeletal, spatially and temporally regulated guidance cues determine trafficking’s directionality during epithelial membrane polarization and that such cues can override apicobasal sorting signals during net polarized membrane expansion.

There is precedence for a regulated change of the secretory pathway’s directionality in non-epithelial cells (*59–61*). In budding yeast, this pathway is actin-dependently oriented to the bud site during polarized bud growth to direct a full set of non-polarized membrane components to the daughter cell (*60*). Membrane-based landmarks other than already polarized membrane domains could similarly orient such trafficking routes in epithelial cells (e.g., the midbody) (*62, 63*). In epithelial cells, however, the directional routing of polarized (e.g., apical) membrane components - proteins and lipids (e.g., GSLs) - will asymmetrically expand and thereby polarize the membrane. Indeed, membrane lipid measurements in polarized versus non-polarized MDCK cells reveal the net addition of apical lipids to the resident complement of membrane lipids (*64*). Therefore, a transient increase in apical cargo/carrier components (e.g., via an increase in GSL biosynthesis) or the transient formation of apical cytoskeletal tracks, could regulate and even initiate membrane polarization. Consistent with this proposition, the transcriptionally regulated biosynthesis of sterols (raft components - like GSLs) (*4, 5*) was shown to override canonical polarity cues and initiate polarity in fission yeast (*65*).

A model of membrane polarization via the vesicle-based directional insertion of apical membrane components can also serve as a morphogenetic model that consolidates inTER- and inTRA-cellular apical domain and lumen biogenesis (Fig.S9). Notably, inTRAcellular lumenogenesis models propose the expansion and polarization of the apical/lumenal membrane by directional vesicle coalescence (*66–68*). If such a process of net apical domain extension within a single cell (e.g., inside the junction-less unicellular *C. elegans* canal) proceeds similarly between pairs of cells (e.g., in the multicellular *C. elegans* intestine), it will position the apical, and consequently the basolateral, domain (Fig. S9). Consistent with this concept, all trafficking components that we found to regulate polarity in the intestine also direct apical membrane expansion in the canal (EPS-8, with a likely intestine-specific function, is the only exception). We also note that the same trafficking molecules were identified in various independent studies by either a function in the polarity of multicellular, or by a function in the expansion of unicellular, tubular epithelia (e.g. Rab35, Rab5, exocyst components, Slp/BTZ(bitesize), CLIC4/EXC-4) (*14, 19-22, 67, 69-74*).

Secretory routes can intersect with recycling routes on their way to the apical membrane (Fig. 6A) (*75, 76*). A biosynthetic-secretory vesicle trajectory, routed from the ER to the nascent apical domain during membrane polarization, could thus also accommodate anterograde arms of apical endocytic- and transcytotic recycling routes. Components of such routes, more amenable to loss-of-function studies than the essential secretory pathway, continue to be identified by their roles in apicobasal and anterior-posterior polarity (itemized in Fig. 9). All components of secretory and recycling carriers that were identified by their ability to position the apical domain could be required to regulate the directionality of a single common apical vesicle trajectory, e.g., by connecting it to a dynamic cytoskeleton at consecutive steps towards the plasma membrane (Fig. 9). Routing of such a trajectory from the ER to the apical membrane might require components of: (1) pre-Golgi secretory carriers (this study); (2) Golgi-/post-Golgi endomembranes/carriers (GSLs, clathrin/AP-1; prior studies by us and others) (*7, 10, 11, 50-53, 77*); (3) apical recycling carriers and the apical-vesicle/membrane interface (e.g. RAB-11(*19*), V-ATPases (*38*), EPS-8 (*40*); this study and other studies; Fig. 9). The concept of one common trajectory is supported by the close genetic interaction that was here observed between all 3 groups of vesicle-based polarity cues (pre-Golgi, post-Golgi/sorting, post-Golgi/recycling), where the combined depletion of any two components, mild enough to maintain polarity when only one is depleted, suffices to synthetically induce apicobasal polarity conversion.

It has been noted that secretory pathway components are conspicuously absent from the collection of trafficking molecules identified in numerous genetic screens on apicobasal polarity (*18*). This may be due, in part, to the narrow range of interference conditions that permit separation of the pathway’s directional from its general secretory function. In contrast, screens on apical secretion and apical membrane expansion -- processes more robust to pathway disruption -- in both flat and tubular, in particular unicellular tubular, epithelia have repeatedly identified early pathway components (e.g. COPI/II subunits).(*24, 78–82*) However, tracking cargo delivery beyond the membrane (assessing secretion) must miss, while tracking cargo accumulation in the cytoplasm (assessing membrane delivery) can mask, changes in the position (apical-versus-basolateral) of cargo at the membrane – which would explain these screens’ failure to identify the pathway’s regulatory function in apicobasal polarity. Secretory molecules were also identified in screens tracking the delivery of single apical cargo, -carrier or -junction components (Tab.S2), an approach likewise unable to identify the pathway’s regulatory polarity function. However, these screens provide additional confirmation for these secretory molecules’ direct role in apical cargo delivery, here largely deduced from the recruitment of the apical domain identity marker ERM-1 (incidentally, membrane polarity defects were occasionally observed in these screens, but set aside or dismissed as non-specific).

While anterograde secretory routes can intersect with recycling routes on their way to the membrane, retrograde recycling routes can intersect with pre- and post-Golgi secretory routes, the former to replenish pre-Golgi intermediates, the latter to increase the fidelity of cargo sorting (*83*). Here, we identify endocytic-recycling routes that redirect polarized membrane components to the biosynthetic-secretory pathway during net membrane expansion, suggesting these membrane components are directly re-utilized to manufacture new membrane. This unexpectedly close interface of polarized secretion and recycling during *de novo* membrane biogenesis could explain the astonishing fluidity of membrane polarity that we observe in the fixed tissue context of the *C. elegans* intestine where lumen positions can shift *in situ* between apical and basolateral sides of single postmitotic cells. Such a dynamic aspect of single cell polarity may be conserved through phylogeny, as indicated by the conservation - to the molecular level - of the trafficking-dependent apicobasal polarity phenotype from worms to humans (*84*). For example, polarity conversion in the *C. elegans* intestine mirrors polarity inversion in MDCK cysts, where the transition to 3D culture conditions, conversely, shifts the apical domain from the future basolateral to its new apical location (Fig.S10) (*19*). These distinct *in vivo* and *in vitro* models may thus provide complementary vistas on the same trafficking-directed process of membrane polarization, with apical recycling molecules such as RAB-11/Rab11 (identified in both) marking shared components of one common secretory vesicle trajectory that traverses the recycling compartment on its way to the nascent apical domain. A developmentally regulated, directionally flexible biosynthetic-secretory pathway, in close apposition to polarized recycling routes that permit exchange between all membrane domains, would provide greater plasticity of *in vivo* single cell polarity than currently appreciated.

## Acknowledgments

Strains and reagents were kindly provided by Barth D. Grant (Rutgers University, Piscataway, New Jersey, USA), Kenneth Kemphues (Cornell University, Ithaca, New York, USA), Christian E. Rocheleau (McGill University, Montreal, Quebec, Canada), Jonathan Pettitt (Institute of Medical Sciences, Foresterhill, Aberdeen, UK), Keith Nehrke (University of Rochester Medical Center, Rochester, New York, USA) and the Caenorhabditis Genetics Center (CGC). TEM work was performed at the Center for Systems Biology/Program in Membrane Biology (Massachusetts General Hospital/MGH, Boston, USA). We thank David Hall (Albert-Einstein College of Medicine, New York, USA) for discussion of ultrastructural images, John Fleming and Frank Solomon for critical reading of the manuscript, and Howard Weinstein and Ronald Kleinman (MGF*f*C, Harvard Medical School, Boston, USA) for ongoing support.

## Funding

NIH Office of Research Infrastructure Programs P40 OD010440 (CGC)

IBD and BADERC grants DK43351 and DK57521 (TEM/MGH)

NIH GM078653, MGH IS 224570 and SAA 223809 (VG)

## Supplementary Materials

Materials and Methods

Figs. S1 to S9

Tables S1 to S4

References 84-93

Movie S1

