## Supplemental text and Figures for "The biosynthetic-secretory pathway, supplemented by recycling routes, specifies epithelial membrane polarity"

**This PDF file includes:**

Materials and Methods
Figs. S1 to S9
Tables S1 to S4
Caption for Movie S1

**Other Supplementary Materials for this manuscript include the following:**

Movie S1

#### Materials and Methods

##### Experimental model and subject details

An introduction to the *in vivo* analysis of polarized membrane biogenesis in the *C. elegans* intestine and excretory canal, with an extended Methods section and a visual demonstration of the techniques used in this study is provided in (*87*, *88*).

##### *C. elegans* Strains, Culture Conditions, Genetics

Wild type (N2 Bristol) and mutant *C. elegans* strains were cultured and genetic crosses performed using standard methods (*89*). Worms were generally maintained at 20-22$^{o}$C on Nematode Growth Medium (NGM) plates seeded with *E. Coli* OP50 (*90*). The list of strains used in this study is provided in **Tab.S4**.

##### RNA interference/loss-of-function analysis

RNAi was carried out by feeding worms *E. coli* HT115 (DE3), producing double stranded/ds RNA of the gene of interest, as previously described (*7*, *91*). For standard RNAi, bacterial feeding clones were inoculated from LB plates into 1ml LB liquid medium containing 50μg/ml ampicillin and incubated for 8-18 hours at 37℃. 200μl cultured RNAi bacteria were seeded onto agar plates supplemented with 2mM IPTG and 25μg/ml carbenicillin. dsRNA was induced at room temperature for at least 6 hours before picking 4 - 6 L4 larvae onto each RNAi plate. Most bacterial clones were derived from the Ahringer genome-wide RNAi feeding library (J. Ahringer, Welcome Trust/Cancer-Research-UK-Gurdon-Institute, Cambridge, UK). The integrity of all RNAi clones was verified by sequencing (discrepancies to reported sequence information are noted in **Tab.S1**, primers to generate RNAi vectors are available on request.

RNAi was determined as the method of choice for the loss-of-function analysis of the chiefly sterile, maternal and embryonic/early-larval lethal intestinal and canal tubulogenesis genes since it can be easily titrated to produce a range of different degrees of severity and since it effectively targets maternal RNA (limiting the use of germline mutants due to the requirement of balancers that provide maternal product). For the tubulogenesis screens (see below), moderately-severe (standard) RNAi conditions were used to generate a broad spectrum of informative reduction-of-function phenotypes. For the analysis of early secretory pathway components, any approach inducing severe reduction- or full loss-of-function phenotypes was not informative due to its disruption of all membrane-directed traffic (see text and below; excluding strong tissue-specific protein targeting approaches). For these genes, mild-to-moderate RNAi conditions were empirically determined for any given gene by: modulating IPTG concentrations in RNAi plates; diluting the RNAi clone of interest with different amounts of mock bacteria (no vector or vector with irrelevant gene); varying the developmental stage (L2 to adult) of the parental strain in which RNAi was induced; using the RNAi-sensitive strain *rrf-3(pk1426).(87, 88)*.

Conditions were furthermore varied to achieve interference at different time points during embryonic and/or later stages of development, e.g. RNAi was induced: in parents (evaluating the F1 progeny; standard parental RNAi); in larvae (evaluating the same generation; conditional larval RNAi); or in adults. Conditional larval RNAi was carried out by bleaching 30-50 gravid adults in one drop bleaching solution (a 1:4 mix of 10M NaOH and household sodium hypochlorite) on the edge of an RNAi plate and allowing hatched larvae to crawl to the bacterial lawn (see (*88*) for details). Appropriate controls were added to ensure that RNAi was effective when induced at later time points during development or in adults (e.g. by using *gfp* RNAi on a GFP expressing strain).

**Tubulogenesis screens and selection of trafficking molecules**
Screens I and II (genome-wide RNAi-based intestinal and excretory canal tubulogenesis screens) first examined all chromosome III genes. This pilot screen revealed that >90% of all genes whose loss induced informative tubulogenesis phenotypes in ERM-1::GFP-labeled intestinal and excretory canal epithelia were lethal, consistent with the essential function of these organs whose structural defects or functional failure typically result in L1-larval lethality. The remaining screens were then restricted to all lethal genes in the genome, with the expectation of achieving >90% whole genome coverage. Mild- to moderately-severe RNAi conditions (standard RNAi, see above) were used, generating a broad spectrum of phenotypes. All animals were examined throughout development under a dissecting microscope with high-power fluorescence attachment, and a selection was additionally assessed by confocal microscopy (see below, microscopy). Emphasis was placed on the analysis of embryonic/early larval development (7 genes identified from this screen have been previously characterized; see text) (*7, 11*). For the ERM-1[++] cystic canal modifier screen, extended classes of genes experimentally demonstrated or predicted to interact with ERM-1/ezrin-radixin-moesin in any species were evaluated for enhancement or suppression (4 genes identified from this screen have been previously characterized) (*31*).

From a broad range of different intestinal and canal morphogenesis phenotypes that were identified in one or several of these screens, only those with defects in apical/lumenal membrane expansion, polarization and positioning (defined by type of ERM-1 displacement; see text) were collected and the identities of the underlying defective genes unblinded. All those genes were selected for further analysis, whose products were implicated in vesicular trafficking in any species based on published experimental data, search engines (such as AceView [http://www.ncbi.nlm.nih.gov/ieb/research/acembly/], Ihop [http://www.ihop-net.org/UniPub/iHOP/]; the Yeast genome database [http://www.yeastgenome.org], Flybase [http://www.flybase.org]; the Mouse Genome Database [http://www.informatics.jax.org] and the Human Genome Database [http://www.gdb.org]) and/or protein blast searches (*87*). All proteins with documented or predicted trafficking function that were identified are highly conserved between species, most from yeast to human, and 49/50 have close human orthologs. Proteins were further classified according to structure and trafficking function and annotated with regard to their yeast and human orthologs, cellular and subcellular localization and, where available, presumed trafficking route, with specific attention to any described function in *C. elegans* (collected in **Tab.S1**).

**Phenotype analysis**
All phenotypes were evaluated in multiple independent experiments (n>5, with N>500 animals/plate), performed in duplicates or triplicates, and animals were characterized alongside a positive control (e.g., *let-767* RNAi for the larval intestinal polarity phenotype and *chc-1* RNAi for the embryonic intestinal polarity phenotype, see text) and a negative control (empty vector or

unrelated gene, typically a larval or embryonic lethal gene [itemized in **Tab.S1**; negative results of the initial tier-1 screens (**Fig.1B**) serve as additionally negative controls). The membrane-level resolution of the visual analysis, tracking fluorescently-labeled apical intestinal and canal membranes or single-cell canal tubules (tier-1/III canal screen), permitted the distinction of specific phenotypes (classes 1 – 10) in a background of moderate-to-severe body morphogenesis defects induced by the loss of the mostly essential genes. For example, the specificity of the intestinal polarity phenotype is emphasized by its low incidence in the tier-1/I intestinal screen that included all lethal genes (N=18/4978 knockdowns). RNAi phenotypes of representative genes of a targeted pathway were confirmed via phenocopy by: germline mutants; RNAi with subunit or pathway components of the targeted gene; dose-dependency; and/or independent studies (see text). Cell-autonomous functions of selected intestinal genes were confirmed by phenocopy in an intestine-specific RNAi strain (*rde-1* mutant background supplemented with an intestine-directed *rde-1* transgene) *(87, 88).*

For the standard analysis of RNAi phenotypes, the progeny of two RNAi plates seeded with 4 L4 stage *erm-1p::erm-1::gfp; rol-6p::rol-6(su1006)* larvae was evaluated (see below, fluorescent fusion proteins). The dominant marker *rol-6*, conferring a circumferential rolling movement, facilitates the 3D (apicobasolateral) evaluation of polarized tubulogenesis. RNAi conditions were empirically determined for each gene (as above), with the goals; (1) to extend development from early-embryonic intestinal and canal tubulogenesis as far as possible through larval development, in order to track polarized membrane biogenesis throughout the entire postmitotic growth phase (see text and **Fig.S1**); and (2) to examine a broad spectrum of mild-to-severe loss-of-function phenotypes of each gene (all trafficking genes examined here generate dose-dependent apical membrane polarization, positioning or expansion loss-of-function defects; **Fig.S2**). The mildest RNAi conditions were used for the analysis of components of the early secretory pathway, to prevent disruption of membrane-directed trafficking, a prerequisite for the analysis of its effect on membrane polarity (see text). All animals were evaluated throughout development by high power dissecting and/or confocal microscopy (see below, microscopy), with most phenotypes present in embryos and/or early larvae. For the initial assessment of lethality (**Tab.S1**), the F1 progeny (both embryos and larvae) was counted on day 3. The morphology of embryos was scored by dissecting microscope and confirmed at higher magnification by Nomarski optics (see below, microscopy). Sterility was scored based on the number of total F1 progeny per plate, and embryonic morphology defects/larval arrest were counted per 100 embryos/larvae, respectively.

In the repeat RNAi analysis of the 50 identified trafficking genes all RNAi clones were examined in two rounds of experiments, one focusing on embryonic, the other on early larval development. Intestinal and canal apical/lumenal membrane biogenesis was assessed in parallel in the same animal. Absence/presence of specific phenotypic subclasses (e.g. basolateral or vacuolar ERM-1::GFP displacement) was examined in separate sets of experiments using empirically determined gene-specific RNAi conditions, as described above.

For the extended RNAi analysis of apical membrane polarization, positioning and expansion defects, embryonic and larval intestinal and excretory canal tubulogenesis and ERM-1::GFP placement were evaluated ~q4-6 hours over day during the window of ~48 hours required for full tube extension (mostly longer, given gene-specific growth delay), throughout the process of net polarized membrane expansion (see text and **Fig.S1**). Later occurring phenotypes (>/=72 hours) were not included, given the typical appearance of nonspecific late effects caused by membrane biogenesis defects. This analysis revealed that intestinal membrane polarity defects (basolateral ERM-1::GFP mislocalization) could be obscured during polarized membrane expansion by: increasing cytoplasmic ERM-1::GFP displacement; the development of structural apical/lumenal membrane defects (e.g. lumen widening); vacuolar ERM-1::GFP displacement; other evolving tubulogenesis defects.

**Genetic interaction screens**

*let-767(s2819)* mutants balanced with the free duplication *sDp3* and labeled with the ERM-1::GFP transgene were used for the genetic interaction screens (genotype: *let-767(s2819) ncl-1(e1865) dpy-17(e164) unc-32(e189)III; sDp3(III;f); erm-1p::erm-1::gfp*; *rol-6p::rol-6(su1006)*). See **Fig.3B** for screen design and read-out. *let-767(s2819)* is a maternal-effect larval lethal mutant, thus even homozygous progeny of the balanced strain are not complete nulls due to presence of maternal product (*7*). Although the same *let-767(s2819)* mutant strain was used for enhancer and suppressor screens, each screen was carried out separately. For the enhancer screen, 40 - 60 gravid hermaphrodites were seeded with bleaching solution (4:1 bleach solution/10M NaOH) onto each RNAi plate. The surviving eggs were allowed to hatch and grow for 42 – 46 hours. The same generation of *let-767(+/-)* animals was evaluated and followed for an additional 2-3 days or until they grew into adults. The suppressor screen was carried out by seeding 5 L4-stage larvae on each RNAi plate. Worms were allowed to grow up and lay eggs on the RNAi plates. After 70 – 74 hours, at least 70 next generation *let-767(-/-)* dumpy L1 larvae per plate were hand-picked onto a new RNAi plate. Their development was followed at least until day five to allow for full expression of the ectopic lumen phenotype. In each set of experiments, 50 - 120 live animals per plate were scored for the presence/absence of the intestinal polarity conversion phenotype under a dissecting fluorescence microscope (see below, microscopy), and each experiment was repeated three to five times.

**Fluorescently labeled fusion proteins**

All strains carrying fluorescently-labeled fusion proteins used in this study were previously generated by us and others and have been described (see **Tab.S4**; *(87, 88)* for technical approach). All integral and submembranes apical and basolateral membrane proteins used in this study are expressed from their endogenous promotors and are translational fluorophore fusions. Their subcellular localization has been confirmed by us and others by different labeling procedures, including distinct transgenes, antibodies, germline knock-ins, chemical staining; and by their ability to rescue the corresponding mutant phenotype (see text and *(87, 88)*). The low copy number *erm-1p::erm-1::gfp* transgene, previously extensively characterized and shown to be devoid of any phenotypic effects, is suited for its use as apical domain identity marker since it avoids any disturbance of the *erm-1* germline locus that might interfere with apical domain biogenesis (*7, 11, 30, 31*). ERM-1's subcellular localization has been previously confirmed by various independent transgenic strains (high and low copy number, labeled with different fluorophores), and by an ERM-1::GFP CRISPR knock-in (*30, 31, 93*). The PAR-6::GFP and PKC-3::GFP strains used in this study are CRISPR knock-ins. All vesicle-based fluorescently labeled proteins are translational fusions driven to the intestine by the *vha-6* promotor to allow for the subcellular positional analysis of these ubiquitously expressed molecules (see **Tab.S4** for list of strains used in this study).

**DsRed feeding**
DsRed HT115 RNAi bacteria (**Tab.S4**) were generated with a DsRed expressing plasmid to produce a faint red color. Animals were fed on plates containing RNAi bacteria targeting secretory pathway components and control RNAi bacteria (no vector) for 2 days. At least 70 animals were transferred to plates containing a 1:1 mixture of gene-specific- and of DsRed containing RNAi bacteria, at least 15 hours before evaluation.

**BODIPY-Ceramide assay**
BODIPY-Ceramide (see **Tab.S4**) at a final concentration of 1% was added to bacteria seeded onto the RNAi feeding plates. 15 hours after seeding, 5 L3 larvae were placed onto the RNAi plates and their L3 progeny was evaluated microscopically. *unc-22* RNAi was used as a positive control for these RNAi conditions. Before microscopic evaluation, animals were transferred onto OP50 plates without BODIPY-Cer, to ensure that their intestinal lumens had cleared of the label and their cuticles were cleaned (animals with BODIPY in their intestinal lumen were excluded from the analysis). For each experiment, one individual animal was mounted on a slide and scanned. The time between mounting the animal and taking the image did not exceed 1.5min. See below (microscopy and quantification) for confocal analysis and fluorescence intensity measurement.

**Enhancement of *let-767(RNAi)*-induced ectopic lumen formation by activated RAB-7**
To test for the ability of RAB7Q68L (**Tab.S4**) to enhance polarity conversion, mild *let-767* RNAi conditions were empirically determined (see above, RNAi), with the goal to generate progeny that developed ectopic lumens in less than 5% at the L1 stage (~100% of L1 larvae exposed to *let-767(RNAi)* bacteria under standard RNAi conditions display ectopic lumens). Ectopic lumens in *let-767(RNAi)* intestines were scored by staining for apical intermediate filaments using the anti-IFB-2 antibody MH33 ((*7*); see below, immunohistochemistry and **Tab.S4**). Presence/absence and number of ectopic lumens was assessed by confocal microscopy and the severity of the phenotype was graded from 0, 1, 2, to 3 (no-, few anterior-, multiple anterior ectopic lumens, ectopic lumens along the length of intestine, respectively). 20 L1s were examined per condition in three separate sets of experiments.

**Yolk assay**
Yolk proteins (YP170) are synthesized in the intestine, secreted into the pseudocoelomic space (body cavity), then endocytosed by oocytes. In wild type adult hermaphrodites, YP170::GFP is enriched in oocytes and embryos. Disrupting endocytosis versus secretion cause YP170::GFP to accumulate in the body cavity versus the intestine, respectively (*95*). 5 YP170::GFP L4-stage larvae were placed on each RNAi plate using mild RNAi conditions (empirically determined) to allow progeny to grow up to hermaphrodites with fully developed gonads (see above, RNAi; e.g. *vha-6* RNAi bacteria were diluted 1:4 with mock RNAi bacteria). The effectiveness of these mild RNAi conditions was deemed sufficient to assess intestinal trafficking defects since they produced significant concomitant endocytosis defects in oocytes. YP170::GFP was also evaluated in late larval intestines under standard RNAi conditions, with the same effect as in adult intestines (no intestinal accumulation; YP170::GFP can also accumulate in developing/larval intestines; our unpublished observation in trafficking mutants unrelated to this study).

**Colocalization studies of endo- and plasma-membrane components**

Standard RNAi conditions were used to evaluate the effect of suppressor knockdowns on double transgenic animals carrying green versus red labeled fluorescent fusion proteins of plasma- and endosomal vesicle membranes, respectively. Confocal analysis of L3 and L4 larvae was carried out as described below (microscopy), with the following adjustments (established in pilot experiments to optimize resolution and avoid corruption of signal, loss of signal and false overlap): images were taken as 5 sections along the z axis at 0.15 µm intervals, dorsal and ventral of the mid-intestinal plane (lumen) and images were averaged twice; DAPI exclusion was used to distinguish autofluorescent gut granules, allowing an increase in (Nikon) laser settings to: DAPI: HV=88, 405=5.57, FITC: HV=50, 488=8.76, TRITC: HV=85, 561=51.88; pinhole was set at 40.0µm. Only sequential imaging was performed, DAPI last, as described below (microscopy), and vesicles were examined on all single sections. Overlapping signals observed in both red and green but not blue channels were considered *bona fide* colocalization events (as opposed to autofluorescent signals of *C. elegans* gut granules that appear in all 3 channels).

**Immunohistochemistry**

L1 larvae were collected in M9 medium (*89*) onto slides coated with 0.1-0.2% poly-L-lysine (Sigma, P5899), covered with overhanging coverslips, and then permeabilized by flash freezing in liquid nitrogen and subsequent flicking off the coverslip (*10, 88*). Fixation was performed by sequential incubation in methanol and acetone at -20℃. Immunofluorescent staining was carried out as described (procedures are demonstrated in (*88*)). For MH33 staining, slides were exposed to the first antibody (1:10 dilution; (*7*)) overnight at 4°C, washed and then exposed to the secondary FITC-labeled antibody (diluted 1/100; (*7*)) for 1 hour at room temperature. Permount (Fisher, SP15-100) was used as cover-slide mounting medium.

**Microscopy**

Typically, live worms were directly scored and phenotypically characterized on their plates under an Olympus SZX12 dissecting microscope (Olympus) equipped with a high-power stereo fluorescence attachment (Kramer Scientific). Differential interference contrast (DIC, Nomarski) and confocal microscopic analysis was carried out using either a Leica TCS SL laser-scanning confocal microscope (Leica Microsystem) or a Nikon C2 laser-scanning confocal mounted on an ECLIPSE Ti-E inverted microscope (Nikon), supplemented with DIC optics. Live worms were mounted and immobilized on glass slides using 10mM sodium azide (Fisher Scientific, BP9221-500) or 5% lidocaine (MP Biomedicals,LLG, 193917). Single-plane images were taken as 10 (6-20) sections along the z axis at 0.2 µm or smaller intervals and integrated for projection images. Most images were obtained by a Nikon 60x/1.40 Oil apochromat objective and scanned at 1024x1024 pixels. Laser settings were set at lowest possible gain and laser power where not indicated otherwise and identical settings were used for all comparisons. Autofluorescent gut granules were either visually eliminated/reduced by confocal settings (Leica: by restricting the wavelengths of the fluorescence filters: green filter spectrum to 500-515nm, red filter spectrum to 630-700nm; Nikon by using the following settings: DAPI: HV=108, 405=5.57, FITC: HV=69, 488=6.35, TRITC: HV=105, 561=7.62), or the DAPI channel was used for their identification (*95*). For multi-channel images, individual channel intensity was adjusted to achieve equivalent brightness of all channels, and samples were scanned sequentially to eliminate bleed-through between channels. The images were adjusted by Adobe Photoshop software for contrast and brightness without further changes. No deconvolution software was used.

**Transmission Electron Microscopy (TEM)**
*vha-5*, *vha-6*, *dab-1* and *sly-1* RNAi was performed by standard conditions (see above, RNAi), using either N2, *fgEx13[erm-1p::erm-1::gfp, rol-6p::rol-6(su1006)]* or *rrf-3(pk1426)II*; *fgEx13[erm-1p::erm-1::gfp, rol-6p::rol-6(su1006)]* transgenic animals. More than 20 RNAi plates (60 x 15mm) were set up with 5 L4 larvae and progeny was allowed to develop to the mid-larval stage if not arrested earlier. Before harvesting the progeny, animals were checked under a fluorescent dissecting scope for the presence of phenotype (e.g. presence of vacuolar displacement of the apical marker). Larvae were washed off in standard M9 medium (*89*) and collected into 1.5ml Eppendorf tubes. They were then fixed in 2.5% glutaraldehyde, 1.0% paraformaldehyde in 0.05M sodium cacodylate buffer (pH 7.4) plus 3.0% sucrose. Prior to fixation, the cuticles were 'nicked' with a razor blade in a drop of fixative under a dissecting microscope to allow the fixative to penetrate. After an initial 2-hour fixation at room temperature, the specimens were transferred into fresh fixative and stored overnight at 4°C. Specimens were rinsed several times in 0.1M cacodylate buffer, then post-fixed in 1.0% osmium tetroxide in 0.1M cacodylate buffer for 2 hours on ice. After post fixation, specimens were rinsed several times in 0.1M cacodylate buffer, then embedded in 2.0% agarose in PBS for ease of handling. The agarose blocks were dehydrated through a graded series of ethanol to 100%, dehydrated briefly in 100% propylene oxide and pre-infiltrated overnight on a rocker in a 1:1 mixture of propylene oxide:Eponate resin (Ted Pella, Redding, CA). The following day, the agarose blocks were infiltrated in 100% Eponate resin for several hours, then embedded in flat molds in fresh Eponate resin and allowed to polymerize a minimum of 24 hours at 60°C. Thin sections were cut on a Leica UC7 ultramicrotome and collected on formvar-coated grids, post-stained with uranyl acetate and Reynold's lead citrate, and viewed in a JEOL 1011 TEM at 80 kV equipped with an AMT digital imaging system (Advanced Microscopy Techniques, Danvers, MA).

**Quantification and statistical analysis**

**Hierarchical clustering**
The phenotypic profiles of the 50 identified trafficking genes were analyzed by average linkage agglomerative hierarchical clustering with a centered correlation coefficient as similarity metric of the gene profile for all genes using Cluster 3.0 software (*96*). Phenotype strength was defined as: 0, no phenotype; 1, phenotype identified by extended phenotypic evaluation; 2, phenotype identified by initial tubulogenesis screens. The hierarchical clustering dendrograms were visualized with Java TreeView software (*97*).

**Quantification of BODIPY-Cer vesicles and fluorescence intensity measurement**
Prior to quantification, inspection of worms under a fluorescence dissecting microscope revealed no appreciable difference in quantity, size and distribution of BODIPY-Cer vesicles between RNAi and control worms at different larval stages. L1 – L4 larvae were tested in several sets of pilot experiments for confocal fluorescence intensity measurements and quantification of BODIPY-Cer vesicles (not shown), and L3-stage worms were found to be best suited for standardization (size, RNAi effect, uptake of BODIPY-Cer). Results reported here reflect one set of quantification experiments performed with L3-stage larvae (5 L3s per each condition; 3

separate sets of experiments). For fluorescence intensity measurements, a square area of 250 x 200 pixels was chosen in the anterior intestine, corresponding to four cells (INT II and INT III) and ImageJ (NIH) software was used for image analysis. For quantification of vesicles, two circles of 100 pixels of the same area were counted.

**Fluorescence intensity measurement and quantification of plasma membrane components**
All images were analyzed by ImageJ software. To determine membrane versus cytoplasmic fluorescence intensity ratios between different animals, identical confocal laser settings were used for each set of experiments. 50-pixel long lines were drawn orthogonally across each of the four anterior-most lateral membranes between the first-to-second, and second-to-third intestinal rings (INTII and III). The mid-point of the line was always placed on the lateral membrane. The medians of the maximum fluorescence intensity values were used for both membranes and the cytoplasm, and the membrane/cytoplasm ratios of the four measured cells were averaged and compared to controls. The average of four membrane/cytoplasm ratios was calculated, and more than 7 sets of serial confocal images were analyzed. The average from three independent experiments was used to draw the bar graph. For quantification of the PEPT-1::DsRed, PAR-6::GFP, GFP::PKC-3 and GFP::RAB-10 membrane/membrane-associated/cytoplasm fluorescence intensity ratio, 3 cells of each worm were used for fluorescence intensity quantification, and >10 worms for calculating the final membrane/cytoplasm intensity ratio. For quantification of the presence/absence of hTAC on the plasma membrane, the four lateral membranes between the first three anterior intestinal INT rings (INT I/II and INT II/III) of suppressor knockdowns versus controls were examined for the absence, reduction and presence of hTAC. More than 30 animals were counted per experiment, and each experiment was repeated three times.

**Quantification of colocalization of endo- and plasma-membrane components**
Animals labeled with fluorescent endo- and plasma-membrane components were scanned by confocal microscopy (see above, colocalization studies). Per worm, fluorescent puncta were counted separately inside two intestinal cells, outlined by a fluorescently-labeled basolateral membrane marker. Puncta were subjectively determined as endosomal vesicles, considering factors such as shape, size, boundary, and signal intensity (all fluorescent markers used were previously determined to reside on endo- and/or plasma membranes, see text). Vesicles were classified into three categories – membrane-associated (puncta overlapping with plasma membrane components), tubular (puncta in connection to a cytoplasmic tubular structure that may or may not be connected to the membrane), and cytoplasmic (puncta separated from surrounding structures). 8 animals were counted per each experiment and more than three independent experiments each were performed.

**Statistics**
Statistical analyses were performed by Graphpad Prism 5 software. All values are mean $\pm$ SEM of three or more independent experimental data sets. P values were calculated by student's two-tailed *t-test*. N (sample size) and n (number of independent experiment) are indicated in the text and in the figure legends where necessary. $^{*}p < 0.05$, $^{**}p < 0.01$, $^{***}p < 0.001$.

Fig. S1

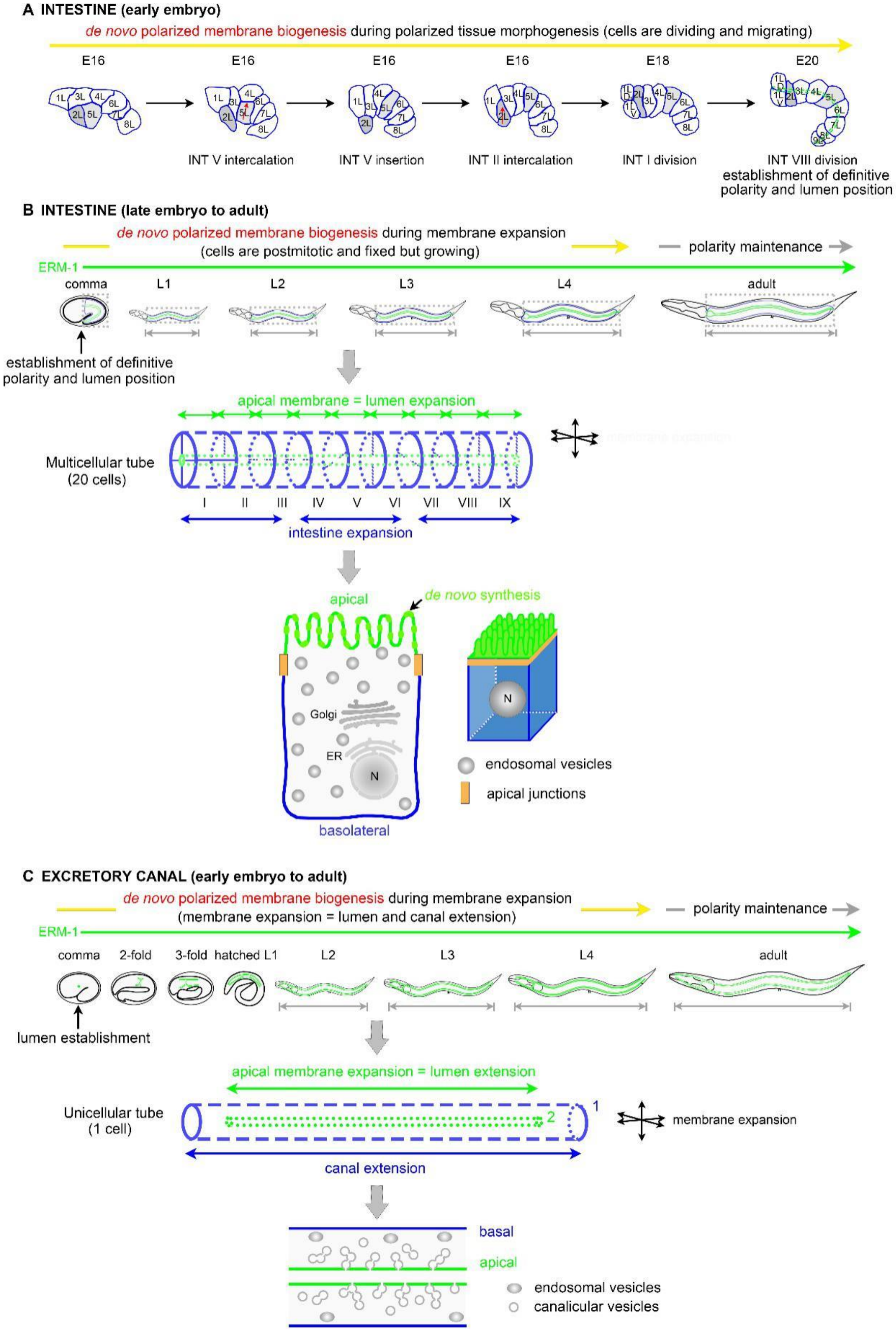

A INTESTINE (early embryo)
de novo polarized membrane biogenesis during polarized tissue morphogenesis (cells are dividing and migrating)
E16
E16
E16
E16
E18
E20
INT V intercalation
INT V insertion
INT II intercalation
INT I division
INT VIII division
establishment of definitive polarity and lumen position
B INTESTINE (late embryo to adult)
de novo polarized membrane biogenesis during membrane expansion
(cells are postmitotic and fixed but growing)
polarity maintenance
ERM-1
comma
L1
L2
L3
L4
adult
establishment of definitive polarity and lumen position
apical membrane = lumen expansion
Multicellular tube (20 cells)
I II III IV V VI VII VIII IX
intestine expansion
apical
de novo synthesis
Golgi
ER
N
basolateral
endosomal vesicles
apical junctions
C EXCRETORY CANAL (early embryo to adult)
de novo polarized membrane biogenesis during membrane expansion
(membrane expansion = lumen and canal extension)
polarity maintenance
ERM-1
comma
2-fold
3-fold
hatched L1
L2
L3
L4
adult
lumen establishment
apical membrane expansion = lumen extension
Unicellular tube (1 cell)
1
2
membrane expansion
canal extension
basal
apical
endosomal vesicles
canalicular vesicles

**Figure S1.** ***De novo*** **polarized membrane biogenesis during** ***C. elegans*** **intestinal and excretory canal tubulogenesis (related to Figure 1A).**
The *C. elegans* intestinal epithelium is single-layered (**A, B**) and its excretory canal consists of one single cell (**C**). Apical membrane and lumen biogenesis occur coincidently in both tubes. These simple organs are thus uniquely suited for the visual distinction of the biogenesis of apical (=lumenal) from lateral and basolateral membranes. In postmitotic cells of growing late-embryonic and larval tubes (**B, C**), *de novo* polarized membrane biogenesis (yellow arrow) can be separated from concomitant effects of polarized tissue morphogenesis (**A**). This allows the separation of changes in polarized trafficking and polarized cytoskeletal dynamics that concomitantly support polarized membrane biogenesis, polarized cell division and polarized cell migration during tissue morphogenesis.

ERM-1, the single *C. elegans* ortholog of the ERM/ezrin-radixin-moesin family of membrane-actin linkers, is asymmetrically positioned at all apical/lumenal membranes of tubular internal organ epithelia that do not secret cuticle (*30*). In tubular epithelia, ERMs denote membranes with apical character, poised to form specific microdomains such as microvilli (*27, 28*). ERM-1 is strictly localized to the apical domain, where it tracks apical/lumenal membrane biogenesis from the time of the establishment of the definitive apicobasal polarity and lumen position in the intestine and the time of lumen initiation in the excretory canal (early embryo, approximately comma stage; **A,** left side of **B-C**). It continues to mark the apical/lumenal domain through its expansion during postmitotic tube growth (late embryo and four larval stages, **B-C**, middle), as well as during its maintenance in the fully grown adult tube (only minimal further expansion; right side of **B-C**). Temporal (early-embryo to adult [yellow]) and spatial vectors for net membrane expansion (apical/lumenal [green], basolateral [blue] and circumferential [black]) are indicated.
**(A)** Early-embryonic intestine: schematics of polarized tissue and membrane biogenesis during establishment of the definitive intestinal membrane polarity and lumen position. The 20 intestinal cells are clonally derived from a single progenitor cell (E). Tube morphogenesis involves transformation of a double-layered into a single-layered tubular epithelium, driven by one intercalation step during which some cells still divide (*42, 43*). Intercalation proceeds via the parallel migration of right and left cells of the lower into the upper tier (one set of an INT 1 – 8 doublet is shown), after which cell division and migration are mostly complete.
**(B)** Late-embryonic, larval and adult intestine: schematics of coincident apical membrane and lumen biogenesis between pairs of expanding postmitotic cells. The mature intestinal tube is bilaterally symmetrical with 9 INT rings consisting of 2 cells each (INT I-IX, 4 cells in first ring; enlarged middle diagram; compare **Fig.1A**). The tube's apical membrane (green) faces and concomitantly builds the lumen, while the basolateral membrane (blue) contacts neighboring cells or the body cavity (top diagrams); apical junctions separate the polarized domains (orange; bottom diagrams of single cell). *De novo* polarized membrane biogenesis continues in expanding postmitotic cells from the comma stage through the 2- and 3-fold embryonic stages (not shown) and four larval stages (L1 - L4) to adulthood (with minimal further expansion; top diagram). **The anterior-posterior length of the apical/lumenal intestinal membrane increases from approximately 20 to 680um from the early-embryonic bean to the L4-larval stage. Note that the apical domain of a single cell additionally requires a multiple of its ~1/6$^{th}$ overall cellular membrane surface for microvilli formation (3D view, bottom diagram).** The subcellular localization of organelles changes during membrane polarization and membrane expansion. This includes the position of the nucleus (N) that moves towards the apical domain at

the beginning of intestinal intercalation in the early embryo (*43*). Golgi mini-stacks are distributed throughout the cytoplasm in the adult epithelium (*48*). Location and size of intracellular organelles are not drawn to scale.

**(C)** Excretory canal: schematics of coincident apical membrane and lumen biogenesis inside the single expanding postmitotic cell. During early embryogenesis, the excretory cell migrates to its final location at the left lateral side of the posterior pharyngeal bulb (not shown), where it sends out two extensions to the left and right lateral surfaces (top diagrams) (*92*). At the lateral hypodermis, each canal extension bifurcates and grows anteriorward and posteriorward (top diagrams; compare **Fig.1A**). Canals grow directional at a speed exceeding that of the animal until the end of the L1 stage, when they reach the tip of the worm's nose and tail. At hatching, the posterior branches are extended halfway through the length of the animal. The apical/lumenal membrane (green) expands inside the cell body and its expansion follows that of the basolateral canal membranes (blue). The excretory cell lumen is connected to the lumen of the adjacent duct cell, where apical and basolateral membranes meet, separated by a single junction (not shown); canals are otherwise junction-less. ***De novo*** **polarized membrane biogenesis in the excretory canal continues from lumen initiation at approximately the time of comma stage in the early embryo through the 2- and 3-fold embryonic and four larval stages (L1 – L4) until adulthood, with the anterior-posterior apical/lumenal membrane length increasing up to ~2000um.** Note that growth of the basal membrane (1, blue) precedes that of the apical/lumenal membrane (2, green; middle diagram). The enlarged view of a canal arm section (bottom diagram) shows canalicular and endosomal vesicles, the former partially interconnected and connected to the apical/lumenal membrane.

### Fig. S2

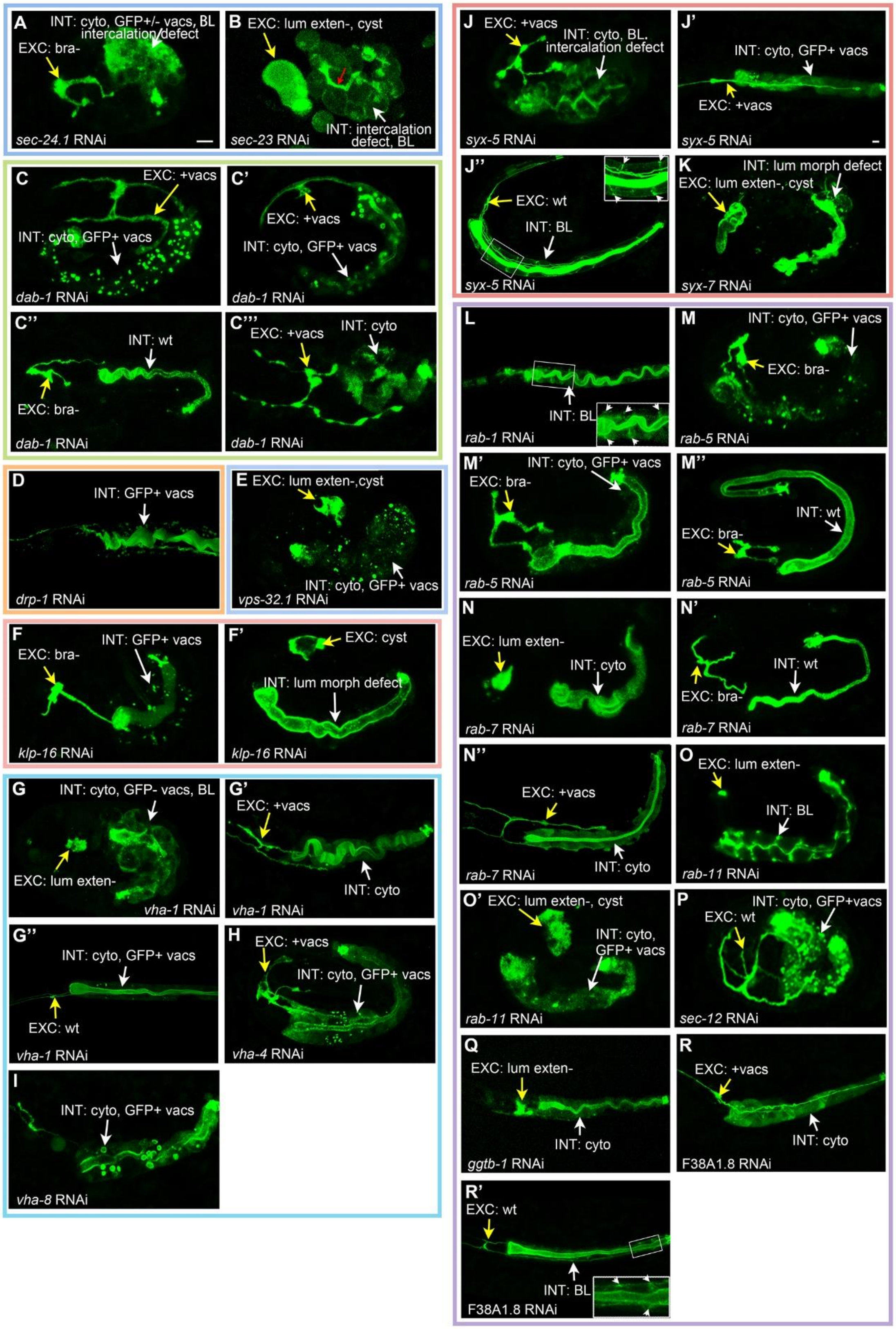

A
INT: cyto, GFP+/- vacs, BL
intercalation defect
EXC: bra-
sec-24.1 RNAi
B
EXC: lum exten-, cyst
INT: intercalation defect, BL
sec-23 RNAi
C
EXC: +vacs
INT: cyto, GFP+ vacs
dab-1 RNAi
C'
EXC: +vacs
INT: cyto, GFP+ vacs
dab-1 RNAi
C''
INT: wt
EXC: bra-
dab-1 RNAi
C'''
EXC: +vacs
INT: cyto
dab-1 RNAi
D
INT: GFP+ vacs
drp-1 RNAi
E
EXC: lum exten-,cyst
INT: cyto, GFP+ vacs
vps-32.1 RNAi
F
EXC: bra-
INT: GFP+ vacs
klp-16 RNAi
F'
EXC: cyst
INT: lum morph defect
klp-16 RNAi
G
INT: cyto, GFP- vacs, BL
EXC: lum exten-
vha-1 RNAi
G'
EXC: +vacs
INT: cyto
vha-1 RNAi
G''
INT: cyto, GFP+ vacs
EXC: wt
vha-1 RNAi
H
EXC: +vacs
INT: cyto, GFP+ vacs
vha-4 RNAi
I
INT: cyto, GFP+ vacs
vha-8 RNAi
J
EXC: +vacs
INT: cyto, BL,
intercalation defect
syx-5 RNAi
J'
INT: cyto, GFP+ vacs
EXC: +vacs
syx-5 RNAi
J''
EXC: wt
INT: BL
syx-5 RNAi
K
INT: lum morph defect
EXC: lum exten-, cyst
syx-7 RNAi
L
INT: BL
rab-1 RNAi
M
INT: cyto, GFP+ vacs
EXC: bra-
rab-5 RNAi
M'
INT: cyto, GFP+ vacs
EXC: bra-
rab-5 RNAi
M''
INT: wt
EXC: bra-
rab-5 RNAi
N
EXC: lum exten-
INT: cyto
rab-7 RNAi
N'
INT: wt
EXC: bra-
rab-7 RNAi
N''
EXC: +vacs
INT: cyto
rab-7 RNAi
O
EXC: lum exten-
INT: BL
rab-11 RNAi
O'
EXC: lum exten-, cyst
INT: cyto,
GFP+ vacs
rab-11 RNAi
P
INT: cyto, GFP+vacs
EXC: wt
sec-12 RNAi
Q
EXC: lum exten-
INT: cyto
ggtb-1 RNAi
R
EXC: +vacs
INT: cyto
F38A1.8 RNAi
R'
EXC: wt
INT: BL
F38A1.8 RNAi

**Figure S2. A shared set of trafficking molecules directs inTER- and inTRAcellular apical membrane polarization, positioning and expansion (related to Figure 2B).**
Selection of additional apical domain biogenesis defects and description of phenotypes. A spectrum of phenotypes of different severity is shown (see Methods, **Fig.2A,B** and **Fig.2B** legend for analysis and distinction of defects along the spectrum of severe to mild intestinal (INT) and excretory canal (EXC) apical membrane biogenesis/lumenogenesis defects; **Tab.S1** for phenotype classes 1 - 10 of all 50 genes; **Figs.S1** for schematics of INT and EXC tubulogenesis). Confocal projections of embryonic and larval phenotypes are shown (embryos: A-C, E-G, H-I, J"-K, M-N', O-P, among them early: A-B, G, J, all others late; L1-larvae: D, G'-G", L, N", Q-R'), with ERM-1::GFP marking the apical domain/lumen in both INT and EXC. See **Fig.2** for acronyms and structural classes of molecules (classes outlined in color) and **Fig.2B** for wild-type body plan of embryos/larvae and corresponding wild-type intestinal and canal ERM-1::GFP localization. Arrows point to selected defects only, see description below for details. Scale bars: 5µm. White arrows point to INT, yellow arrows to EXC. Color coding of functional classes (boxes) as in **Fig.1C**.

**Narrative** (main **Fig.2B** images are included in description; not all aspects of phenotypes are described). ***sec-23/24***: early-embryonic apical membrane polarization, positioning and expansion defects. INT: no ERM-1 at apical domain (no apical membrane polarization) in all 4 images (including those of Fig.2B); partial presence in *sec-23* in this figure; red arrow in Fig.S2B), with ERM-1 displacement to the cytoplasm and all sides of the membrane (no apicobasal membrane polarity); EXC: no lumen extension with cystic deformation to short lumen/lumen arms (no apical membrane expansion). ***syx-5***: spectrum of early-embryonic to larval apical membrane polarization and positioning defects. Fig.2B, Fig.S2J – J": INT: both cytoplasmic (vacuolar; Fig.2B) and basolateral ERM-1 mislocalization in embryo, predominant basolateral mislocalization in larvae. EXC: mildly reduced lumen extension and cytoplasmic ERM-1 displacement. ***syx-7*:** defect during late step of apical domain assembly and apical membrane expansion defect. INT: near complete apical membrane polarization. EXC: minimal lumen extension and cystic deformation (EXC should be extended beyond bifurcation at this stage; compare to similar EXC phenotypes in *sec-23* and *rab-11(RNAi)* animals). Polarity defects not shown. ***dab-1***: spectrum of late embryonic to early larval apical membrane polarization but not positioning defects. Fig.2B, Fig.S2C – C"': INT/EXC: decreasing severity of vacuolar ERM-1 displacement from full displacement (no apical membrane polarization) to no or homogeneous cytoplasmic displacement. Note absence of basolateral ERM-1 mislocalization (no apicobasal membrane polarity defect). ***rab-1***: spectrum of late-embryonic (Fig.2B) to larval apical domain polarization, positioning and expansion defects. INT: ERM-1 displacement to cytoplasmic vacuoles and basolateral membrane in embryo (Fig.2B), mild basolateral displacement in larva. EXC: mild extension defect (Fig.2B). ***rab-5***: spectrum of late-embryonic apical membrane polarization and expansion defects. Fig.2B, Fig.S2 M – M": INT: full vacuolar ERM-1 displacement; mild cytoplasmic ERM-1 displacement; wild-type apical ERM-1. No apicobasal membrane polarity defect. EXC: no lumen extension; lumen branch loss. ***rab-7***: spectrum of late-embryonic apical membrane polarization and expansion defects. Fig.2B, Fig.S2 L - R: INT: decreasing severity from: almost full vacuolar ERM-1 displacement (Fig.2B); cytoplasmic ERM-1 displacement (Fig.S2N,N"); apical ERM-1 (Fig.S2N'). No membrane polarity defect. EXC: from Fig.2B to Fig.S2N": no lumen extension; short lumen; full lumen extension. ***rab-11***: late-embryonic apical membrane polarization, positioning and expansion defect. INT: strong cytoplasmic ERM-1 displacement; basolateral ERM-1 is only transiently detected during late-embryonic apical membrane biogenesis. EXC: no lumen extension. ***sec-12***: intestine-specific apical membrane polarization and positioning defects in early and late embryo. INT: no apical membrane polarization in early (Fig.S2P), and cytoplasmic and basolateral displacement in late, embryo (Fig.2B); wild-type EXC in these images (posterior arms twisted in section shown in Fig.S2P). ***ggtb-1***: discordant severity of INT/EXC apical membrane biogenesis defects in early larvae. Strong vacuolar displacement (INT) and wild-type EXC (Fig.2B) versus mild cytoplasmic displacement (INT) and short lumen (EXC; polarity defect not shown). **F38A1.8**: spectrum of early-embryonic to larval apical membrane polarization and positioning defects. Range of INT phenotypes from cytoplasmic plus vacuolar- to strong cytoplasmic-, to mild basolateral ERM-1 displacement in larvae. ***drp-1*****:** mild apical membrane polarization defect. Small-vesicular ERM-1 displacement phenotype (compare to large vacuolar ERM-1 displacement in *dab-1* and *vha-8(RNAi)* INTs). ***vps-32*****:** embryonic apical membrane polarization but not positioning defects. INT: full vacuolar ERM-1 displacement from the apical domain but no appearance at the basolateral membrane (compare to *dab-1*). ***klp-16***: spectrum of embryonic apical membrane polarization and expansion but no positioning (apicobasal polarity) defects. Spectrum ranges from full vacuolar ERM-1 displacement (no apical membrane polarization, Fig.2B) to mild vacuolar displacement and partial apical membrane polarization (Fig.S2F) to full apical membrane polarization (Fig.S2F') in the INT; and from no lumen extension (Fig.2B) to varying degrees of extension defects (EXC). ***vha-1/4***: spectrum of early-embryonic to larval apical membrane polarization and positioning defects. INT: combination of cytoplasmic vacuoles with ERM-1-positive versus ERM-1-negative membranes. EXC: cytoplasmic ERM-1 displacement reveals previously noted enlarged and persistent varicosities in short canals (*31*). ***vha-8:*** apical membrane polarization but no positioning defects. INT: note large vacuolar ERM-1 displacement, distinct from *vha-1/4* (see text for discussion of discrepancy to *(38)*).

### Fig. S3

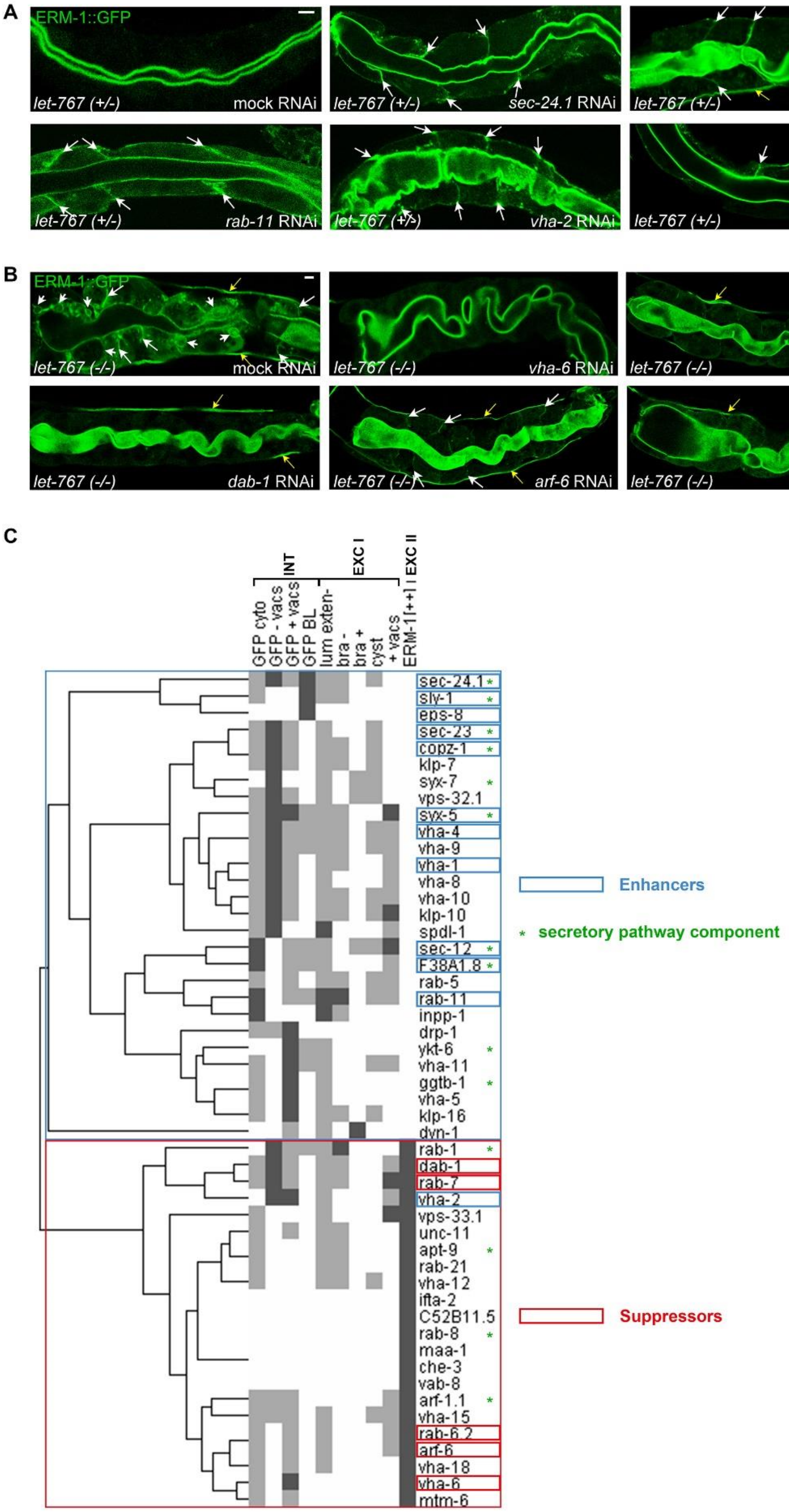
A
ERM-1::GFP
let-767 (+/-) mock RNAi
let-767 (+/-) sec-24.1 RNAi
let-767 (+/-) copz-1 RNAi
let-767 (+/-) rab-11 RNAi
let-767 (+/-) vha-2 RNAi
let-767 (+/-) sly-1 RNAi
B
ERM-1::GFP
let-767 (-/-) mock RNAi
let-767 (-/-) vha-6 RNAi
let-767 (-/-) rab-7 RNAi
let-767 (-/-) dab-1 RNAi
let-767 (-/-) arf-6 RNAi
let-767 (-/-) rab-6.2 RNAi
C
INT
EXC I
EXC II
GFP cyto
GFP - vacs
GFP + vacs
GFP BL
lum exten-
bra -
bra +
cyst
+ vacs
ERM-1[++]
sec-24.1
sly-1
eps-8
sec-23
copz-1
klp-7
syx-7
vps-32.1
syx-5
vha-4
vha-9
vha-1
vha-8
vha-10
klp-10
spdl-1
sec-12
F38A1.8
rab-5
rab-11
inpp-1
drp-1
ykt-6
vha-11
ggtb-1
vha-5
klp-16
dyn-1
rab-1
dab-1
rab-7
vha-2
vps-33.1
unc-11
apt-9
rab-21
vha-12
ifta-2
C52B11.5
rab-8
maa-1
che-3
vab-8
arf-1.1
vha-15
rab-6.2
arf-6
vha-18
vha-6
mtm-6
Enhancers
* secretory pathway component
Suppressors

**Figure S3 Genetic interaction screens identify 17 modifiers of the GSL-dependent apicobasal polarity conversion (related to Figure 3).**

**(A)** Enhancement (compare **Fig.3B**). Additional examples of *de novo* appearance of basolateral ERM-1::GFP (arrows) in growing double *let-767(s2819)(+/-)/enhancer(RNAi)* larval intestines on day 3 post RNAi (Methods). Note intact polarity with strict apical confinement of ERM-1::GFP in *let-767(s2819)(+/-)* haplosufficient mutant background (*sDp3* duplication provides one *let-767* copy; upper left image). Lateral ERM-1::GFP mislocalization is indicated by arrows and basal ERM-1::GFP mislocalization outlines the intestine. Note punctate nature of basolateral ERM-1::GFP recruitment with additional cytoplasmic ERM-1::GFP puncta in membrane vicinity.

**(B)** Suppression (compare **Fig.3B***). vha-6*, *rab-7*, *dab-1*, *arf-6* and *rab-6.2* RNAi suppress basolateral ERM-1::GFP mislocalization and ectopic lumen formation in *let-767(s2819)(-/-)* intestines. On day 5 post RNAi (standard RNAi, Methods), greater than 90% of *let-767(-/-)* mutants (no *let-767* copy, *sDp3* duplication lost; upper left image) display basolateral ERM-1::GFP displacement (arrows) and ectopic lumens (arrowheads). In double mutant/RNAi animals (all other images) ERM-1::GFP is confined to the apical membrane (*vha-6*, *dab-1*) and/or only mildly displaced to basolateral membranes (*rab-7, arf-6, rab-6.2*; representative day 5 images are shown). The brightness of images of double mutant/RNAi intestines is increased to show the absence/reduction of basolateral ERM-1::GFP (resulting in the false appearance of an ERM-1::GFP increase at the apical domain). Animals are short and fat and lumens are wide due to Dpy background.

Confocal sections of larval intestines are shown throughout. Yellow arrows: EXC. Scale bars: 5μm.

**(C)** Hierarchical clustering of phenotype profiles (Methods). Most enhancers (blue rectangles) were recovered from intestinal screens and all suppressors (red rectangles) from excretory canal screens. Secretory pathway components (green asterisk) track with phenotype signatures of enhancers.

**Fig. S5**

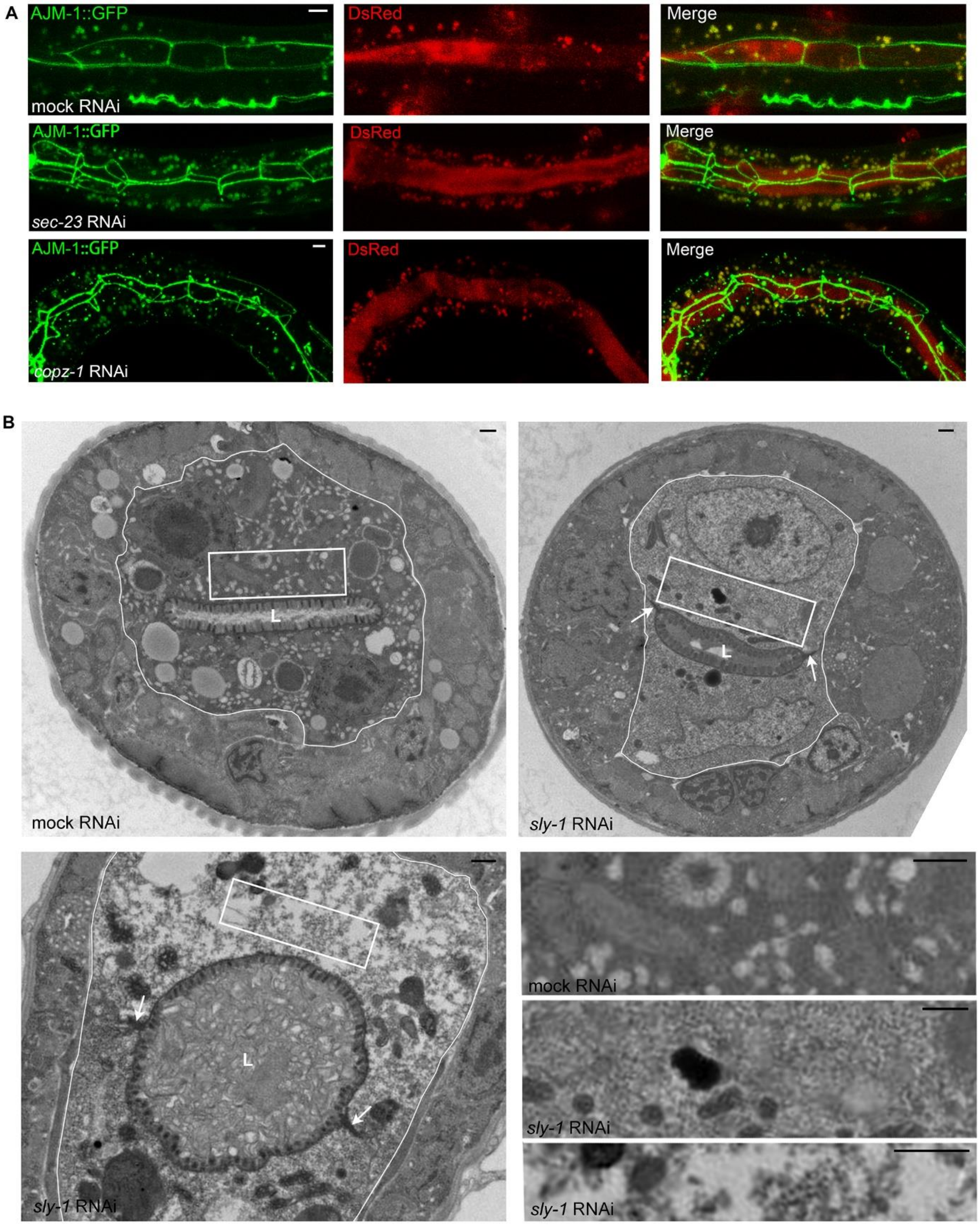
A
AJM-1::GFP
DsRed
Merge
mock RNAi
AJM-1::GFP
DsRed
Merge
sec-23 RNAi
AJM-1::GFP
DsRed
Merge
copz-1 RNAi
B
L
mock RNAi
L
sly-1 RNAi
L
sly-1 RNAi
mock RNAi
sly-1 RNAi
sly-1 RNAi

**Figure S5. Effects of early secretory pathway disruption on the assembly of apical junctions and endomembranes (related to Figure 5).**

**(A)** Integrity of apicolateral junctions but presence of ectopic junction material. *sec-23*- and *copz-1(mildRNAi)* larvae, fed with DsRed-labeled bacteria, maintain wild-type contiguity of AJM-1-labeled junctions and do not allow leakage of DsRed between basolateral membranes (Methods). Note increase of punctate junction material (bright small GFP spots, distinct from larger autofluorescent granules; see below) in *copz-1(RNAi)* intestinal cytoplasm (a few such puncta are occasionally also seen in wild-type intestines, e.g. in **Fig.5B**). Broadening of junctions into the lateral membrane (indicated by arrows in **Fig.5B**) is here obscured by slightly distorted, albeit intact, junction pattern. For junction contiguity in these areas, see **Movie S1** (rotation of 3D projection). Enlarged autofluorescent gut granules are present throughout and can be distinguished by their appearance in both green and red channels (yellow in overlay).

**(B)** TEM images of cross-sections of wild-type and *sly-1(RNAi)* larvae. Intestine (2 cells visualized) is delineated by white line. Moderately-severe (upper right) and strong (lower left) *sly-1(RNAi)* phenotypes are shown. Note mild apical/lumenal membrane defects with reduced number and length of microvilli and widened lumen (L), marked paucity of all endo-/vesicle membranes in *sly-1(RNAi)* intestines. Apical junctions appear structurally intact (arrows). Representative sections are shown (75 random sections of *sly-1(RNAi)* L1 larval intestines with typical endomembrane defects were evaluated; compare **Fig.5C**). Increasing dispersion of endomembranes in *sly-1(RNAi)* cytoplasm shown at bottom right at higher magnification (corresponding to boxed areas in cross sections). Note that these effects of *sly-1* RNAi are largely confined to intestines. Normal appearing cross-sections of excretory canal arm lumens and surrounding canalicular vesicles in both *sly-1(RNAi)* larvae (arrowheads).

Confocal projections of portions of larval intestines are shown in **(A)**. Scale bars, 5µm in **(A)**; 500nm in **(B)**.

### Fig. S6

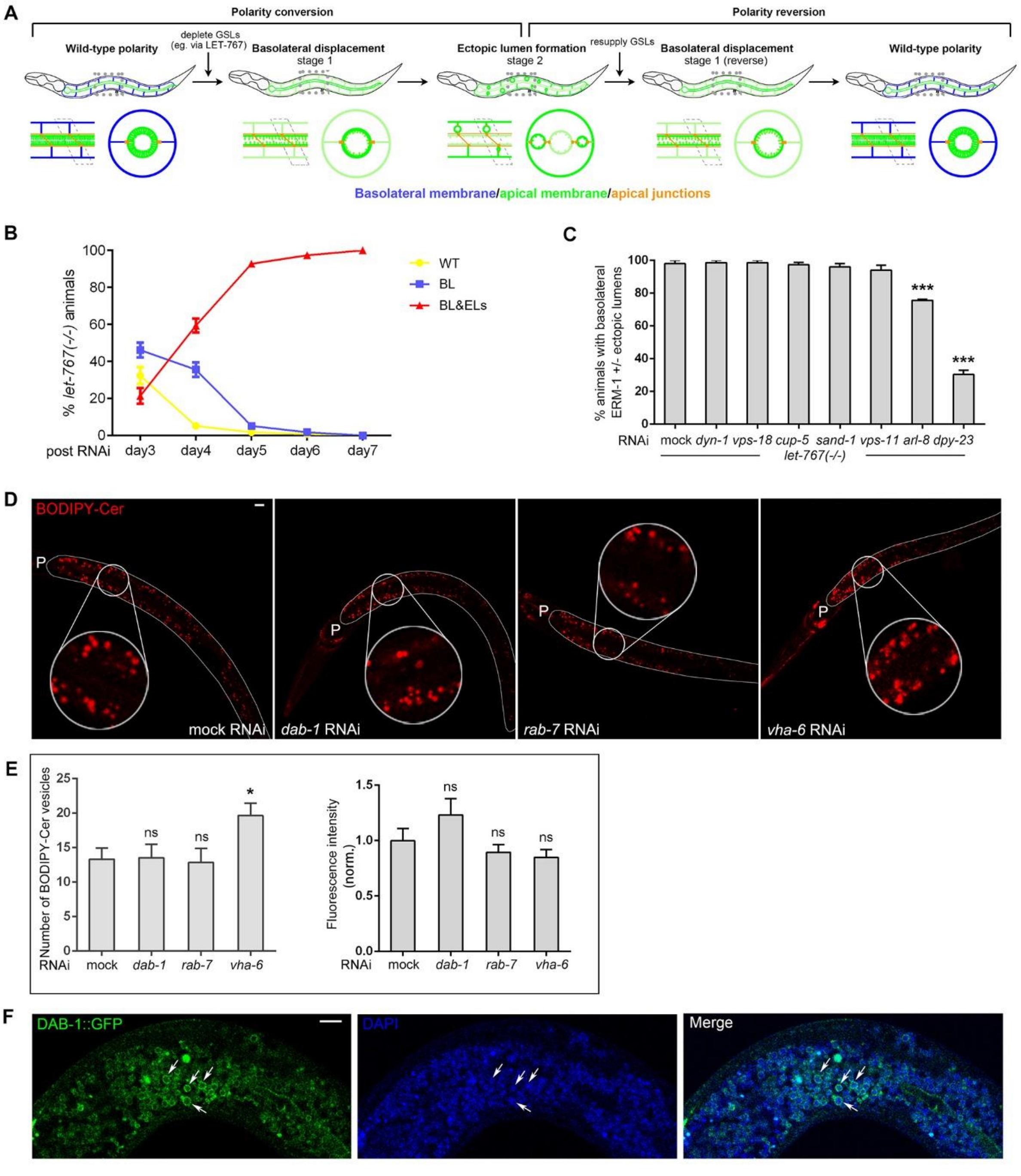
A
Polarity conversion
Polarity reversion
Wild-type polarity
deplete GSLs (eg. via LET-767)
Basolateral displacement stage 1
Ectopic lumen formation stage 2
resupply GSLs
Basolateral displacement stage 1 (reverse)
Wild-type polarity
Basolateral membrane/apical membrane/apical junctions
B
% let-767(-/-) animals
WT
BL
BL&ELs
post RNAi day3 day4 day5 day6 day7
C
% animals with basolateral ERM-1 +/- ectopic lumens
***
***
RNAi mock dyn-1 vps-18 cup-5 sand-1 vps-11 arl-8 dpy-23
let-767(-/-)
D
BODIPY-Cer
P
mock RNAi
dab-1 RNAi
rab-7 RNAi
vha-6 RNAi
E
Number of BODIPY-Cer vesicles
ns
ns
*
RNAi mock dab-1 rab-7 vha-6
Fluorescence intensity (norm.)
ns
ns
ns
RNAi mock dab-1 rab-7 vha-6
F
DAB-1::GFP
DAPI
Merge

**Figure S6. Analysis of suppression (related to Figure 6).**

**(A)** Schematic of apicobasal polarity conversion and reversion in postmitotic cells in the fixed tissue context of the growing *C. elegans* intestine by depletion and repletion of GSL biosynthesis (text; Methods). Growth in GSL-depleted animals is delayed, extending the window for the observation of polarized membrane expansion. From left to right (1-5):

(1) wild-type apicobasal membrane polarity;

(2) stage 1 polarity conversion: basolateral mislocalization of all tested apical membrane components (day 2-3 post standard RNAi induction/Methods; slowly growing L1-larvae);

(3) stage 2: ectopic (at the site of the initial basolateral membrane) apical membrane biogenesis generates basolateral lumens with apical-membrane specific microvilli, sub membranous terminal web and junctions (days 3-5; larvae arrest at L1 stage);

(4) stage 1 (reverse) polarity reversion subsequent to restoring GSL biosynthesis: basolateral mislocalization of the apical domain without ectopic lumen formation;

(5) restoration of wild-type apicobasal polarity and growth (larvae develop into fertile adults).

GSL depletion/repletion can be modulated by various GSL biosynthetic enzymes (*7*). GSL biosynthesis is reduced by placing animals on RNAi bacteria and resumed by removing them from RNAi bacteria to wild-type (OP50) feeding plates. Speed and strength of conversion and reversion can be modulated by the timing of placing and removing animals from RNAi plates and by different RNAi conditions (Methods; standard *let-767* RNAi conditions are used in the suppressor analysis). Top: whole animal; bottom: longitudinal (left) and transverse section (right) through one INT ring. Animals are not drawn to scale (they arrest during polarity conversion and resume growth during reversion).

**(B)** Time-course of polarity conversion in *let-767(s2819)(-/-)* larval intestines (compare to **Fig.6C**). WT, wild-type; BL, basolateral displacement only; BL&ELs, basolateral displacement and ectopic lumens.

**(C)** Disruption of endocytic and degradative trafficking pathways in *let-767(-/-)* mutants. The GTPase DYN-1/dynamin and DPY-23, the mu2 subunit of the clathrin AP2 adaptor (components of endocytic routes) and the mucolipin ortholog CUP-5, the Arf-like GTPase ARL-8, SAND-1/MON1 and the HOPS complex components VPS-11 and VPS-18 (components of vesicle degradative trafficking routes) (*48*) were depleted in *let-767(-/-)* animals using the same conditions as those used in the suppressor screen (compare **Fig.3B**; milder RNAi conditions were used for *dyn-1* to avoid sterility; Methods). Depletion of VPS-33.1, another HOPS complex components (initially identified in tier-1 screens), also failed to suppress, as demonstrated in tier-2 genetic interaction screens (**Fig.2**, **Tab.S1**). n=3; N>30.

**(D)** BODIPY-Cer+ vesicles in *dab-1-*, *rab-7* and *vha-6(RNAi)* larval intestines. Note similar contribution and size of BODIPY-Cer+ vesicles in all intestines. P = pharynx. Intestines are outlined by white lines. BODIPY-Cer in head area labels amphids.

**(E)** Quantification of BODIPY-Cer+ vesicles and fluorescence intensity in *dab-1-*, *rab-7* and *vha-6(RNAi)* larval intestines. Ceramide (Cer) is the chemical backbone of GSLs. Animals were fed BODIPY-Cer, previously shown to reproduce localization and behavior of endogenous intestinal vesicle-associated GSLs (*11*). Left graph shows the number of intestinal BODIPY-Cer+ vesicles counted in the defined area circled in images above, right graph the corresponding fluorescence intensity (Methods). Note increase in BODIPY-Cer+ vesicles but not fluorescence intensity in *vha-6(RNAi)* intestines (compare to **(D)**). n=3; N=5.

**(F)** Vesicular localization of DAB-1. DAB-1::GFP resides on at least two morphologically distinct vesicle populations, one of them visible in the DAPI channel. Note GFP ring around blue vesicles (arrows; compare **Fig.6F**).
Confocal images of larval intestines are shown in **D**, pair of 2 opposing larval intestinal cells in **F.** All data shown as mean ± SEM. *$p < 0.05$, **$p < 0.01$, ***$p < 0.001$. Scale bars: 5μm.

### Fig. S7

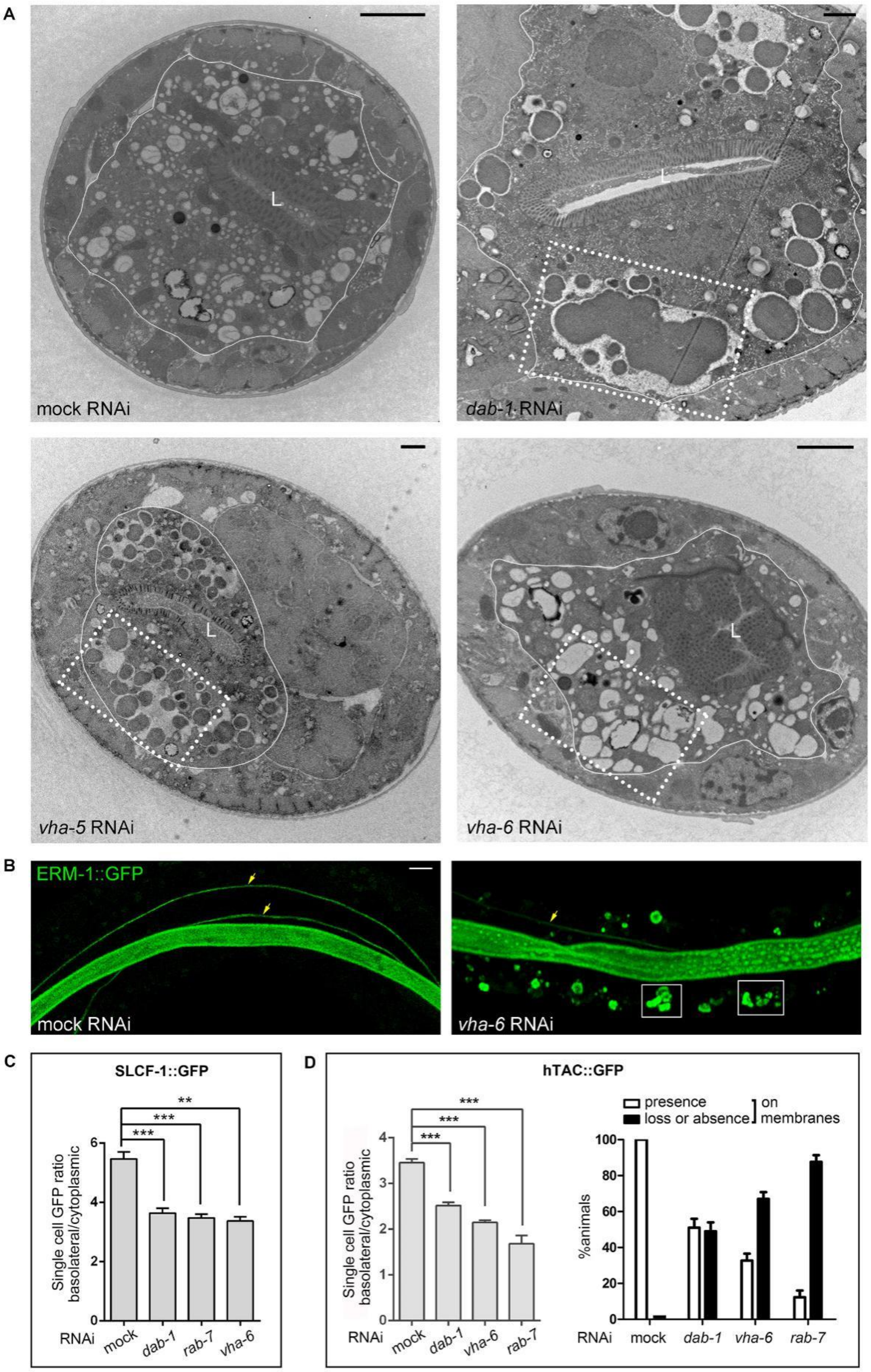
A
L
mock RNAi
dab-1 RNAi
vha-5 RNAi
vha-6 RNAi
B
ERM-1::GFP
mock RNAi
vha-6 RNAi
C
SLCF-1::GFP
Single cell GFP ratio basolateral/cytoplasmic
RNAi mock dab-1 rab-7 vha-6
D
hTAC::GFP
presence
loss or absence
on membranes
%animals
RNAi mock dab-1 vha-6 rab-7

**Figure S7. Cytoplasmic apical membrane inclusions in *dab-1* and *vha-6(RNAi)* intestines are not ectopic lumens. Quantification of the suppressor effect on wild-type basolateral membrane biogenesis and recycling (related to Figure 7).**

**(A)** Representative TEM cross sections of whole *dab-1-, vha-5- and vha-6(RNAi) rrf-3* larval intestines. VHA-5, another V-ATPase subunit and a paralog of VHA-6, copies VHA-6's intestinal loss-of-function phenotype and was independently identified in tier-1 screens (**Tab.S1**). Images show similar vacuolar aggregates in all 3 knockdowns (boxed), with a mixture of amorphous material and intact vesicles, some of which may contain lipids (N>50 random sections each were evaluated; representative images are shown). Subcellular localization, size and clustering of aggregates suggest they correspond to the ERM-1::GFP+ vacuolar clusters (compare to **(B)**). No inward-pointing microvilli were detected in any of these vacuoles in serial TEM sections, suggesting they are not ectopic lumens. The two intestinal cells per image are delineated by a white line. L: lumen. Scale bars: 2µm.

The depletion of V-ATPases has previously been suggested to generate ectopic lumens in *C. elegans* intestines (*38*). We consider it unlikely that ectopic lumens were missed on multiple TEM sections of *vha-5-* and *vha-6(RNAi) rrf-3* intestines. This and other discrepancies to Bidaud-Meynard et al. (*38*) could be explained by our finding that V-ATPase subunits assemble into different pumps in the *C. elegans* intestine that function in different aspects of polarized membrane biogenesis and polarity (LA Khan et al., in preparation). There is precedence for the organelle-specific assembly of different V-ATPase subunits in yeast (*35*). Note that VHA-1, -2 and -4 were identified as enhancers of GSLs' polarity function and are required for apical domain positioning (membrane polarity). In contrast, VHA-6 was identified as GSL suppressor and is required for apical domain polarization but not positioning (**Fig.3D**).

**(B)** Confocal projections of ERM-1::GFP+ vacuolar clusters (2 boxed) in *rrf-3 vha-6(RNAi)* larval intestinal cells (set of 3 opposing cells). Yellow arrows indicate excretory canal arm lumens. Scale bar: 5µm.

**(C)** Basolateral-membrane/cytoplasm SLCF-1::GFP fluorescence intensity ratio in expanding intestines (compare to **Fig.7B**; Methods). Data shown as mean ± SEM, n=3, *p < 0.05, **p<0.01, ***p< 0.001.

**(D)** hTAC::GFP membrane/cytoplasm fluorescence intensity ratio (L) and hTAC::GFP presence/absence on expanding larval intestinal membranes (R; compare to **Fig.7C**; Methods). n=3. N≥7 (left), N≥30 (right).

#### Fig. S8

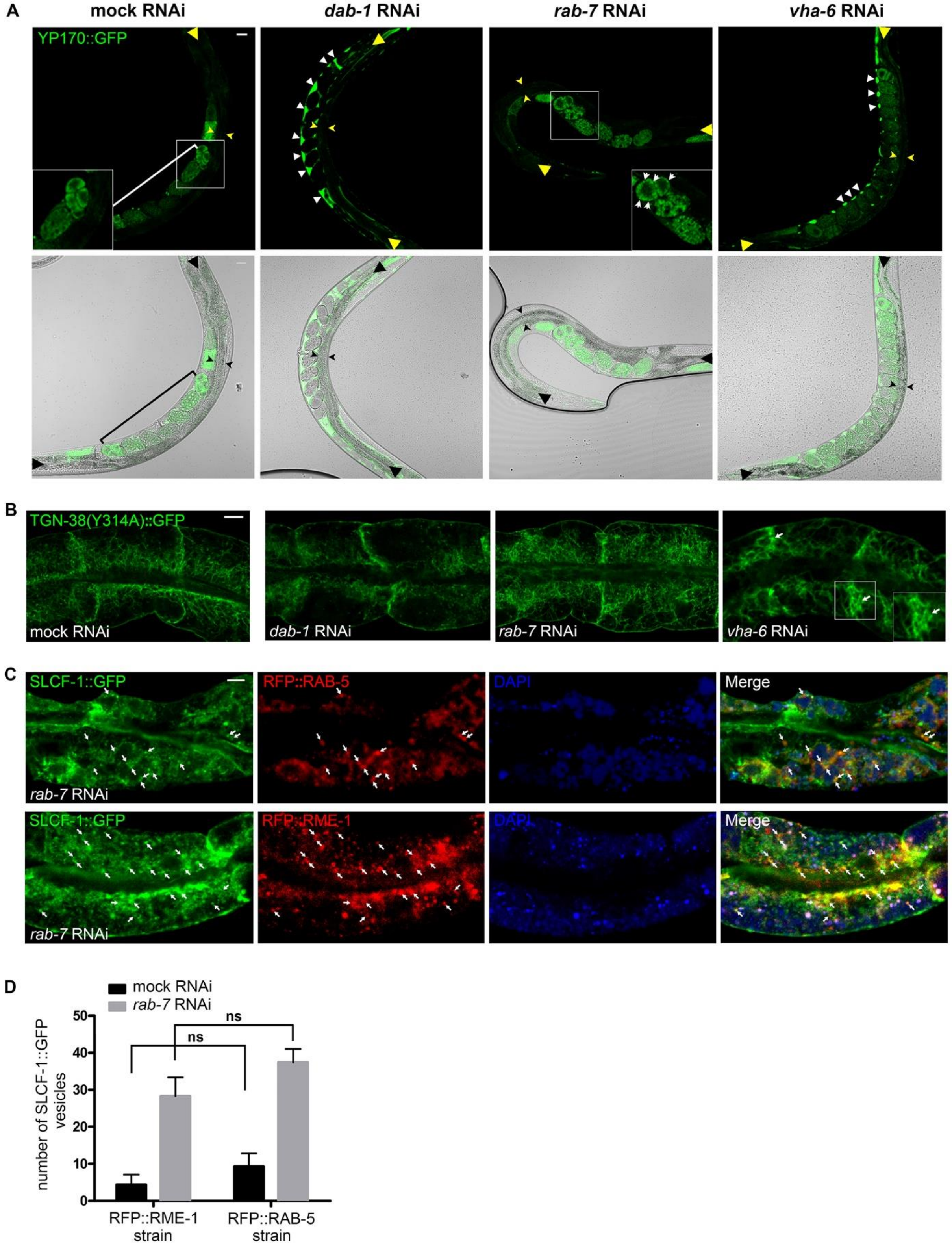
A
mock RNAi
dab-1 RNAi
rab-7 RNAi
vha-6 RNAi
YP170::GFP
B
TGN-38(Y314A)::GFP
mock RNAi
dab-1 RNAi
rab-7 RNAi
vha-6 RNAi
C
SLCF-1::GFP
RFP::RAB-5
DAPI
Merge
rab-7 RNAi
SLCF-1::GFP
RFP::RME-1
DAPI
Merge
rab-7 RNAi
D
mock RNAi
rab-7 RNAi
ns
ns
number of SLCF-1::GFP vesicles
50
40
30
20
10
0
RFP::RME-1 strain
RFP::RAB-5 strain

**Figure S8. DAB-1, RAB-7 and VHA-6 shuttle polarized membrane components on recycling, not secretory, routes during net membrane expansion (related to Figure 8).**
**(A)** *dab-1, rab-7 and vha-6* RNAi fail to disrupt Yolk secretion in intestines of larvae and young adults. YP170/Yolk::GFP is secreted from *C. elegans* intestines into the body cavity (pseudocoelum), then endocytosed by oocytes (Methods). Disrupting endocytosis and secretion cause YP170::GFP accumulation in the body cavity and intestine, respectively (*89*). Note absence of intestinal YP170::GFP accumulation in all suppressor knockdowns, but pseudocoelomic YP170::GFP accumulation in *dab-1(RNAi)* animals (white arrowheads) with coincident lack of Yolk uptake into oocytes/embryos (previously described; (*46*)); YP170::GFP distribution defects in *rab-7(RNAi)* oocytes/embryos (see insets, white arrowheads; also previously described; (*93*); and mild YP170::GFP accumulation in the body cavity of *vha-6(RNAi)* animals (white arrowheads) with reduced uptake in oocytes/embryos. Mild RNAi conditions (Methods) were used to allow animals to grow to maturity for which this assay was established. YP170::GFP accumulation was also absent in intestines of L2-L3-stage larvae treated with stronger RNAi conditions (not shown; Methods). Top, confocal images, bottom, corresponding confocal/Normaski overlays. Insets show higher magnification images of embryos (uterus with embryos is bracketed in first column). Yellow arrowheads (top) and corresponding black arrowheads (bottom) bracket widths (small arrowheads) and lengths (large arrowheads) of intestines.
**(B)** The recycling defective mutant TGN-38(Y314A) reaches the plasma membrane in expanding *dab-1, rab-7 and vha-6(*RNAi) larval intestinal cells. Note lateral membrane broadening into a tubular network in the *vha-6(RNAi)* intestine (inset, arrows).
**(C)** Co-localization of SLCF-1 with RAB-5 and RME-1 in *rab-7(RNAi)* larval intestinal cells. Corresponding single color images to **Fig.8C, D.** Arrows point to GFP+/RFP+/DAPI- vesicles, identifying double labeled vesicles (as opposed to autofluorescent triple positive LROs). See **Fig.8C, D** for legend.
**(D)** The RFP::RME-1 and RFP::RAB-5 transgenic backgrounds do not affect SLCF-1::GFP displacement to cytoplasmic vesicles by *rab-7* RNAi in larval intestinal cells. Also note the significant increase of cytoplasmic SLCF-1+ vesicles in *rab-7(RNAi)* intestinal cells that demonstrates effectiveness of *rab-7* RNAi in these colocalization experiments and quantifies the vesicular SLCF-1 displacement during membrane expansion for this specific suppressor (see Methods for quantification).
Confocal images are shown throughout, full young adult intestines in **(A)**, 2 pairs of opposing larval intestinal cells in **(B, C)**. All data shown as mean $\pm$ SEM, *$p < 0.05$, **$p < 0.01$, ***$p < 0.001$. Scale bars, 5μm.

### Fig. S9

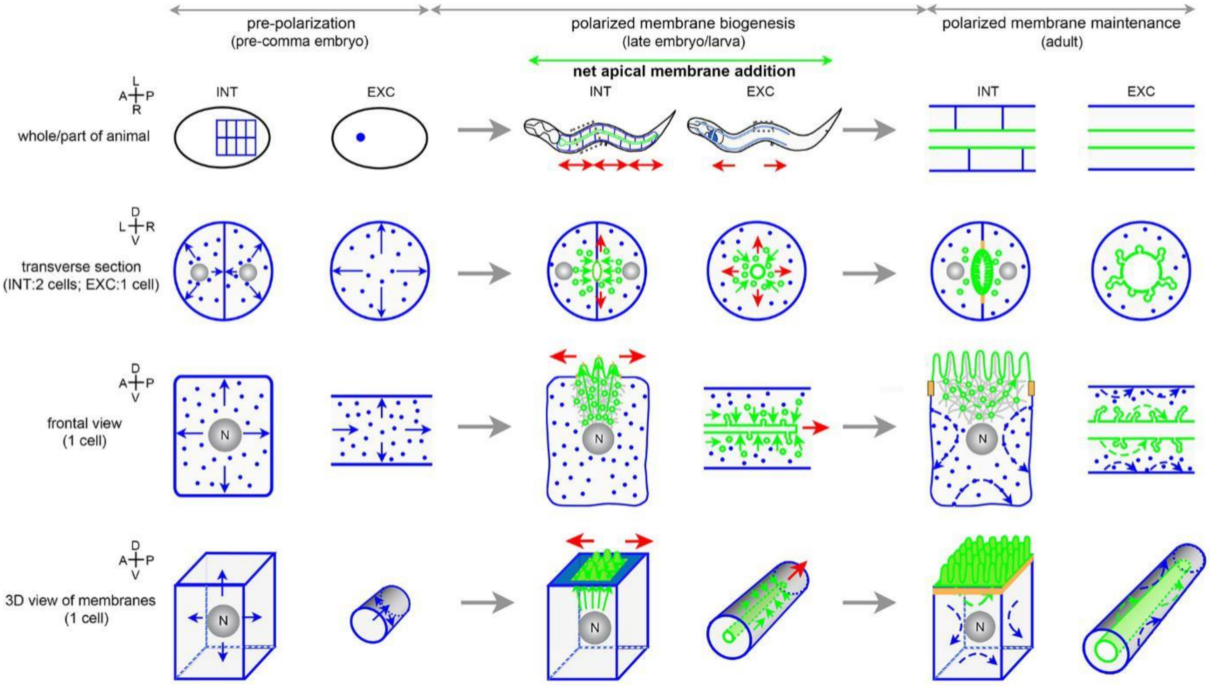
pre-polarization
(pre-comma embryo)
polarized membrane biogenesis
(late embryo/larva)
net apical membrane addition
polarized membrane maintenance
(adult)
INT
EXC
whole/part of animal
transverse section
(INT:2 cells; EXC:1 cell)
frontal view
(1 cell)
3D view of membranes
(1 cell)
N

**Figure S9 An apicobasal polarity model based on net apical membrane addition (related to Figure 9).**

The asymmetric addition of apical membrane that extends the inTRAcellular lumen in the excretory canal can serve as prototype for a mode of membrane polarization in the intestine, where its hypothesized inTERcellular addition between pairs of cells would position the apical domain of each cell and thereby define the membrane's polarity. Each of the three double columns shows intestinal (INT; left) and excretory canal (EXC; right) membrane biogenesis side by side. Directional apical membrane insertion (middle columns; red arrows) partitions the non-polarized membrane (left double columns) into apical and basolateral domains (right double columns).

Whole/parts of animals/tissues shown in upper row (boxed areas in middle column magnified in right column), different views of single cells beneath. Apical and basolateral (or non-polarized) endo-, plasma membranes green and blue, respectively (**Figs.1A, S1A** for intestinal and canal tubulogenesis). Trafficking routes/directions are simplified for clarity: only secretory routes (solid arrows) shown in left and middle double columns (only apical secretory routes in middle double columns); only recycling routes (dashed arrows) in right double columns. Indirect routes (via endosomes) and interfacing (e.g. transcytotic) routes are omitted (compare **Fig.6A**). Hypothesized transient dynamic cytoskeleton moving vesicles towards the nascent apical domain indicated in 3$^{rd}$ row (grey) and junctions (orange) in 3$^{rd}$ and 4$^{th}$ row. N: nucleus.

**Fig. S10**

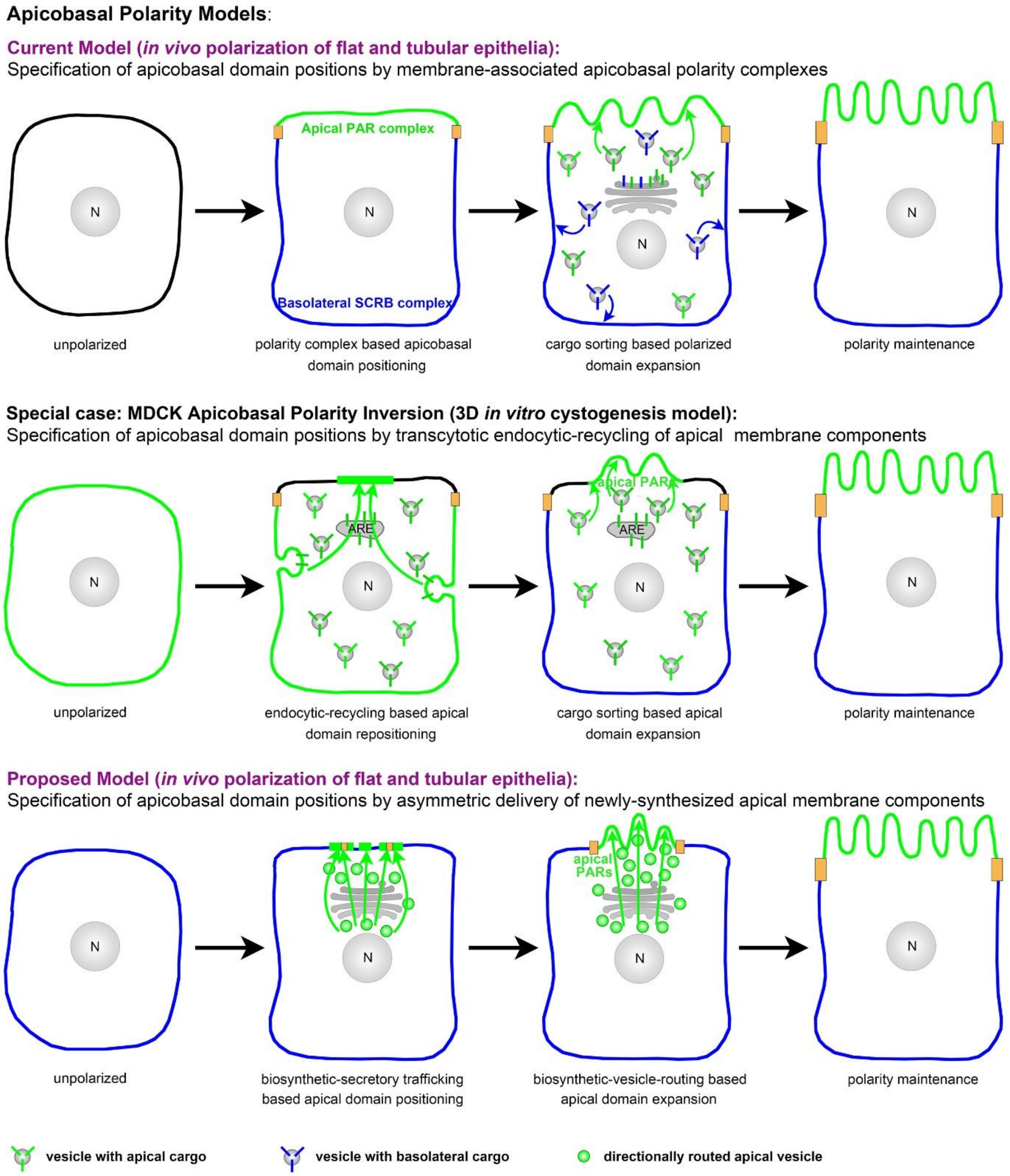
Apicobasal Polarity Models:
Current Model (in vivo polarization of flat and tubular epithelia):
Specification of apicobasal domain positions by membrane-associated apicobasal polarity complexes
N
Apical PAR complex
Basolateral SCRB complex
unpolarized
polarity complex based apicobasal domain positioning
cargo sorting based polarized domain expansion
polarity maintenance
Special case: MDCK Apicobasal Polarity Inversion (3D in vitro cystogenesis model):
Specification of apicobasal domain positions by transcytotic endocytic-recycling of apical membrane components
ARE
apical PARs
unpolarized
endocytic-recycling based apical domain repositioning
cargo sorting based apical domain expansion
polarity maintenance
Proposed Model (in vivo polarization of flat and tubular epithelia):
Specification of apicobasal domain positions by asymmetric delivery of newly-synthesized apical membrane components
apical PARs
unpolarized
biosynthetic-secretory trafficking based apical domain positioning
biosynthetic-vesicle-routing based apical domain expansion
polarity maintenance
vesicle with apical cargo
vesicle with basolateral cargo
directionally routed apical vesicle

**Figure S10 Apicobasal membrane polarization by domain composition change (current model), domain transcytosis (MDCK cystogenesis), and domain insertion (proposed model; related to Figure 9).**
Schematics of epithelial cells during *in vivo* membrane polarization (row 1 and 3) and *in vitro* membrane polarity inversion, initiated by an extracellular cue in Madin-Darby-Canine-Kidney/MDCK cells (change to 3D culture conditions; row 2; see text; (*19*)). See **Fig.9** legend for description of cartoons and details of rows 1 and 3 (current and proposed polarity models). The special case of MDCK cyst polarity inversion (row 2), requiring apical recycling routes, mirrors *C. elegans* intestinal polarity conversion, induced by interference with apical secretory routes that may traverse the recycling compartment (not shown; see text). Note that during MDCK cystogenesis the apical membrane shifts from the future basolateral to the new apical location via transcytosis. The apical recycling marker RAB-11/Rab11 is required for apical domain positioning during both MDCK cystogenesis and *C. elegans* intestinal tubulogenesis, suggesting that these different trafficking routes might converge in the vicinity of the nascent apical domain (e.g. at the ARE; see **Fig.9** inset). Endocytic-recycling routes required for apicobasal polarity conversion in the *C. elegans* intestine (see this study's suppressor analysis), on the other hand, might interface with the apical biosynthetic-secretory route (proposed model, row 3) as do basolateral endocytic routes that initiate MDCK cyst polarity inversion (not shown).

**Table S1. Vesicular trafficking genes for apical membrane biogenesis/lumen morphogenesis, identified in several *C. elegans* multi- and unicellular tubulogenesis screens (tier-1 genes; related to Figure 2).**

See Excel spreadsheet.

**Table S2. Classification of the identified tier-1 trafficking genes by trafficking route/function. Tier-1 trafficking genes previously identified in genetic screens on apical cargo delivery and secretion and tier-1 trafficking genes previously implicated in apical domain positioning (related to Figure 4).**

Classification of the identified tier-1 trafficking genes by trafficking route/function. Tier-1 trafficking genes previously identified in genetic screens on apical cargo delivery and secretion and tier-1 trafficking genes previously implicated in apical domain positioning (related to Figure 4).

| Gene | Classification of identified genes by trafficking route/function[1] | | | | Prior implication in apical cargo delivery, secretion and polarity | | |
|---|---|---|---|---|---|---|---|
| | I Biosynthetic/secretory pathway | II V-ATPase subunit components | III Endocytosis/ recycling/ degradation | IV Vesicular motors and associated | Apical transport (PEPT-1/ intestine, E-cadherin/ hypodermis; *C. elegans*; other) | Apical secretion (*Drosophila* tracheal tubes/*C. elegans* hypodermis) | Apical domain positioning (MDCK cysts/mouse/*C. elegans* intestine) |
| *sec-24.1* | + | | | | √ [4] | √ [7, 8] | |
| *sec-23* | + | | | | √ [4,5] | √ [7] | |
| *copz-1* | + | | | | √ [4] | | |
| *syx-5* | + | | | | √ [5] | √ [9] | |
| *ykt-6* | + | | | | | | |
| *syx-7*[2] | + | | | | | | |
| *sly-1* | + | | | | | | |
| *dab-1* | | | + | | | | |
| *apt-9* | + | | | | | | |
| *unc-11* | | | + | | | | |
| *rab-1* | + | | | | | √ [9] | |
| *rab-5* | | | + | | √ [4,5] | | √ [11] |
| *rab-6.2* | | | + | | | | |
| *rab-7* | | | + | | √ [5] | √ [9] | |
| *rab-8* | + | | | | | √ [9] | √ [12,13,14] |
| *rab-11* | | | + | | √ [4] | √ [9] | √ [13,14,15] |
| *rab-21* | | | + | | | | |
| *arf-1.1* | + | | | | | | |
| *arf-6* | | | + | | | | |
| C52B11.5 | | | + | | | | |
| *sec-12* | + | | | | √ [5] | | |
| *ifta-2* | | | + | | | | |
| *ggtb-1* | + | | | | | | |
| *F38A1.8* | + | | | | √ [5] | | |
| *drp-1* | | | + | | | | |
| *dyn-1* | | | + | | √ [4] | | |
| *vps-32.1* | | | + | | √ [4,5] | √ [9] | |
| *vps-33.1* | | | + | | √ [5] | | |
| *vha-1* | | + | | | √ [4,5] | | √ [16] |
| *vha-2* | | + | | | √ [4,5] | | |
| *vha-4* | | + | | | √ [4,5] | | √ [16] |
| *vha-5* | | + | | | √ [5] | √ [10] | |
| *vha-6* | | + | | | √ [4,5] | | |
| *vha-8* | | + | | | √ [4,5] | | |
| *vha-9* | | + | | | √ [4,5] | √ [9] | |
| *vha-10* | | + | | | √ [5] | | |
| *vha-11* | | + | | | | | |
| *vha-12* | | + | | | √ [4,5] | | |
| *vha-15* | | + | | | √ [4,5] | | |
| *vha-18* | | + | | | | | |
| *klp-7* | | | | + | | | |
| *klp-10* | | | | + | | | |
| *klp-16* | | | | + | √ [6] | | |
| *vab-8* | | | | + | | | |
| *spdl-1* | | | | + | | | |
| *che-3* | | | | + | | | |
| *mtm-6* | | | + | | | | |
| *inpp-1* | | | + | | | | |
| *maa-1* | | | + | | | | |
| *eps-8*[3] | | | + | | | | √ [16] |

[1]Classified according to demonstrated or predicted predominant trafficking function or route (compare Table S1)
[2]Classified as secretory pathway component based on strongest homology to yeast SSO2
[3]function in apical membrane microdomain biogenesis, possibly unrelated to trafficking; ref *(105)*
[4]ref *(100)*
[5]ref *(101)*
[6]ref *(34)*
[7]ref *(81)*
[8]ref *(102)*
[9]ref *(103)*
[10]ref *(36)*
[11]ref *(33)*
[12]ref *(32)*
[13]ref *(19)*
[14]ref *(104)*
[15]ref *(39)*
[16]ref *(40)*

**Table S3. Timecourse of polarity conversion in double *let-767(-/-)* mutant/suppressor knockdowns (related to Figure 6B, C).**

| Table S3 Timecourse of polarity conversion in double *let-767(-/-)* mutant/suppressor knockdowns (related to Figure 6B, C) | | | | | | | | | | | | | | | | | | | | |
|---|---|---|---|---|---|---|---|---|---|---|---|---|---|---|---|---|---|---|---|---|
| | day3 | | | | day4 | | | | day5 | | | | day6 | | | | day7 | | | |
| | WT (Dpy)[1] | BL[2] | BL&EL[3] | N[4] | WT (Dpy) | BL | BL&EL | N | WT (dpy) | BL | BL&EL | N | WT (Dpy) | BL | BL&E | N | WT (Dpy) | BL | BL&E | N |
| MOCK RNAi | 32% | 46% | 21% | 376 | 5% | 36% | 59% | 359 | 2% | 5% | 93% | 394 | 1% | 2% | 97% | 371 | 0 | 0 | 100% | 318 |
| *vha-6* RNAi | 96% | 4% | 0% | 235 | 88% | 11% | 1% | 220 | 64% | 17% | 18% | 243 | 40% | 12% | 48% | 203 | 20% | 11% | 68% | 214 |
| *rab-7* RNAi | 41% | 54% | 6% | 238 | 14% | 74% | 12% | 250 | 7% | 63% | 31% | 232 | 5% | 43% | 53% | 211 | 1% | 35% | 64% | 215 |
| *dab-1* RNAi | 63% | 34% | 3% | 196 | 35% | 46% | 19% | 212 | 16% | 26% | 58% | 220 | 12% | 15% | 73% | 214 | 5% | 8% | 84% | 191 |
| *rab-6.2* RNAi | 31% | 53% | 16% | 239 | 12% | 43% | 44% | 224 | 3% | 25% | 72% | 224 | 0% | 9% | 91% | 209 | 0% | 2% | 97% | 199 |
| *arf-6* RNAi | 37% | 56% | 7% | 185 | 17% | 52% | 31% | 224 | 9% | 24% | 67% | 204 | 4% | 10% | 86% | 197 | 2% | 5% | 92% | 165 |

Note: All RNAi experiments were carried out in *a let-767(s2819); sDp3* mutant background supplemented with ERM-1::GFP (see text). Standard RNAi conditions were used, with evaluation of progeny (Methods). Day of RNAi set-up is refered to as day 0. On day 3 Dpy-marked *let-767* (s2819)-/- L1-larvae were transferred to a new RNAi plate and their polarity phenotypes were recorded over the next 5 days. Independent experiments were performed 3 times for each gene.

[1]WT = wild-type apical ERM-1::GFP

[2]BL = basolateral ERM-1::GFP displacement

[3]BL&EL = basolateral ERM-1::GFP displacement and ectopic lumen formation

[4] Total number of animals counted; Dpy-marked L1 larvae were picked to separate plates for counting

**Table S4. Reagent and Strain list.**

| **Table S4. Reagent and Strain list** | | |
|---|---|---|
| **REAGENT or RESOURCE** | **SOURCE** | **IDENTIFIER** |
| **Antibodies** | | |
| anti-IFB-2 (mouse) | Developmental Studies Hybridoma Bank (DSHB) | MH33 |
| FITC (anti-mouse) | Sigma | Cat#F9006 |
| **Bacterial and Virus Strains** | | |
| E. coli: OP50: E. coli B, uracil auxotroph | Caenorhabditis Genetics Center (CGC) | WB Strain: OP50 |
| E. coli: HT115 (DE3): F-, mcrA, mcrB, IN(rrnD-rrnE)1, rnc14::Tn10(DE3 lysogen: lavUV5 promoter -T7 polymerase) (IPTG-inducible T7 polymerase) (RNAse III minus). | CGC | WB Strain: HT115(DE3) |
| E. coli: DsRed HT115 | Zhang et al., 2011 | N/A |
| E. coli: Lethal RNAi library | Gary Ruvkun | N/A |
| E. coli: Ahringer genome-wide RNAi feeding library | Julie Ahringer | N/A |
| **Chemicals, Peptides, and Recombinant Proteins** | | |
| BODIPY-Cer | Invitrogen | Cat#B34400 |
| **Strains** | | |
| N2 (Bristol) | CGC | WB Strain: N2 |
| *let-767(s2819) ncl-1(e1865) dpy-17(e164) unc-32(e189)III; sDp3(III;f)* | David L. Baillie | BC4849 |
| *slcf-1(tm2258); Exfs254[slcf-1::gfp]* | Florence Solari | FS254 |
| *pha-1(e2123ts)III, him-5(e1490)V, rnyEx060 [pELA2 (vha-6p::vha-6::mCherry); myo-3p::pHluorin; pha-1(+)]* | Keith Nehrke | KWN117 |
| *pha-1(e2123) III,rnyEx133[opt-2(aa1-412)::GFP) + pha-1(+)]* | CGC | WB strain: KWN246 |
| *dab-1(gk291)II; feEx43[dab-1::gfp rol-6(su1006)]* | Jonathan Pettitt | PE291 |
| *vhIs34 [vha-6p::GFP::rab-7Q68L, Cbunc-119 ]* | Christian E. Rocheleau | QR124 |
| *pwIs23[vit-2::gfp]* | Barth D. Grant | RT130 |
| *unc-119(ed3)III; pwIs69 [vha-6p::gfp::rab-11, Cbunc-119(+)]* | CGC/Barth D. Grant | WB strain: RT311 |
| *unc-119(ed3); pwIs112[vha-6p::hTAC::GFP, Cbunc-119(+)]* | Barth D. Grant | RT393 |
| *unc-119(ed3)III; pwIs170[vha-6p::gfp::rab-7, Cbunc-119(+)]* | CGC/Barth D. Grant | WB strain: RT476 |
| *unc-119(ed3)III; pwIs206[pvha-6::gfp::rab-10 Cbunc-119(+)]* | CGC/Barth D. Grant | WB strain: RT525 |
| *pwIs846[vha-6p::RFP::RAB-5]* | Barth D. Grant | RT2293 |

| | | |
|---|---|---|
| *pwIs852[vha-6p::RFP::RME-1]* | Barth D. Grant | RT2299 |
| *tgn38p::tgn-38(Y314A)::gfp* | Barth D. Grant/Bai&Grant, 2015 | RT3292 |
| *jcEx44 [ajm-1::GFP+ rol-6(su1006)]* | CGC/Jeff Hardin | WB strain: SU159 |
| *rab-7(ok511)/mIn1 [mIs14 dpy-10(e128)] II* | CGC | WB strain: VC308 |
| *dab-1(gk291) II* | CGC | WB strain: VC616 |
| *vha-6(ok1825)/mIn1 [mIs14 dpy-10(e128)] II* | CGC | WB strain: VC1336 |
| *fgEx12[(act-5p::act-5::gfp);unc-119(+);unc-119(ed3)III]* | Verena Gobel | VJ268 |
| *fgIs100[erm-1p::erm-1::gfp, rol-6p::rol-6(su1006)]* | Verena Gobel | VJ610 |
| *rrf-3(pk1426)II; fgEx13[erm-1p::erm-1::gfp, rol-6p::rol-6(su1006)]* | Verena Gobel | VJ462 |
| *let-767(s2819) ncl-1(e1865) dpy-17(e164) unc-32(e189)III; sDp3(III;f); fgEx13[erm-1p::erm-1::gfp;rol-6p::rol-6(su1006)]* | Verena Gobel | VJ501 |
| *fgEx17[aqp-1p::aqp-1::gfp;rol-6p::rol-6(su1006)]* | Verena Gobel | VJ507 |
| *fgIs2(erm-1p::erm-1, rol-6p::rol-6(su1006); fgEx20(vha-1p::gfp, vha-2p::gfp, unc-119+); unc-119(ed3)III* | Verena Gobel | VJ536 |
| *fgIs2[erm-1p::erm-1, rol-6p::rol-6(su1006)] ;pha-1(e2123ts)III; rnyEx188 [pFH6-K12.2 (Psulp-5::GFP); pCL1 (pha-1+)]* | Verena Gobel | VJ557 |
| *pwIs846[vha-6p::RFP::RAB-5]; slcf-1(tm2258); Exfs254[slcf-1::gfp]* | this study | VJ561 |
| *pwIs852[vha-6p::RFP::RME-1]; slcf-1(tm2258); Exfs254[slcf-1::gfp]* | this study | VJ562 |
| *par-6(it319[par-6::GFP]) I* | CGC/Kenneth Kemphues | KK1248 |
| *pkc-3(it309[GFP::pkc-3]) II* | CGC/Kenneth Kemphues | KK1228 |

**Movie S1. 3D view of apical junction integrity in *copz-1(RNAi)* larval intestine (related to Figure 4, S4).**
